# A suite of activity-based probes to dissect the KLK activome in drug-resistant prostate cancer

**DOI:** 10.1101/2021.04.15.439906

**Authors:** Scott Lovell, Leran Zhang, Thomas Kryza, Anna Neodo, Nathalie Bock, Elizabeth D. Williams, Elisabeth Engelsberger, Congyi Xu, Alexander T. Bakker, Elena De Vita, Maria Maneiro, Reiko J. Tanaka, Charlotte L. Bevan, Judith A. Clements, Edward W. Tate

**Author notes:** Correspondence should be addressed to E.W.T.

## Abstract

Kallikrein-related peptidases (KLKs) are a family of secreted serine proteases, which form a network – the KLK activome – with an important role in proteolysis and signaling. In prostate cancer (PCa), increased KLK activity promotes tumor growth and metastasis through multiple biochemical pathways, and specific quantification and tracking of changes in the KLK activome could contribute to validation of KLKs as potential drug targets. Herein we report a technology platform based on novel activity-based probes (ABPs) and inhibitors with unprecedented potency and selectivity enabling simultaneous orthogonal analysis of KLK2, KLK3 and KLK14 activity in hormone-responsive PCa cell lines and tumor homogenates. Using selective inhibitors and multiplexed fluorescent activity-based protein profiling (ABPP) we dissect the KLK activome in PCa cells and show that increased KLK14 activity leads to a migratory phenotype. Furthermore, using biotinylated ABPs we show that active KLK molecules are secreted into the bone microenvironment by PCa cells following stimulation by osteoblasts suggesting KLK-mediated signaling mechanisms could contribute to PCa metastasis to bone. Together our findings show that ABPP is a powerful approach to dissect dysregulation of the KLK activome as a promising and previously underappreciated therapeutic target in advanced PCa.

## Introduction

Prostate cancer (PCa) is the most frequently diagnosed cancer among men in industrialized nations and remains a leading cause of cancer-related deaths.^1^ Localized malignancies are treated with surgery and radiation therapy and the 5-year survival rate of patients is close to 100%; however, post-operative recurrence often progresses to advanced PCa. Initial treatment for advanced tumors exploits the dependence of PCa cells on androgens by combining androgen-deprivation therapy (ADT) with direct targeting of the androgen receptor (AR),^2,3^ but many patients stop responding and develop castrate-resistant prostate cancer (CRPC). In CRPC, PCa cells evolve resistance to androgen-targeting therapies by restoring AR signaling through diverse mechanisms, and the majority of patients present with bone metastases with increased risk of morbidity and mortality due to alterations in skeletal integrity.^4^ Mapping critical pathways involved in establishing CRPC may enable identification of novel therapeutic targets which can ultimately reduce disease recurrence.^5^

AR is a transcription factor that dimerizes and translocates to the nucleus upon binding of androgens such as dihydrotestosterone (DHT), where it induces expression of a variety of genes important for PCa cell proliferation and survival (**Fig. 1A**).^6^ Among these genes are specific serine proteases from the fifteen-member human kallikrein-related peptidase (KLK) family, which have versatile and crucial roles in extracellular proteolysis and signaling.^7,8^ For example, increased expression and subsequent leakage of KLK2 and KLK3, also known as Prostate-Specific Antigen (PSA), into the vasculature are used in PCa diagnosis and progression monitoring.^9,10^ KLK2 and KLK3 also have functional roles in PCa, and contribute to disease progression. KLK2 can activate protease-activated receptors (PARs) on the surface of neighboring fibroblasts, which in turn release cytokines that stimulate PCa cell proliferation,^11^ whilst in the bone microenvironment KLK3 promotes osteoprogenitor cell proliferation and osteoblast differentiation, and establishment of bone metastases.^12,13^ More recently, KLK14 has also been implicated in CRPC development; whilst KLK14 is normally suppressed by AR signaling (**Fig. 1A**), treatment with AR-targeted drugs may increase KLK14 expression to promote PCa cell migration and bone matrix colonization.^14,15^

**Figure 1:**
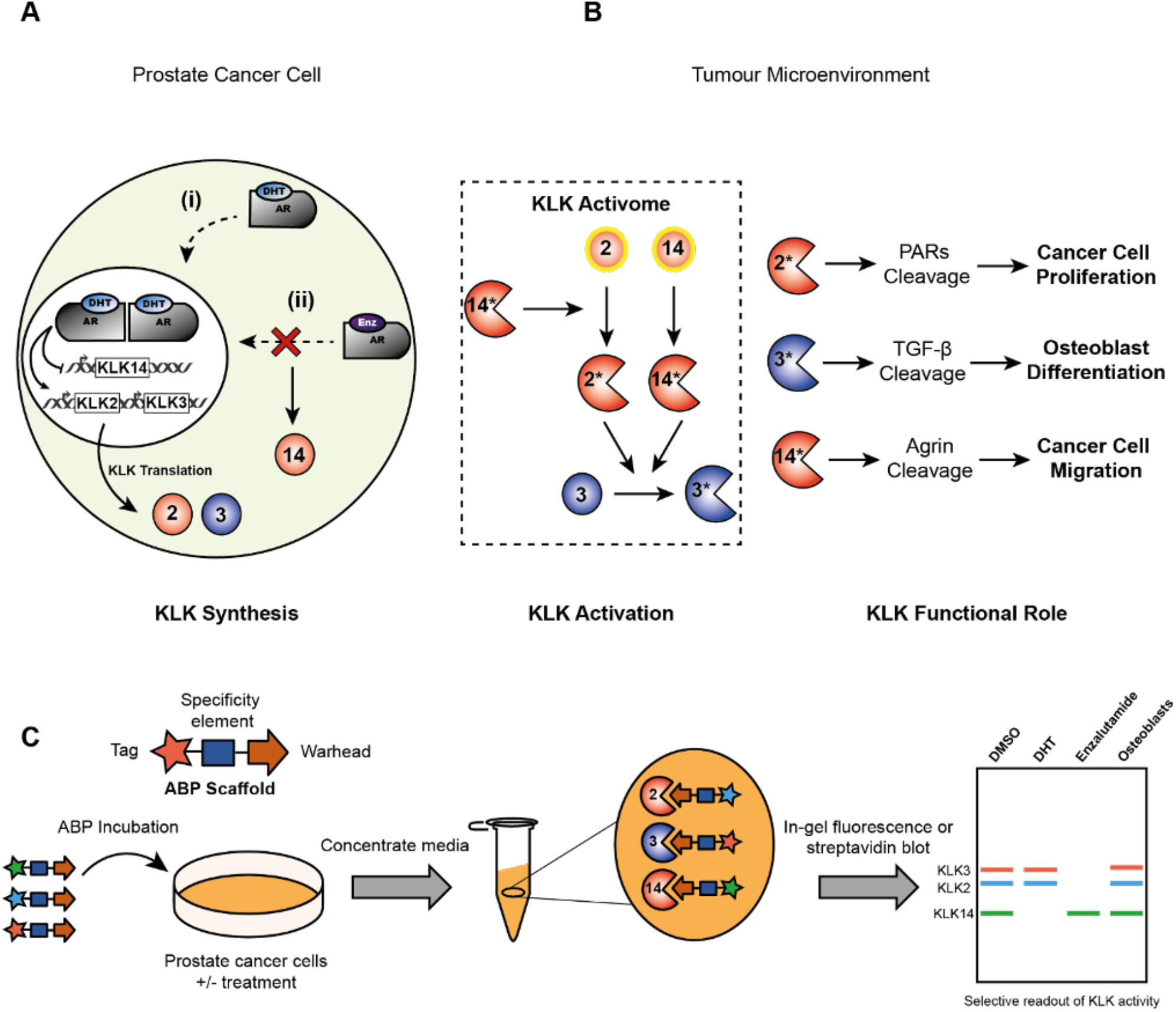
The human kallikrein-related peptidases in prostate cancer (**A**) **KLK Synthesis**: KLK2 and KLK3 expression is induced by the transcriptional activity of the AR. Conversely, KLK14 expression is negatively regulated by AR signaling and is upregulated upon treatment with enzalutamide, a competitive AR inhibitor; (**B**) **KLK Activation**: KLKs are secreted to the tumor microenvironment as pro-enzymes and mostly require trypsin-like proteolytic processing to become catalytically active. In prostate cancer a regulatory cascade termed the ‘KLK activome’ has been proposed whereby the trypsin-like proteases KLK2 and KLK14 auto-activate prior to cross-activating other pro-KLK molecules. Red = trypsin-like activity, blue = chymotrypsin-like activity, yellow outline = auto-activation; **KLK Functional Roles**: In addition to degrading components of the ECM, active KLK molecules are involved in key stages of PCa progression including cell proliferation, cell migration and bone metastasis (**C**) A schematic representation of the proposed ABPP workflow to enable quantification of KLK activity in PCa cell conditioned media and tumor homogenate. ABPs are incubated with PCa cells, conditioned media is collected and concentrated using a MWCO spin filter, protein samples are subjected to SDS-PAGE and labeling is then visualized by in-gel fluorescence or streptavidin blotting.

Despite recent progress in determining the pathophysiological roles of individual KLKs in PCa, the potential of KLK inhibitors for therapeutic intervention remains to be determined. Once secreted by PCa cells, KLKs do not work in isolation but as components of a complex network called the KLK activome (**Fig. 1B**) that is tightly regulated by proteolytic KLK autoactivation, cross-activation or deactivation, and inhibition by endogenous serine protease inhibitors.^16–19^ A method to quantify the activity levels of each KLK directly in a complex biological system would enable the dynamic response of the KLK activome to be determined in relevant PCa models, drug-resistant tumors, and during drug/hormone-mediated perturbations, and may lead to validation of specific active KLKs as drug targets or novel biomarkers.

Here we present a technology platform which enables simultaneous orthogonal readout of the activity of KLK2, KLK3 and KLK14 in androgen-responsive PCa cell lines and patient-derived xenografts (PDX), based on a chemical toolbox of first-in-class selective activity-based probes (ABPs) and inhibitors for activity-based protein profiling (ABPP, **Fig. 1C**). We use this platform to show that active KLK molecules are secreted into the bone microenvironment by PCa cells following stimulation by osteoblasts, supporting a double paracrine signaling mechanism that may contribute to the establishment of bone metastases mediated by KLK activity. Furthermore, using selective inhibitors and multiplexed fluorescent ABPP we dissect the KLK activome directly in PCa cells and show that KLK14 drives PCa cell migration, a key step in the formation of distant metastases. Together these findings demonstrate that ABPP can provide a unique window on KLK activity and inhibition in PCa and suggest that KLK-activome dysregulation is a promising and previously underappreciated therapeutic target in CRPC.

## Results

### Development of a selective inhibitor and activity-based probe for KLK3

Proof-of-concept for the first selective ABP targeting the PCa KLK activome started with KLK3, which exhibits chymotrypsin-like specificity, in contrast to KLK2 and KLK14 which are trypsin-like proteases.^20^ An optimized tetrapeptide designed to occupy the KLK S1-S4 subsites was envisaged for the ABP specificity element, grafted onto a peptidyl-diphenyl phosphonate (DPP) warhead which reacts specifically and exclusively with the Ser195 residue in the KLK3 active site.^21^ DPP **1** with a tyrosine-mimicking phenol in the first position (P1) was previously identified as a modestly potent inhibitor of KLK3,^22^ while peptidyl-boronic acid **2** is a potent and selective KLK3 inhibitor (**Fig. 2A**).^23^ Both DPP and boronic acid inhibitors covalently modify Ser195, thereby mimicking the tetrahedral intermediate during amidolysis, however boronic acids form reversible covalent complexes and are generally less effective as ABPs. We therefore replaced the boronic acid in **2** with the DPP moiety in **1** and capped the N-terminus with a tetramethyl rhodamine (TAMRA) fluorophore to afford **3**, which served as a prototype ABP for KLK3. A mixed solid-phase and solution-phase approach was used for the synthesis of peptidyl-DPP compounds, as described previously (**Fig. S1** and **S2**).^24^ KLKs are typically activated after their secretion from the cell, so we first examined the labelling profile of **3** in conditioned media (CM) obtained from LNCaP cells treated with androgen (10 nM DHT). LNCaP is an androgen-responsive human prostate adenocarcinoma cell line widely used to model key features of clinical disease, including secretion of KLK2 and KLK3 following androgen stimulation.^8^ LNCaP CM was treated with different concentrations of **3** for 1 h and proteins separated by SDS-PAGE. In-gel fluorescence revealed a single major target at ca. 32 KDa, labeled in a concentration-dependent manner (**Fig. 2B**), and selective covalent modification of active KLK3 was confirmed by immunoprecipitation (**Fig. S3**).

**Figure 2:**
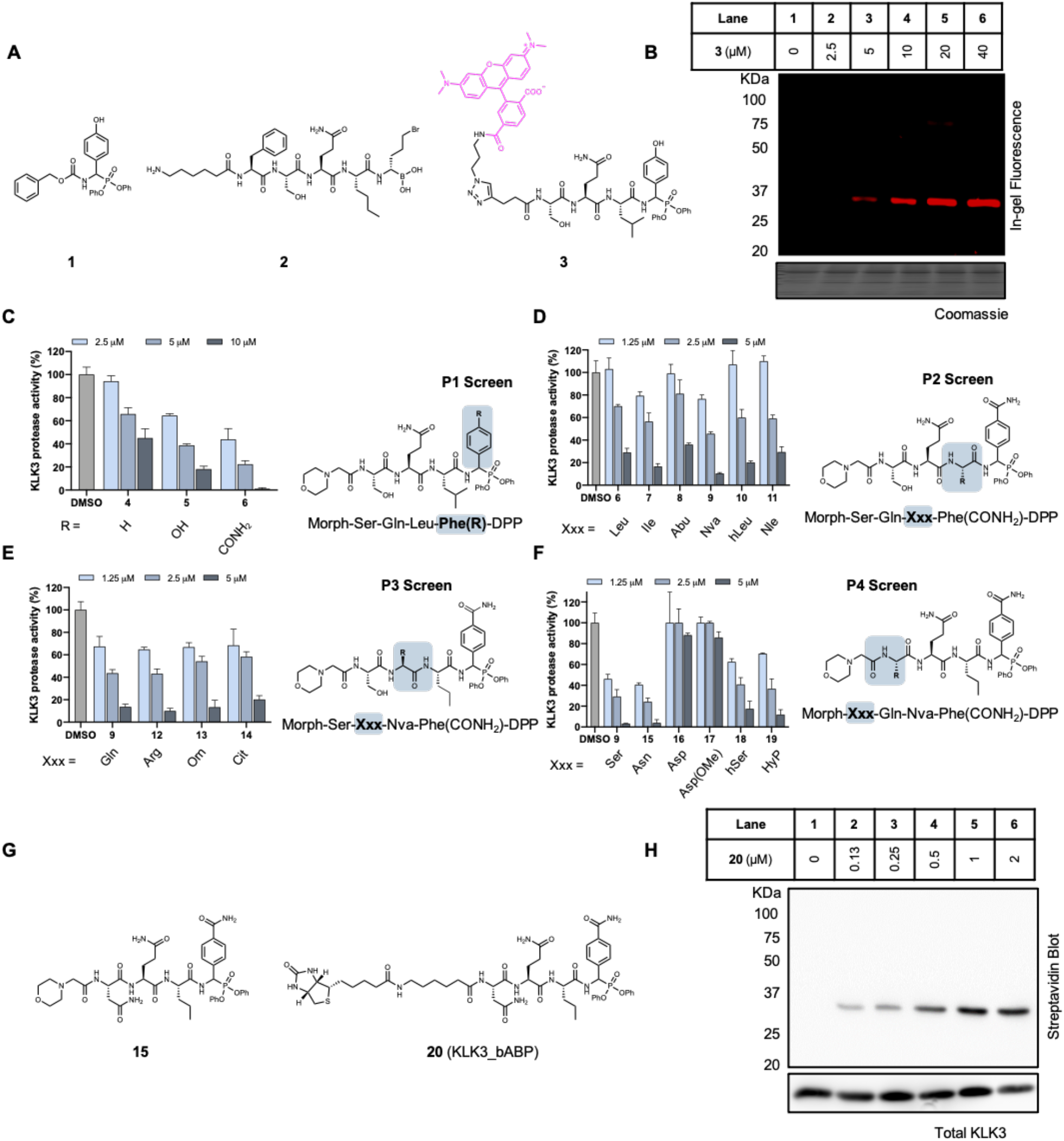
Development and optimization of a KLK3 ABP using competitive ABPP (**A**) Structures of reported KLK3 inhibitors **1** and **2** and first-generation fluorescent ABP **3** (**B**) In-gel fluorescence shows selective, dose-dependent labeling of KLK3 after treatment of LNCaP conditioned media with **3** for 1 h (**C-F**) Inhibition of KLK3 protease activity by morpholine-capped peptidyl-DPP analogues. LNCaP conditioned media was first incubated with the analogue for 1 h and then with **3** for 1 h. Protein samples were separated by SDS-PAGE and residual KLK3 activity was calculated by densitometry. See Fig. S4 and S5 for fluorescent gels (**G**) Chemical structures of optimised KLK3 inhibitor **15** and ABP **20** (**H**) Streptavidin blotting shows potent and selective labeling of KLK3 after treatment of LNCaP conditioned media with **20** for 1 h.

Despite exquisite selectivity, maximal KLK3 labeling was achieved only after treatment with 20 μM **3** for 1 h. We therefore optimized probe potency by systematically modifying each amino acid sidechain in the P1-P4 positions of a morpholine-capped peptidyl-DPP inhibitor scaffold, and determined the potency of each analogue using competitive-ABPP against **3**.^25^ LNCaP CM was first incubated with a peptidyl-DPP analogue for 1 h, followed by 1 h treatment with 20 μM **3** to assess the degree of KLK3 inhibition by densitometry (**Fig. S4** and **S5** for fluorescent gels and **Fig. S6** for structures of each morpholine-capped peptidyl-DPP analogue). Out of 16 analogues tested, compound **15** was identified as a potent KLK3 inhibitor (**Fig. 2C-G**). Despite its preference for Tyr in P1, KLK3 can also process substrates with P1 Gln,^26^ and a ‘hybrid’ benzamide sidechain in P1 resulted in a significant increase in potency (**Fig. 2C**). The P2-P4 specificity of KLK3 has been mapped in detail previously and only minor changes were required to further optimize potency, including substitution of P2 Leu for norvaline (Nva), and P4 Ser for Asn.^27^ Second generation biotinylated **20** and fluorescent **21** ABPs (**Fig. 2G** and **S7**) based on this scaffold retained exquisite selectivity for KLK3 in LNCaP CM but in contrast to first-generation ABP **3** achieved maximum labeling at only 1 μM and readily detectable labeling down to 130 nM **(Fig. 2H and S7)**. Importantly, **20** and **21** showed no inhibition of KLK2 and KLK14 even at concentrations as high as 50 μM (**Fig. S8**). Consequently, these probes are the most selective covalent inhibitors of KLK3 reported to date.

### Development of selective inhibitors and activity-based probes for KLK2 and KLK14

Both KLK2 and KLK14 display a strong preference for Arg in P1,^20^ and we anticipated that development of selective ABPs would require optimization of the P2-P4 positions to differentiate between unique preferences of the S2-S4 subsites. We generated a positional scanning substrate library derived from 19 natural amino acids (excluding Cys) and 86 structurally diverse non-natural amino acids to provide a detailed analysis of active site preferences at each protease subsite.^28^ Three sub-libraries consisting of 105 mixtures of 361 fluorogenic peptidyl coumarins were generated and screened against KLK2 and KLK14 (**Fig. 3A**; see **Fig. S9** and **S10** for library synthesis and structures of non-natural amino acids), and scatter plots of relative initial rates of hydrolysis for each amino acid revealed specificity preferences at the P2, P3 and P4 positions (**Fig. 3B-D** and **Fig. S11-S14**). Despite a general preference for aromatic residues at P2, KLK2 processed substrates with a P2 cyclohexyl alanine (Cha) at the highest rate, whereas KLK14 demonstrated dual specificity at P2, processing both aliphatic residues such as aminobutyric acid (Abu) and aromatic residues such as benzyl histidine (His(Bzl)) (**Fig. 3B**). Both proteases favored basic amino acids at P3, with KLK14 preferring Lys while KLK2 preferred the shorter chain diaminobutyric acid (Dab) (**Fig. 3C**). At P4, KLK2 processed aliphatic (e.g., Abu) and aromatic residues with a flexible linker such as benzyl-serine (Ser(Bzl)), and KLK14 preferred medium and large aromatic residues such as 4-bromophenylalanine (Phe(4-Br)) and benzothiazol-2-yl alanine (Ala(Bth)) (**Fig. 3D**).

**Figure 3:**
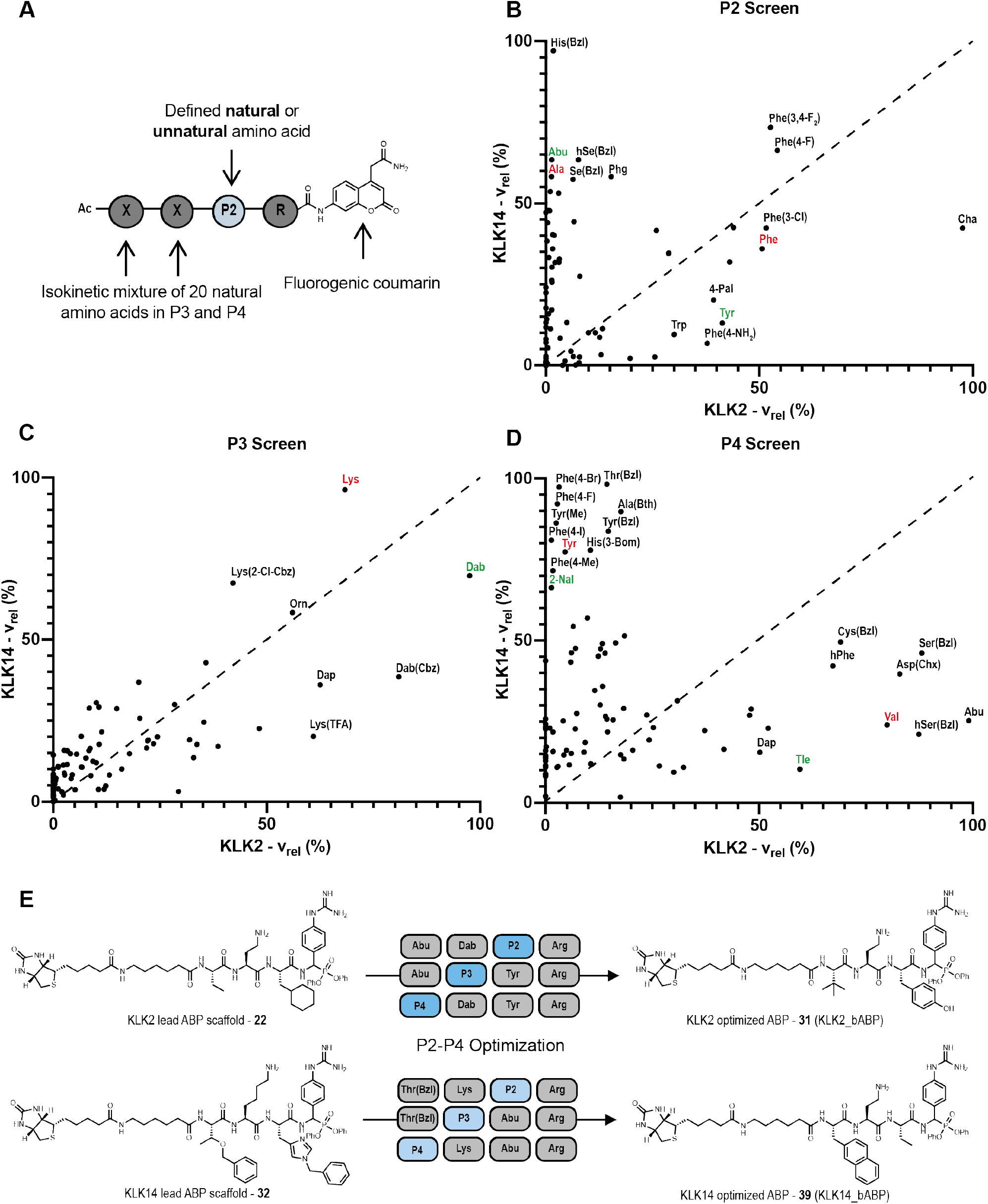
Comparative analysis of the S1-S4 subsite preferences of KLK2 and KLK14 using a positional scanning library approach (**A**) Structure of the peptidyl-coumarin scaffold used, exemplified with the P2 sub-library. P2 = any defined natural or unnatural amino acid. X = isokinetic mixture of 19 natural amino acids, excluding cysteine (**B-D**) Scatter plots showing relative initial rates of hydrolysis of each amino acid for KLK2 (x-axis) and KLK14 (y-axis). The optimal amino acid residue is set to 100 %. Amino acids that are close to the dotted diagonal line are equally tolerated by both KLKs. The preferred natural amino acid for each protease is shown in red and the amino acid that was used in the final optimised ABP is shown in green. The position of each amino acid represents the mean value of three replicates. See Fig. S11-S14 for complete data (**E**) Structures of KLK2 and KLK14 ABPs before and after P2-P4 optimization.

Based on these data, we hypothesized that the divergent specificity preferences of KLK2 and KLK14 could be exploited to develop selective chemical probes. The preferred amino acid for each sub-site was incorporated into a peptidyl-DPP scaffold to afford prototype probes **22** and **32** directed toward KLK2 and KLK14, respectively (**Fig. 3E**), using a phenylguanidine Arg mimic in P1 to ease synthesis.^29^ Kinetic analyses (**Fig. S15**) revealed that although **22** is a potent inhibitor of KLK2 (k_inact_/K_i_ = 3274 ± 89 M^−1^s^−1^) it has significant cross-reactivity with KLK14 (k_inact_/K_i_ = 511 ± 18 M^−1^s^−1^), and we therefore further optimized the KLK2 inhibitor scaffold by systematically altering P2, P3 and P4 residues with other hits from the substrate library screen. From nine peptidyl-DPP analogues, compound **31** was most optimal (**Table 1**, **Fig. S16**) with similar potency towards KLK2 (k_inact_/K_i_ = 3076 ± 87 M^−1^s^−1^) delivered by a P2 Tyr substitution, and >30-fold selectivity over KLK14 (k_inact_/K_i_ = 94 ± 7 M^−1^s^−1^) thanks to increased steric bulk at P4 (Tle in place of Abu). We took a similar approach to optimize compound **32** against KLK14 (k_inact_/K_i_ = 523 ± 46 M^−1^s^−1^), identifying compound **39** (**Table 1**, **Fig. S17**) with 80-fold higher potency towards KLK14 (44474 ± 1976 M^−1^s^−1^) and 220-fold selectivity over KLK2 (k_inact_/K_i_ = 204 ± 13 M^−1^s^−1^).

**Table 1.**
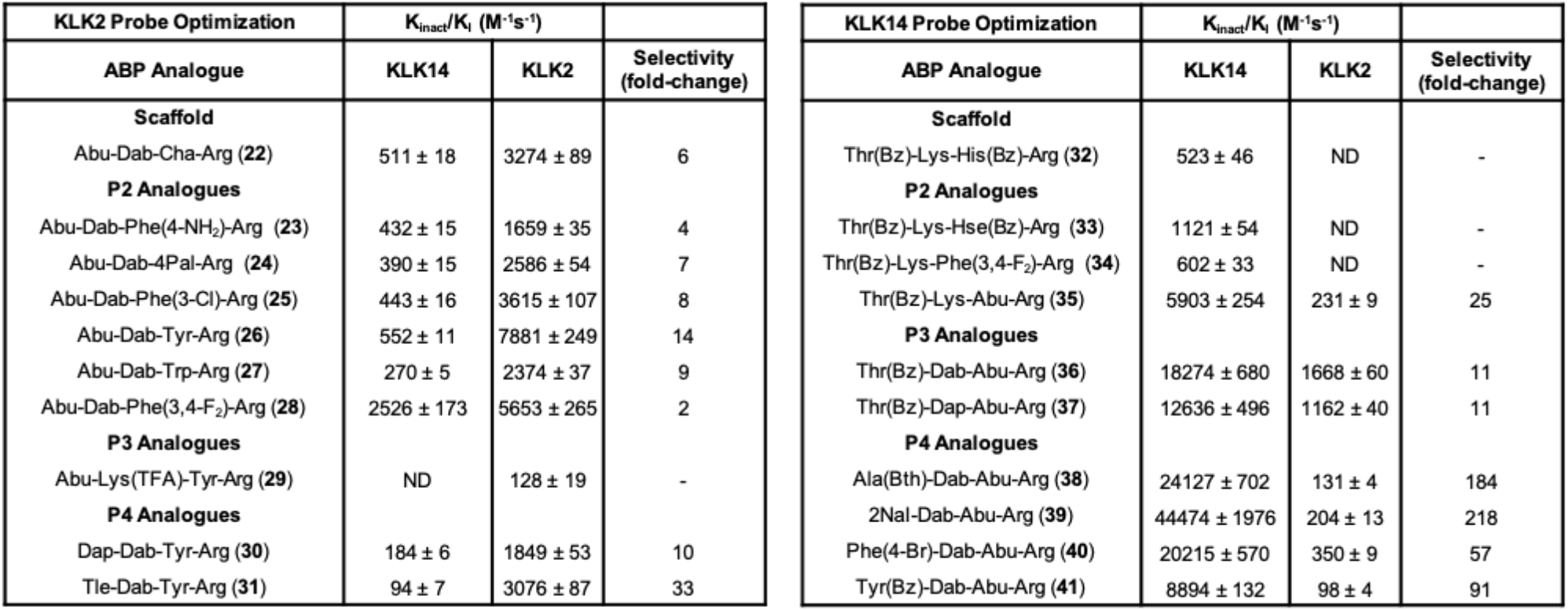
k_inact_/K_I_ values of peptidyl-DPP analogues for KLK2 (left) and KLK14 (right). Each data point is a mean value ± SEM (N=3).

To gain further insights into site preference we performed molecular modelling studies using previously determined KLK2 and KLK3 crystal structures^31,32^ in the Molecular Operating Environment (MOE) software package^33^ to propose binding determinants of KLK2_bABP and KLK3_bABP. For the purpose of modelling the specificity elements of each probe class, the biotin in KLK2_bABP was replaced with an acetyl group, and morpholino-capped derivative **15** was used to model KLK3_bABP. We found that KLK2_bABP and KLK3_bABP likely bind to their cognate KLK active sites in a conventional extended conformation, with the amino acid side chains of P1-P4 occupying the S1-S4 pockets (**Fig. S18A and S18B**). In both poses the phosphonate warhead is located in close proximity to the catalytic serine, with the P=O moiety establishing an H-bonding interaction with the NH group on the backbone of Gly193, which would activate the phosphonate for nucleophilic attack (**Fig. S18C and S18D**). Our modelling highlighted the differing properties of the S1 and S4 pockets of KLK2 and KLK3 as key determinants of probe specificity. For example, the P1 phenylguanidine of KLK2_bABP forms a salt bridge with the carboxylate group of Asp-189 at the bottom of the S1 pocket of KLK2. In contrast, for KLK3 the residue at the bottom of the S1 pocket is serine, which is also flanked by the polar side chains of Ser227 and Thr190. Consequently, the KLK3 S1 pocket is polar at the bottom and hydrophobic on the sides and has a preference for medium hydrophobic side chains with polar neutral head groups. In agreement with this, the P1 benzamide group of KLK3_bABP forms key hydrogen bonds with Ser227 and Thr190. Furthermore, our model suggests that the P4 Asn in KLK3_bABP forms a hydrogen bond with Gln174, which is positioned in the small, polar S4 pocket of KLK3. In contrast, the bulky P4 Tle residue of KLK2_bABP is accommodated by the S4 pocket of KLK2, which has significant hydrophobic character imparted from several residues of the kallikrein loop.^34^ A crystal structure for KLK14 is yet to be solved but a homology model by de Veer *et al.* provides a potential explanation for the selectivity of our KLK2 and KLK14 probes.^35^ The S2 pocket of KLK14 is narrow owing to the flanking sidechain of His99 and thus small aliphatic residues, such as the P2 Abu of KLK14_bABP, are preferred. Conversely, the S4 pocket of KLK14 is large with Trp215 positioned at the base, which is predicted to form π-stacking interactions with large aromatic residues such as the P4 2-Nal of KLK14_bABP.^35^ The selectivity for KLK2_bABP is likely achieved due to the combination of sub-optimal interactions of the S2 and S4 pockets of KLK14 with P2 Tyr and P4 Tle, respectively. Conversely, the KLK14_bABP selectivity is likely derived from the fact that the S4 pocket of KLK2 cannot accept the large aromatic 2-Nal side chain present at P4.

With potent and selective ABPs for KLK2, KLK3 and KLK14 in hand, we next addressed selective readouts of KLK activities in a complex biological system. Compounds **20**, **31** and **39** were incubated with CM obtained from LNCaP-K14 cells, in which KLK14 expression has been placed under the control of a doxycycline-inducible promoter, stimulated with 10 nM DHT and 200 nM doxycycline (dox).^15^ Streptavidin blotting revealed a single target between 25-32 KDa for each compound, labeled in a concentration-dependent manner (**Fig. 4A**), showing that the only detectable targets of these probes are their cognate proteases to the exclusion of other off-targets, emphasizing their impressive selectivity. In line with previous reports KLK14 appears as a double band due to protein glycosylation.^15,30^ Selective covalent modification of KLK2, KLK3 and KLK14 by **31**, **20** and **39**, respectively, was confirmed by streptavidin enrichment and immunoblotting (**Fig. S19 and S20**). In the following experiments, biotinylated ABPs (bABPs) **31**, **20** and **39** are named KLK2_bABP, KLK3_bABP and KLK14_bABP, respectively.

**Figure 4:**
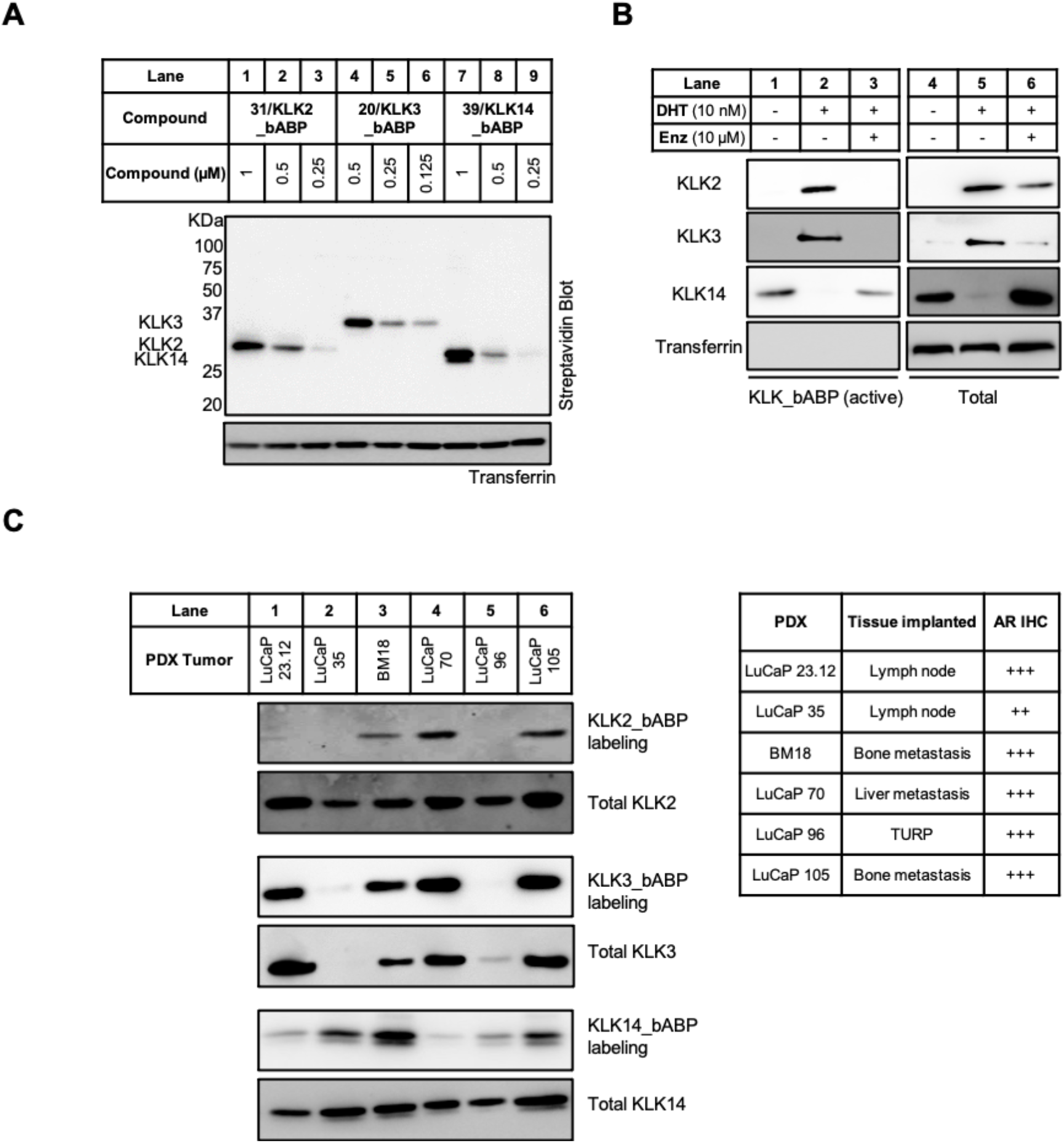
KLK activity profiling in PCa cells and PDX homogenates (**A**) Streptavidin blotting shows potent and selective labeling of KLK2, KLK3 and KLK14 by **31**, **20** and **39** respectively after treatment of LNCaP-K14 conditioned media for 1 h (**B**) KLK activity and expression in the LNCaP cell-line after treatment with 10 nM DHT ± 10 μM Enz. KLK activity was assessed by streptavidin enrichment of probe-labeled KLK molecules following treatment of conditioned media with 1 μM of either **31**, **20** or **39** for 1 h. See Fig. S21 for blot quantification. (**C**) KLK activity and expression in six different PDX homogenates. KLK activity was assessed using the same method as in B. The tissue origin and AR expression levels of the six PDX samples is shown. The + sign indicates the staining intensity of PDX samples by an AR antibody; Abbreviation: IHC = immunohistochemistry, TURP = Transurethral resection of the prostate. See Fig. S22 for blot quantification.

Taken together, detailed analyses of specificity preferences of KLK2, KLK3 and KLK14 led to design and discovery of covalent inhibitors and probes with unprecedented biochemical and cellular potency and selectivity for their cognate proteases.

### Profiling KLK activity in hormone-dependent prostate cancer

As noted above, KLKs 2, 3 and 14 have been proposed to play roles at different stages of PCa progression including facilitating initial prostate tumor expansion and invasion, and establishment of distant metastases.^8^ Given that AR activity is crucial in the development of CRPC and for continued tumor growth, we next assessed how KLK activity changes in response to specific AR signaling perturbations. In the LNCaP-K14 cells used in previous experiments the expression of KLK14 is increased by activation of a doxycycline-inducible promoter, and to assess AR-mediated changes in KLK14 activity we reverted to WT LNCaP cells, which are known to have low basal KLK14 expression.^15^ To enable detection of each KLK in WT LNCaP cells, we incubated LNCaP conditioned media with 1 μM of each KLK ABP, enriched labelled proteins on streptavidin beads and visualized activity levels by immunoblotting with the appropriate KLK antibody (see **Fig. S19** for workflow). As expected, the expression and activity of KLK2 and KLK3 in LNCaP cells was upregulated in response to treatment with DHT and this increase was nullified by co-treatment with Enzalutamide (Enz), a competitive AR inhibitor employed in the clinic to treat CRPC. In contrast, KLK14 expression and activity was decreased upon treatment with DHT but restored by co-incubation with Enz (**Fig. 4B** and **Fig. S21** for blot quantification).

To assess KLK activity levels at different disease stages we profiled six PCa patient-derived xenograft (PDX) tumors isolated from diverse tissues including prostate, and metastases in lymph node, liver and bone.^36^ Fresh-frozen PDX samples were homogenized in 1% triton in PBS and incubated with 1 μM KLK ABP, followed by enrichment and immunoblotting. KLK2 was expressed in all samples, and KLK3 in all but one; however, expression and activity were substantially decoupled, with KLK2 and KLK3 activity seen only in PDXs generated from metastases including lymph node, liver and bone. LuCaP 35 PDX, which lacked KLK3 expression and had decreased KLK2 expression, has also previously been shown to have low AR expression.^36^ KLK14 activity decoupled from expression was evident in all samples, with higher KLK14 activity in LUCaP 35, BM18 and LUCaP 105 (**Fig. 4C** and **Fig. S22** for blot quantification).

Strikingly, we found that simultaneous activation of all three KLKs was observed only in PDX cells from bone metastases (BM18 and LuCaP 105). Advanced PCa has a propensity to metastasize to bone and dysregulate bone resorption and bone formation through complex paracrine signaling events with osteoblasts and osteoclasts (cells which mediate bone generation or absorption, respectively). Treatment options for PCa bone metastases remain inadequate and generally palliative, and there is an urgent need to identify novel therapeutic targets.^37^ Osteoblasts secrete growth factors including IL-6 which can induce AR signaling in PCa cells even in the absence of androgens, as seen during ADT,^38–40^ and we hypothesized that osteoblasts might therefore also induce PCa cell secretion of active KLK molecules into the bone microenvironment under androgen-deprived conditions. To test this hypothesis primary human osteoprogenitor cells were isolated from bone tissue and cultured in osteogenic media (1M β-glycerophosphate, 200 mM ascorbate-2-phosphate and 0.1M dexamethasone) for six weeks as described previously,^41,42^ forming a dense mineralized collagen-rich bone-like matrix (**Fig. S23**). LNCaP cells were then either directly co-cultured with osteoblasts or treated with conditioned media from osteoblasts (here termed ‘indirect co-culture’) in androgen-depleted media (**Fig. 5A**), and CM obtained from indirect or direct co-culture treated with 1 μM of each KLK ABP, followed by enrichment and immunoblotting. In agreement with our hypothesis, both direct and indirect co-culture of LNCaP cells with osteoblasts resulted in an increase of total and active KLK2 and KLK3. In line with the AR-dependent and prostate-restricted expression of these proteases, neither KLK was detected in osteoblast CM alone, and very low expression in CM obtained from androgen-deprived LNCaP cells (**Fig. 5B** and **Fig. S24** for blot quantification). The concentration of active and total KLK14 also increased in co-culture compared to LNCaP cells alone, but in contrast to KLK2 and KLK3 this was due to secretion by osteoblasts, suggesting that osteoblasts may contribute to KLK14 activity in bone metastases. KLK2 and KLK3 stimulate osteoblasts to proliferate, differentiate and secrete growth factors through a TGFβ- dependent mechanism,^43–46^ and these growth factors further enrich the tumor microenvironment and drive pathology in bone metastases.^47,48^ Consequently, our data provides support for a KLK activome-mediated double paracrine signaling mechanism that could contribute towards the establishment of PCa-induced osteoblastic lesions (**Fig. 5C**).

**Figure 5:**
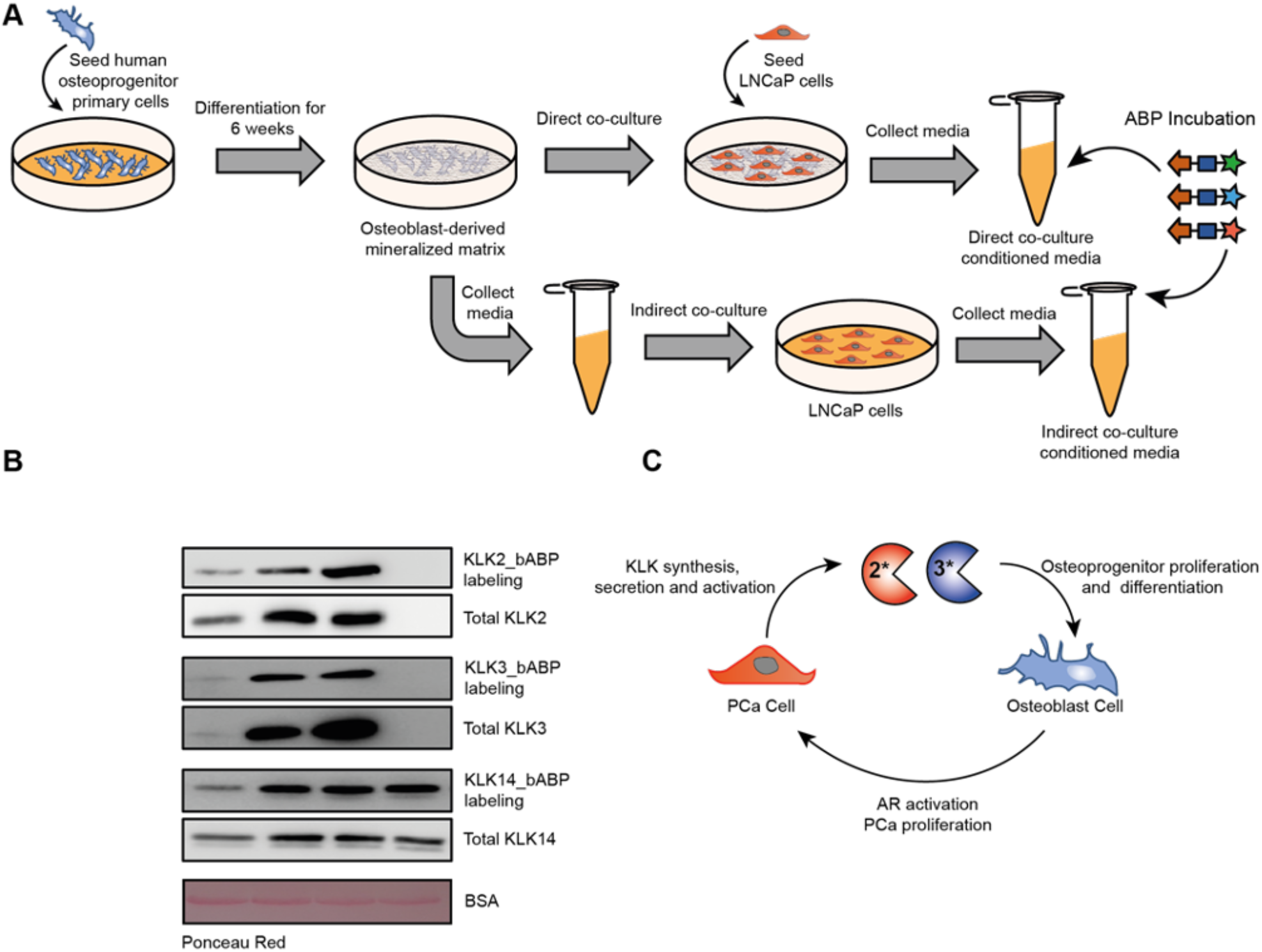
KLK activity profiling in co-culture models of LNCaP cells and primary human osteoblasts. (**A**) Primary human osteoprogenitor cells were seeded and then osteogenically differentiated for 6 weeks resulting in the deposition of a dense, mineralized ECM. In the direct co-culture model LNCaP cells were seeded onto the matrix in androgen-depleted growth media and after 48 hours conditioned media was collected. In the indirect co-culture model osteoblasts were cultured in androgen-deprived growth media for 48 hours, conditioned media was collected and then added to LNCaP cells. The conditioned media was then collected after 48 h. (**B**) KLK activity and expression in the co-culture model of LNCaP cells and osteoblasts. KLK activity was assessed using the same method as described in figure 4B. See Fig. S24 for blot quantification. (**C**) The proposed role of KLK2 and KLK3 in PCa growth in the bone microenvironment. Osteoblasts secrete soluble factors such as IL-6 resulting in ligand-independent activation of the AR. AR signalling results in PCa cell proliferation and secretion of KLK2 and KLK3 into the bone microenvironment. KLK2 and KLK3 stimulate proliferation of osteoprogenitor cells and osteoblast differentiation via a TGFβ-dependent mechanism. This results in further ligand-independent AR activation and establishment of bone metastases.

### Dissecting the role of the KLK activome in prostate cancer progression

The data presented above provide evidence that KLK activity in PCa is regulated by the AR at a level beyond simple changes in expression and raise the possibility that KLK activities may be actionable targets for therapeutic intervention at different disease stages, in particular during PCa metastasis to bone. However, KLKs are predicted to form complex proteolytic networks and it is often not clear which protease(s) should be inhibited for maximum phenotypic response in a particular disease setting. Experiments with purified proteins show that KLK14 can activate pro-KLK2 and pro-KLK3 by proteolytic cleavage of their N-terminal pro-peptide sequence,^17^ and it has been suggested that KLK2 can also activate pro-KLK3.^50^ To enable dissection of the PCa KLK activome directly in LNCaP cells we further expanded the KLK probe toolbox by synthesizing KLK2_fABP (**42**) and KLK14_fABP (**43**) (**Fig. 6A** and **Fig. S2**), fluorescent ABP (fABP) analogues of KLK2_bABP and KLK14_bABP bearing dyes fluorescing at discrete wavelengths, which retain excellent potency and selectivity for KLK2 and KLK14, respectively (**Fig. S25**). KLK2_fABP and KLK14_fABP were combined with KLK3-selective fABP **21** (KLK3_fABP) in an ‘fABP cocktail’ to a final concentration of 1 μM each and incubated for 20 mins with CM obtained from LNCaP-WT, LNCaP-mK14 or LNCaP-K14 cells following 48 h treatment with 10 nM DHT, to enable simultaneous multicolor readout of KLK2, KLK3 and KLK14 activity. Both LNCaP-WT and Dox-treated LNCaP-mK14 cells expressing catalytically inactive mutant KLK14[S195A] under the control of a Dox-inducible promoter^15^ had low levels of active KLK3, while KLK2 and KLK14 activity was below the detection limit (**Fig. 6B**). However, upon Dox-induced KLK14 expression in DHT-treated LNCaP-K14 cells a significant increase in the activity of all three KLKs was evident (**Fig. 6B** and **Fig. S26** for gel quantification). Importantly, the total abundance of secreted KLK2 and KLK3 remained constant upon treatment with Dox, demonstrating an increase in the ratio of active to inactive KLK independent from protein expression. To enable selective inhibition of active KLK molecules, LNCaP-K14 cells were co-incubated with 1 μM KLK2_bABP, KLK3_bABP or KLK14_bABP, 10 nM DHT and 200 nM dox for 48 h. Residual KLK activity in CM was then assessed using the fABP cocktail (**Fig. 6B**). These data firstly demonstrate that even following 48 h incubation KLK bABPs retained exquisite selectivity for their respective target KLK. Secondly, they show that inhibition of either KLK2 or KLK3 activity did not affect the activity of other KLKs, suggesting that KLK2 and KLK3 do not cross-activate the other PCa KLKs. However, inhibition of KLK14 resulted in a significant decrease in the activity levels of both KLK2 and KLK3, suggesting that KLK14 cross-activates the pro-forms of both these proteases.

**Figure 6:**
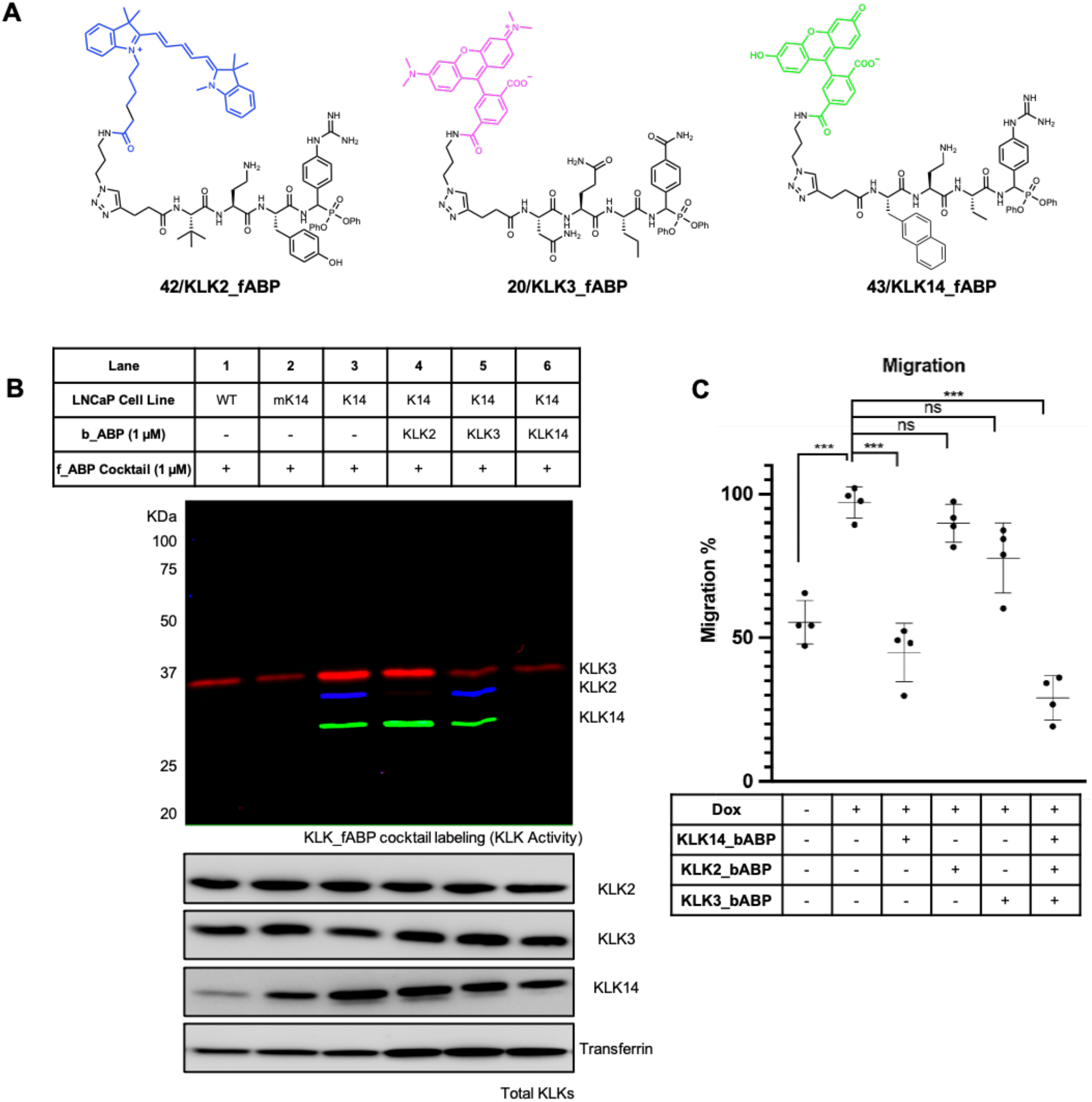
Dissecting the KLK activome in LNCaP cells. (**A**) Structures of fluorescent ABPs **42**, **20** and **43** for KLK2, KLK3 and KLK14, respectively. (**B**) KLK activity profiling by in-gel fluorescence of conditioned media obtained from LNCaP-WT, LNCaP-mK14 and LNCaP-K14 cells following treatment with DHT (10 nM), Dox (100 ng/mL) and the appropriate biotinylated ABP using fluorescent ABPs **20**, **42** and **43**. See Fig. S26 for blot and gel quantification. (**C**) Assessment of KLK activity inhibition on LNCaP cell migration. LNCaP-K14 cells were cultured in serum-free RPMI media supplemented with 10 nM DHT ± Dox (100 ng/mL) for 24 h. Cells were added to transwell inserts (100,000 cells in 100 μL of serum-free RPMI) and after 4 h the appropriate ABP was added, and cultures were incubated for a further 24 h before the number of migrating cells was quantified.

Prior to the establishment of bone metastases circulating PCa cells must first enter the bone microenvironment by migrating across the sinusoidal wall.^49^ We have previously shown that increased KLK14 activity drives PCa cell migration and colonization of mineralized bone matrices.^15^However, the specific contributions of the KLK activome (KLK2, KLK3 and KLK14) towards PCa cell migration is yet to be elucidated. We first confirmed that treatment of LNCaP-K14 cells with 10 μM of KLK2_bABP, KLK3_bABP and KLK14_bABP resulted in no cytotoxicity over a 96-h time period (**Fig. S27**). We then assessed the migratory capacity of LNCaP-K14 cells using a transwell assay (see **Fig. S28** for standard curve). Induction of KLK14 expression in LNCaP-K14 cells (200 nM Dox) resulted in increased migration, as expected, which was nullified by co-treatment with 1 μM KLK14_bABP (**Fig. 6C**). Conversely, treatment with 1 μM KLK2_bABP or KLK3_bABP had no significant effect on the number of migrating LNCaP-K14 cells. Simultaneous treatment of LNCaP-K14 cells with 1 μM each KLK2_bABP, KLK3_bABP and KLK14_bABP resulted in a significant decrease in cell migration, suggesting that KLK14 activity is a key driver of PCa cell migration, while KLK2 and KLK3 activities individually make minor contributions.

Overall, these data dissect the KLK activome in LNCaP cells and show that KLK14 activity drives PCa cell migration.

## Discussion

Despite recent progress in determining the pathophysiological roles of individual KLKs at different stages of PCa, the potential of selectively inhibiting these proteases for therapeutic intervention is yet to be realized.^51^ It has become clear that KLKs do not work alone, but instead are individual components of complex networks that when deregulated drive disease progression through amplified proteolysis.^52–54^ However, delineating the overlapping, synergistic and opposing activities of individual prostatic KLKs in complex environments remains very challenging. We therefore developed the first chemical toolkit enabling simultaneous assessment of the activity of each PCa-relevant KLK in PCa samples. Peptidyl-DPP ABPs were established for each protease, and potency and specificity further optimized in PCa supernatants by competitive ABPP, an approach that enables assessment of the properties of each covalent inhibitor directly in a complex biological environment. Development of selective ABPs for KLK2 and KLK14 was complicated by their similar trypsin-like activity; indeed, proteases with the same primary (P1) specificity are often co-expressed in tissues, and to generate selective tools it is necessary to explore more complex specificity determinants.^55^ Here we applied positional scanning libraries to identify mutually exclusive preferences in the S2-S4 pockets of KLK2 and KLK14; however this approach does not account for potential cooperativity between protease subsites in driving selectivity. Further optimization across ABPs incorporating different combinations of preferred amino acids illustrated the importance of exploiting cooperativity, and in the case of KLK14 identified an ABP with optimal selectivity which features none of the top amino acid hits identified for each individual sub-site in the library screen. Interestingly, both KLK2 and KLK14 demonstrated dual specificity in certain sub-pockets, suggesting a role for these subsites in tuning substrate profiles.^56^ We note that whilst the present set of probes has been optimized for PCa, each tissue type expresses a different set of secreted proteases and thus the specificity sequence required to obtain probe selectivity may depend on the physiological context.

Recent development of KLK-knockout and transgenic mice^18,57^ and selective inhibitors^30^ have revealed key associations between individual KLK activities and the onset or progression of diverse diseases. However, whether these associations are due to the activity of an individual KLK or because of cross-activation of other proteases, including other KLKs, is poorly understood. The KLK activome has to date been modeled only using purified proteins, and it remains very challenging to deconvolute the biological relevance of a specific cross-activation event in more complex systems.^16,17^ We leveraged our chemical toolkit to dissect the KLK activome in PCa cells by inhibiting one KLK and assessing the change in activity of the remaining KLKs, demonstrating that KLK14 cross-activates KLK2 and KLK3. We suggest that our approach could be used in principle to dissect any disease-related or tissue KLK activome and reveal which KLKs represent potential targets for therapeutic intervention in a specific context.

Our data using LNCaP cells demonstrates that KLK activity in the tumor microenvironment is likely regulated by AR signaling **(Fig. 4B**). However, while KLK2 and KLK3 activity is upregulated by AR activation, KLK14 activity increases upon AR inhibition. The latter observation suggests a role for active KLK14 in the development of resistance to AR competitive inhibitors. Despite the divergent response to AR signaling, we show that KLK2, KLK3 and KLK14 are co-expressed and active in hormone responsive cells from different sites of PCa metastasis, including lymph nodes, liver and bone. An explanation for this paradox lies in the heterogeneity of prostate tumors, which develop resistance to AR inhibition through diverse mechanisms including upregulation of AR-V7 (an AR splice-variant with constitutive transcriptional activity that lacks a ligand-binding domain) and glucocorticoid receptor (GR).^58–60^ Both AR-V7 and GR signaling drive increased KLK2 and KLK3 expression even in the face of Enz treatment. Similarly, a subset of AR-null PCa cells have high KLK14 expression, which is not perturbed by DHT treatment.^15^ We hypothesize that these diverse PCa cell populations contribute to a buildup of KLK activities in the tumor microenvironment.

In this study we found that androgen-independent activation of AR in PCa cells by osteoblasts results in an increase of KLK2 and KLK3 activity in the bone microenvironment. Previous studies have shown that KLK3 increases bone volume and osteoblast numbers *in vivo* via a TGFβ-dependent mechanism.^43–46^ It has also been demonstrated *in vitro* that KLK2 and KLK14 can activate TGFβ-1 and TGFβ-2.^61^ Furthermore, our data show that osteoblasts, as well as PCa cells, secrete active KLK14 into the bone microenvironment and that KLK14 can cross-activate KLK2 and KLK3, thus providing a proteolytic cascade which may amplify osteoblast proliferation and differentiation and result in a double paracrine signaling event that drives both tumor growth and woven bone formation. We propose that the KLK activome warrants further study as an actionable therapeutic target in bone metastatic PCa, either alone or in combination with enzalutamide.

Finally, the chemical tools developed in this study may allow KLK activity to be explored as a novel biomarker in PCa. Population-based screening with the PSA test has decreased PCa mortality; however, due to the test’s relatively poor specificity, it has also increased the detection of indolent cancers and resulted in overtreatment of patients. Identification of novel biomarkers with increased specificity for aggressive PCa detection may aid in risk stratification and the appropriate identification of men for prostate biopsy.^9^ An early hallmark of PCa is the downregulation of zinc transporter proteins, which results in a decrease in the concentration of Zn^2+^ ions in the prostate.^62^ Zinc ions allosterically regulate the activity of KLKs and thus there is an increase in KLK activity in prostatic fluid obtained from patients with PCa.^19^ Measurement of KLK activity in first void urine samples, which contain prostatic fluid, may be worth exploring in the quest for a more accurate diagnostic tool.^63–65^ To this end, we note that biotinylated ABPs have previously been integrated into ELISA platforms to enable highly sensitive and high throughput assessment of protease activity.^66^ Similarly, as KLK activity is higher in PCa tissue compared to neighboring healthy tissue, KLK probes could in future be used as fluorescent guides for surgeons striving to obtain negative margins during tumor resection,^67^ potentially facilitated by the development of quenched-fluorescent derivatives.^68,69^

In conclusion, this study describes the development of a versatile chemical probe platform to enable dissection of KLK activome activity in PCa, leading support for the hypothesis that the KLK activome drives PCa progression and holds promise as an actionable therapeutic target in CRPC.

## Acknowledgments

The authors thank L. Haigh (Department of Chemistry Mass Spectrometry Facility, Imperial College London) for assistance in acquiring high-resolution mass spectrometry (HRMS) data. S.L. was supported by the EPSRC Centre for Doctoral Training in Physical Sciences Innovation in Chemical Biology for Bioindustry and Healthcare (Grant EP/LO15498/1) and an EPSRC Doctoral Prize fellowship. M.M thanks the Xunta de Galicia for a postdoctoral fellowship. E.D.V. is supported by a H2020 MSCA-IF fellowship (890900). L.Z. was supported by Cancer Research UK (grant C24523/A25192). E.W.T. thanks Worldwide Cancer Research for support (grant 19-0059). A.N. was supported by a postdoctoral mobility fellowship from the Swiss National Science Foundation.

---

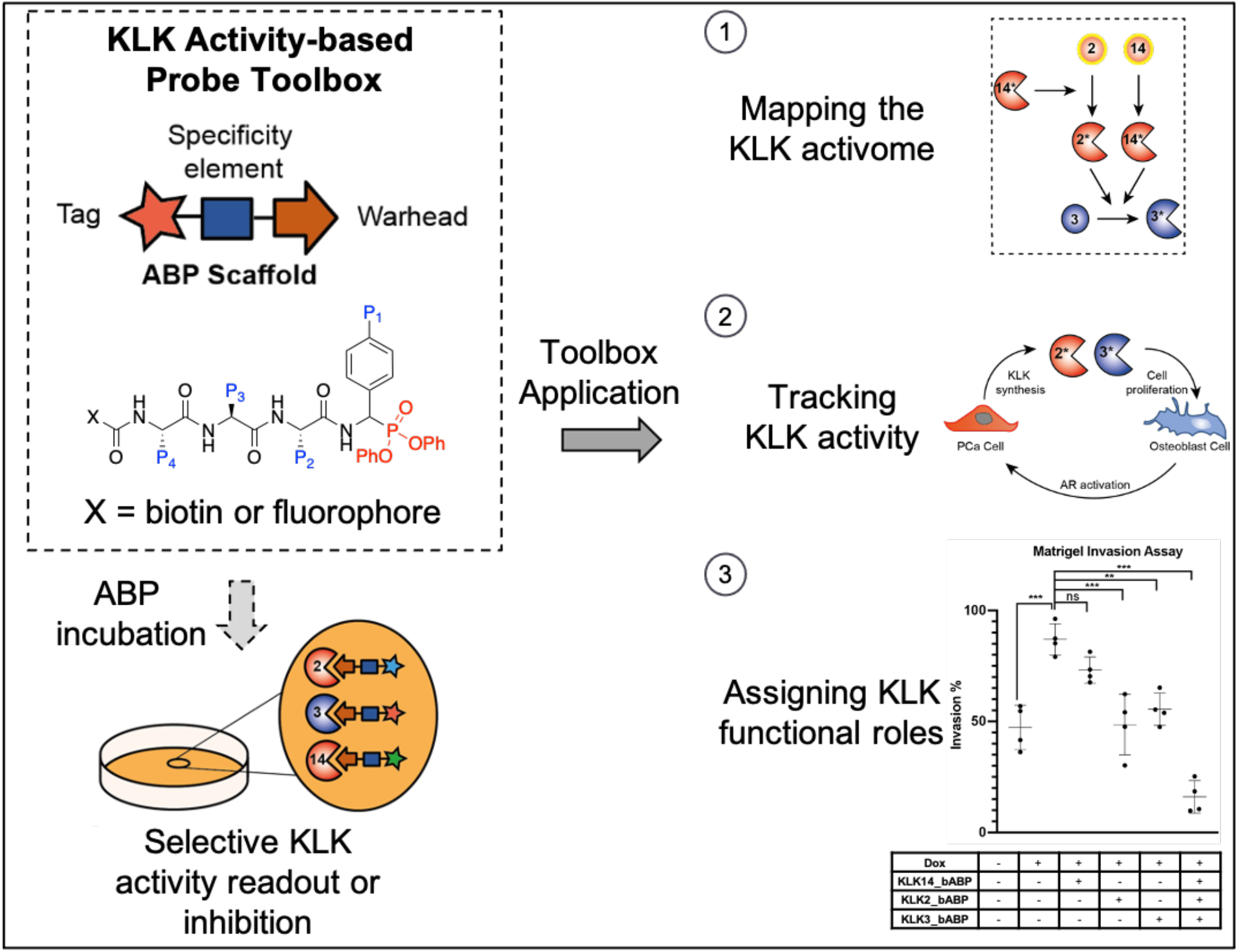

## References

1. Pascale, M. et al. The outcome of prostate cancer patients treated with curative intent strongly depends on survival after metastatic progression. BMC Cancer 17, (2017).

2. Ryan, C. J. et al. Abiraterone acetate plus prednisone versus placebo plus prednisone in chemotherapy-naive men with metastatic castration-resistant prostate cancer (COU-AA-302): final overall survival analysis of a randomised, double-blind, placebo-controlled phase 3 study. Lancet Oncol. 16, 152–160 (2015).

3. Rice, M. A., Malhotra, S. V. & Stoyanova, T. Second-generation antiandrogens: From discovery to standard of care in castration resistant prostate cancer. Frontiers in Neurology vol. 10 801 (2019).

4. Body, J.-J., Casimiro, S. & Costa, L. Targeting bone metastases in prostate cancer: improving clinical outcome. Nat. Rev. Urol. 12, 340–356 (2015).

5. Watson, P. A., Arora, V. K. & Sawyers, C. L. Emerging mechanisms of resistance to androgen receptor inhibitors in prostate cancer. Nat. Rev. Cancer 15, 701–711 (2015).

6. Morova, T. et al. Androgen receptor-binding sites are highly mutated in prostate cancer. Nat. Commun. 11, 832 (2020).

7. Prassas, I., Eissa, A., Poda, G. & Diamandis, E. P. Unleashing the therapeutic potential of human kallikrein-related serine proteases. Nat. Rev. Drug Discov. 14, 183–202 (2015).

8. Fuhrman-Luck, R. A., Loessner, D. & Clements, J. A. Kallikrein-Related Peptidases in Prostate Cancer: From Molecular Function to Clinical Application. EJIFCC 25, 269–81 (2014).

9. Lilja, H., Ulmert, D. & Vickers, A. J. Prostate-specific antigen and prostate cancer: prediction, detection and monitoring. Nat. Rev. Cancer 8, 268–278 (2008).

10. Darst, B. F. et al. The Four-Kallikrein Panel Is Effective in Identifying Aggressive Prostate Cancer in a Multiethnic Population. Cancer Epidemiol. Biomarkers Prev. 29, 1381–1388 (2020).

11. Mize, G. J., Wang, W. & Takayama, T. K. Prostate-specific kallikreins−2 and −4 enhance the proliferation of DU-145 prostate cancer cells through protease-activated receptors−1 and −2. Mol. Cancer Res. 6, 1043–1051 (2008).

12. Yonou, H. et al. Prostate-specific antigen stimulates osteoprotegerin production and inhibits receptor activator of nuclear factor-KappaB ligand expression by human osteoblasts. Prostate 67, 840–848 (2007).

13. Nadiminty, N. et al. Prostate-Specific Antigen Modulates Genes Involved in Bone Remodeling and Induces Osteoblast Differentiation of Human Osteosarcoma Cell Line SaOS-2. Clin. Cancer Res. 12, 1420–1430 (2006).

14. Lose, F. et al. The kallikrein 14 gene is down-regulated by androgen receptor signalling and harbours genetic variation that is associated with prostate tumour aggressiveness. Biol. Chem. 393, 403–412 (2012).

15. Kryza, T. et al. The molecular function of kallikrein-related peptidase 14 demonstrates a key modulatory role in advanced prostate cancer. Mol. Oncol. 14, 105–128 (2020).

16. Yoon, H. et al. Activation Profiles and Regulatory Cascades of the Human Kallikrein-related Peptidases. J. Biol. Chem. 282, 31852–31864 (2007).

17. Yoon, H. et al. A completed KLK activome profile: investigation of activation profiles of KLK9, 10, and 15. Biol. Chem. 390, 373–7 (2009).

18. Kasparek, P. et al. KLK5 and KLK7 Ablation Fully Rescues Lethality of Netherton Syndrome-Like Phenotype. PLoS Genet. 13, e1006566 (2017).

19. Goettig, P., Magdolen, V. & Brandstetter, H. Natural and synthetic inhibitors of kallikrein-related peptidases (KLKs). Biochimie 92, 1546 (2010).

20. Rawlings, N. D. et al. The MEROPS database of proteolytic enzymes, their substrates and inhibitors in 2017 and a comparison with peptidases in the PANTHER database. Nucleic Acids Res. 46, D624–D632 (2018).

21. Maślanka, M. & Mucha, A. Recent Developments in Peptidyl Diaryl Phoshonates as Inhibitors and Activity-Based Probes for Serine Proteases. Pharmaceuticals (Basel). 12, (2019).

22. Kojtari, A., Shah, V., Babinec, J. S., Yang, C. & Ji, H.-F. Structure-Based Drug Design of Diphenyl α-Aminoalkylphosphonates as Prostate-Specific Antigen Antagonists. J. Chem. Inf. Model. 54, 2967–2979 (2014).

23. LeBeau, A. M., Banerjee, S. R., Pomper, M. G., Mease, R. C. & Denmeade, S. R. Optimization of peptide-based inhibitors of prostate-specific antigen (PSA) as targeted imaging agents for prostate cancer. Bioorg. Med. Chem. 17, 4888–4893 (2009).

24. Kasperkiewicz, P., Altman, Y., D’Angelo, M., Salvesen, G. S. & Drag, M. Toolbox of Fluorescent Probes for Parallel Imaging Reveals Uneven Location of Serine Proteases in Neutrophils. J. Am. Chem. Soc. 139, 10115–10125 (2017).

25. Wright, M. H. & Sieber, S. A. Chemical proteomics approaches for identifying the cellular targets of natural products. Nat. Prod. Rep. 33, 681–708 (2016).

26. Malm, J., Hellman, J., Hogg, P. & Lilja, H. Enzymatic action of prostate-specific antigen (PSA or hK3): substrate specificity and regulation by Zn(2+), a tight-binding inhibitor. Prostate 45, 132–9 (2000).

27. LeBeau, A. M., Singh, P., Isaacs, J. T. & Denmeade, S. R. Potent and selective peptidyl boronic acid inhibitors of the serine protease prostate-specific antigen. Chem. Biol. 15, 665–74 (2008).

28. Poreba, M., Salvesen, G. S. & Drag, M. Synthesis of a HyCoSuL peptide substrate library to dissect protease substrate specificity. Nat. Protoc. 12, 2189–2214 (2017).

29. van Soom, J. et al. The first potent diphenyl phosphonate KLK4 inhibitors with unexpected binding kinetics. Medchemcomm 6, 1954–1958 (2015).

30. de Veer, S. J. et al. Selective Substrates and Inhibitors for Kallikrein-Related Peptidase 7 (KLK7) Shed Light on KLK Proteolytic Activity in the Stratum Corneum. J. Invest. Dermatol. 137, 430–439 (2017).

31. Skala, W. et al. Structure-function analyses of human kallikrein-related peptidase 2 establish the 99-loop as master regulator of activity. J. Biol. Chem. 289, 34267–34283 (2014).

32. Ménez, R. et al. Crystal Structure of a Ternary Complex between Human Prostate-specific Antigen, Its Substrate Acyl Intermediate and an Activating Antibody. J. Mol. Biol. 376, 1021–1033 (2008).

33. Chemical Computing Group ULC. Molecular Operating Environment (MOE). 1010 Sherbooke St. West, Suite #910, Montreal, QC, (2020).

34. Singh, P., LeBeau, A. M., Lilja, H., Denmeade, S. R. & Isaacs, J. T. Molecular insights into substrate specificity of prostate specific antigen through structural modeling. Proteins Struct. Funct. Bioinforma. 77, 984–993 (2009).

35. De Veer, S. J., Swedberg, J. E., Parker, E. A. & Harris, J. M. Non-combinatorial library screening reveals subsite cooperativity and identifies new high-efficiency substrates for kallikrein-related peptidase 14. in Biological Chemistry vol. 393 331–341 (Biol Chem, 2012).

36. Nguyen, H. M. et al. LuCaP Prostate Cancer Patient-Derived Xenografts Reflect the Molecular Heterogeneity of Advanced Disease an--d Serve as Models for Evaluating Cancer Therapeutics. Prostate 77, 654–671 (2017).

37. Nevedomskaya, E., Baumgart, S. J. & Haendler, B. Recent advances in prostate cancer treatment and drug discovery. International Journal of Molecular Sciences vol. 19 (2018).

38. Lu, Y. et al. Osteoblasts induce prostate cancer proliferation and PSA expression through interleukin-6-mediated activation of the androgen receptor. Clin. Exp. Metastasis 21, 399–408 (2004).

39. Sung, S.-Y. et al. Coevolution of prostate cancer and bone stroma in three-dimensional coculture: implications for cancer growth and metastasis. Cancer Res. 68, 9996–10003 (2008).

40. Nguyen, D. P., Li, J. & Tewari, A. K. Inflammation and prostate cancer: The role of interleukin 6 (IL-6). BJU International vol. 113 986–992 (2014).

41. Bock, N. et al. Engineering osteoblastic metastases to delineate the adaptive response of androgen-deprived prostate cancer in the bone metastatic microenvironment. Bone Res. 7, 13 (2019).

42. Shokoohmand, A. et al. Microenvironment engineering of osteoblastic bone metastases reveals osteomimicry of patient-derived prostate cancer xenografts. Biomaterials 220, 119402 (2019).

43. Killian, C. S., Corral, D. A., Kawinski, E. & Constantine, R. I. Mitogenic Response of Osteoblast Cells to Prostate-Specific Antigen Suggests an Activation of Latent TGF-β and a Proteolytic Modulation of Cell Adhesion Receptors. Biochem. Biophys. Res. Commun. 192, 940–947 (1993).

44. Dallas, S. L. et al. Preferential production of latent transforming growth factor ?-2 by primary prostatic epithelial cells and its activation by prostate-specific antigen. J. Cell. Physiol. 202, 361–370 (2005).

45. Yonou, H. et al. Prostate-Specific Antigen Induces Osteoplastic Changes by an Autonomous Mechanism. Biochem. Biophys. Res. Commun. 289, 1082–1087 (2001).

46. Cumming, A. P., Hopmans, S. N., Vukmirović-Popović, S. & Duivenvoorden, W. C. PSA affects prostate cancer cell invasion in vitro and induces an osteoblastic phenotype in bone in vivo. Prostate Cancer Prostatic Dis. 14, 286–294 (2011).

47. Guise, T. A. et al. Basic Mechanisms Responsible for Osteolytic and Osteoblastic Bone Metastases. Clin. Cancer Res. 12, 6213s–6216s (2006).

48. Iwamura, M., Hellman, J., Cockett, A. T. K., Lilja, H. & Gershagen, S. Alteration of the hormonal bioactivity of parathyroid hormone-related protein (PTHrP) as a result of limited proteolysis by prostate-specific antigen. Urology 48, 317–325 (1996).

49. Mundy, G. R. Metastasis to bone: Causes, consequences and therapeutic opportunities. Nature Reviews Cancer vol. 2 584–593 (2002).

50. Williams, S. A., Xu, Y., De Marzo, A. M., Isaacs, J. T. & Denmeade, S. R. Prostate-specific antigen (PSA) is activated by KLK2 in prostate cancer ex vivo models and in prostate-targeted PSA/KLK2 double transgenic mice. Prostate 70, 788–96 (2010).

51. Sotiropoulou, G. & Pampalakis, G. Targeting the kallikrein-related peptidases for drug development. Trends Pharmacol. Sci. 33, 623–34 (2012).

52. Beaufort, N. et al. Interdependence of kallikrein-related peptidases in proteolytic networks. Biol. Chem. 391, 581–7 (2010).

53. Ohler, A., Debela, M., Wagner, S., Magdolen, V. & Becker-Pauly, C. Analyzing the protease web in skin: meprin metalloproteases are activated specifically by KLK4, 5 and 8 vice versa leading to processing of proKLK7 thereby triggering its activation. Biol. Chem. 391, 455–60 (2010).

54. Tanaka, R. J., Ono, M. & Harrington, H. A. Skin Barrier Homeostasis in Atopic Dermatitis: Feedback Regulation of Kallikrein Activity. PLoS One 6, e19895 (2011).

55. Kasperkiewicz, P., Poreba, M., Groborz, K. & Drag, M. Emerging challenges in the design of selective substrates, inhibitors and activity-based probes for indistinguishable proteases. FEBS J. 284, 1518–1539 (2017).

56. Shaw, J. L. & Diamandis, E. P. Distribution of 15 Human Kallikreins in Tissues and Biological Fluids. Clin. Chem. 53, 1423–1432 (2007).

57. Furio, L. et al. Transgenic kallikrein 5 mice reproduce major cutaneous and systemic hallmarks of Netherton syndrome. J. Exp. Med. 211, 499–513 (2014).

58. Shah, R. B. et al. Androgen-independent prostate cancer is a heterogeneous group of diseases: lessons from a rapid autopsy program. Cancer Res. 64, 9209–16 (2004).

59. Arora, V. K. et al. Glucocorticoid receptor confers resistance to antiandrogens by bypassing androgen receptor blockade. Cell 155, 1309–22 (2013).

60. Guo, Z. et al. A Novel Androgen Receptor Splice Variant Is Up-regulated during Prostate Cancer Progression and Promotes Androgen Depletion-Resistant Growth. Cancer Res. 69, 2305–2313 (2009).

61. Emami, N. & Diamandis, E. P. Potential role of multiple members of the kallikrein-related peptidase family of serine proteases in activating latent TGF beta 1 in semen. Biol. Chem. 391, 85–95 (2010).

62. Costello, L. C., Feng, P., Milon, B., Tan, M. & Franklin, R. B. Role of zinc in the pathogenesis and treatment of prostate cancer: critical issues to resolve. Prostate Cancer Prostatic Dis. 7, 111–7 (2004).

63. CV, O. et al. Human Kallikrein 4: Quantitative Study in Tissues and Evidence for Its Secretion Into Biological Fluids. Clin. Chem. 51, (2005).

64. Theodorescu, D. et al. Discovery and validation of urinary biomarkers for prostate cancer. Proteomics. Clin. Appl. 2, 556–570 (2008).

65. Webb, M. et al. Methodology for the at-home collection of urine samples for prostate cancer detection. Biotechniques 68, 65–71 (2020).

66. Eitelhuber, A. C. et al. Activity-based probes for detection of active MALT1 paracaspase in immune cells and lymphomas. Chem. Biol. 22, 129–38 (2015).

67. Nguyen, Q. T. & Tsien, R. Y. Fluorescence-guided surgery with live molecular navigation — a new cutting edge. Nat. Rev. Cancer 13, 653–662 (2013).

68. Oresic Bender, K. et al. Design of a Highly Selective Quenched Activity-Based Probe and Its Application in Dual Color Imaging Studies of Cathepsin S Activity Localization. J. Am. Chem. Soc. 137, 4771–4777 (2015).

69. Serim, S., Baer, P. & Verhelst, S. H. L. Mixed alkyl aryl phosphonate esters as quenched fluorescent activity-based probes for serine proteases. Org. Biomol. Chem. 13, 2293–2299 (2015).

