## Supplementary Information for "A suite of activity-based probes to dissect the KLK activome in drug-resistant prostate cancer"

### 1 Materials and Methods

#### 1.1 Chemicals and Reagents

All chemicals and reagents were purchased from commercial suppliers and used without further purification. Rink Amide resin (100-200 mesh) and 2-Chlorotrityl chloride resin (100-200 mesh, 1% DVB) were obtained from Merck. Natural Fmoc-amino acids were obtained from AGTC Bioproducts Ltd. Unnatural Fmoc-amino acids were obtained from Fluorochem, Alfa Aesar, VWR, Merck, Iris Biotech, Bachem or Sigma Aldrich. Culture media, fetal bovine serum, trypsin, antibiotics and chemicals required for differentiation of osteoprogenitor cells were obtained from Sigma Aldrich, Gibco or Invitrogen. Amicon 3 KDa Centrifugal filter units were obtained from Merck. Recombinant human KLK2 and KLK14 were obtained from R&D Systems and activated using the supplier's protocol. All remaining chemical reagents were obtained from VWR, Sigma Aldrich or Merck.

#### 1.2 Equipment List

In-gel fluorescence was recorded using a Typhoon FLA 9500 Imaging System (GE Healthcare) at Cy-2, Cy-3 or Cy-5 wavelengths. Western blots were visualized by chemiluminescence using a LAS-4000 Imaging System (FujiFilm). Fluorescent plate-based enzymatic assays were carried out using an Envision 2104 Multilabel Plate Reader (PerkinElmer) operating in kinetic mode for a minimum of 20 min with excitation/emission wavelengths of 355 nm/460 nm.

### 2 Biological and Biochemical Methods

#### 2.1 Cell Culture

**Routine cell culture conditions**: Wild-type LNCaP cells (passage <15) were grown in RPMI media supplemented with 5 % FBS. LNCaP cells transfected with the coding sequence for mKLK14 or wild-type KLK14 were grown using the same conditions but also with the addition of puromycin (1 µg/mL). Human osteoprogenitor cells were grown in Minimal Essential Media (MEM) supplemented with 10 % FBS and 1 % penicillin – streptomycin (P/S).

**Androgen-signaling Experiments**: LNCaP cells grown to 80 % confluency in a T175 plate were incubated with phenol red-free RPMI media supplemented with 5 % charcoal-stripped FBS for 24 h. After 24 h, the media was aspirated, and the cells were incubated with serum-free RPMI media (0 % FBS and phenol red-free) supplemented with the appropriate treatment condition (10 nM DHT or 10 nM DHT + 10 µM Enzalutamide) for a further 48 h. After 48 h the media was collected, passed through a 0.2 µm syringe filter and stored at – 80 °C until needed.

**Doxycycline induction of mKLK14- and KLK14-LNCaP cells**: Cell lines were processed using the same method as for androgen-signaling experiments. However, serum-free RPMI media was supplemented with 10 nM DHT and 100 ng/mL of doxycycline.

**Primary cell isolation from bone tissue:** Human primary osteoprogenitor cells were isolated from hip and knee bone tissue obtained under informed consent from male donors undergoing joint replacement surgery (QUT ethics approval number 1400001024, surgery performed by Professor Ross Crawford, QUT), as described previously.[1] Non-sclerotic, trabecular bone was collected either from the tibial plateau/femoral condyles from the knee or from the acetabular ground from the hip. Bone fragments (4-5 mg) were minced and washed in sterile phosphate buffer saline (PBS, Gibco, Australia). Bone fragments were transferred to flasks and grown under growth media (GM), containing α-MEM with ribonucleosides, deoxyribonucleosides, phenol Red and L-glutamine, with 10% fetal bovine serum (FBS) and 1% penicillin/streptomycin (P/S, 10,000 U/mL stock solution), all from Gibco. Osteoprogenitor cell outgrowth occurred after 7-10 days. Cells were collected using 0.25% w/v Trypsin – 1 mM EDTA (Gibco) at 80% confluence, and further expanded using GM and frozen down in 90% FBS and 10% DMSO until used for experiments (under 5 passages).

**PCa-Osteoblast Co-Culture**: Osteoprogenitor cells (passage 3) were seeded in T175 flasks at a density of 3000 cells/cm$^2$ and cultured under standard conditions until 100 % confluent. At 100 % confluency, the cells were incubated with osteogenic media (1 M β-glycerophosphate, 50 mg/mL ascorbate-2-phosphate and 0.1 M dexamethasone) and cultured for a further 6 weeks with media changes every 3 days. **Direct Co-culture** - After 6 weeks, the media was aspirated and LNCaP cells (5 million cells in 25 mL of RPMI with 5 %

FBS) were seeded onto the bone matrix. As a control the same number of LNCaP cells were seeded in a T175 plate lacking osteoblast cells. After 12 h, the media was aspirated, and the co-culture was incubated with serum-free RPMI media (0 % FBS and phenol red-free) for a further 48 h. After 48 h the media was collected, passed through a 0.2 µm syringe filter and stored at – 80 °C until needed. **Indirect Co-culture** – After 6 weeks, the media was aspirated, and the differentiated osteoblasts were incubated with serum-free RPMI media (0 % FBS and phenol red-free) for a further 48 h. After 48 h the media was collected, passed through a 0.2 µm syringe filter and then added to a T175 plate of LNCaP cells grown to 80 % confluency and hormone starved for 24 h in RPMI media supplemented with 5 % charcoal stripped FBS. After 48 h the media was collected, passed through a 0.2 µm syringe filter and stored at – 80 °C until needed.

#### 2.2 Media and tumor sample preparation

**Preparation of PDX homogenates**: PCa PDX lines were maintained as subcutaneous tumors in the lateral flanks of SCID mice as described previously.[2] This was approved by the Animal Research Ethics Committees (TRI/QUT/370/17, QUT1800000289). All PDX work was carried out in accordance with the Australian National Health & Medical Research Council (NHMRC) guidelines for the care and use of laboratory animals. Following excision tumors were snap frozen and stored at – 80 °C. Approximately 20 mg of fresh-frozen PDX sample was removed from – 80 °C storage and homogenized in 1 % triton in PBS using a BeadBlaster24 Tissue Homogenizer (Benchmark Scientific). Homogenized samples were centrifuged for 5 min (13,000 rpm, 4 °C) before being passed through a 0.2 µm syringe filter. The protein concentration of the homogenate was measured using the DC Protein Assay (BioRad), following the manufacturer's instructions. Samples were then diluted to approximately 1 mg/mL and stored at -80 °C until further use.

**Preparation of conditioned media**: Samples of conditioned media were removed from – 80 °C storage and concentrated by centrifugation (3000 g, 10 °C) in a 3 KDa molecular weight cut-off spin filter (Amicon) for approximately 3 h. This reduced the sample volume from 25 mL to approximately 1 mL. The protein concentration of the media was measured using the DC Protein Assay (BioRad), following the manufacturer's instructions. Samples were then diluted to approximately 1 mg/mL, aliquoted in to 100 µg fractions and stored at -80 °C until further use.

#### 2.3 Probe Labeling Protocols

**Streptavidin Blot:** LNCaP conditioned media (10 µg) was treated with a biotinylated activity-based probe for 1 h. After 1 h, 3 µL of sample loading buffer (250 mM Tris, 8 % SDS (w/v),

0.2 % bromophenol blue (w/v), 40 % glycerol (v/v), 10 % β-mercaptoethanol, pH 6.8) was added and the sample was boiled for 5 min. Aliquots were then loaded on to hand cast 12 % Bis-Tris gels. SDS-PAGE gels were typically run for 10 min at 80 volts and 50 min at 180 volts. Proteins were transferred to a nitrocellulose membrane (GE Healthcare, Hybond ECL, pore size 0.45 μm) using a wet transfer setup and a Tris-glycine transfer buffer supplemented with 20 % MeOH. Membranes were washed with TBS-T (1 x TBS, 0.1 % Tween-20), blocked for 1 h with a 3 % BSA solution in TBS-T and incubated with Streptavidin-HRP (1:1000 dilution in 0.3 % BSA) for 1 h. After 1 h, the membrane was washed with TBS-T (3 x 10 min) and then developed with Luminata Crescendo Western HRP substrate (Millipore), according to the manufacturer's instructions.

**Streptavidin Enrichment**: LNCaP conditioned media or PDX homogenate (200 μg, 1 mg/mL) was treated with a biotinylated activity-based probe for 1 h. After 1 h, proteins were precipitated using a chloroform/methanol precipitation procedure: 2 vol. MeOH, 0.5 vol. $CHCl_3$ and 1 vol. $H_2O$ was added to samples, which were then centrifuged at 17,000 g for 5 min. The top organic layer was removed and an additional 4 vol. of MeOH was added to the sample, which was centrifuged as before. The liquid was carefully decanted, and the protein pellet was left to dry for exactly 5 min at RT. The pellet was then resuspended in 2 % SDS, 10 mM DTT in PBS to a concentration of 10 mg/mL before being diluted 10-fold in 1 x PBS to give a 1 mg/mL protein solution. Next, 10 μL of protein was reserved for the 'before pull-down' sample and the remainder was added to 25 μL of pre-washed (3 x 200 μL) Streptavidin C1 Dynabeads ® MyOne™ (Thermo Fisher Scientific) followed by gentle agitation for 2 h at RT. After 2 h, the supernatant was removed (retaining 10 μL for the 'supernatant' sample) and the beads were washed with 1 % SDS in PBS (3 x 200 μL). 15 μL of PBS and 5 μL of sample loading buffer were added to the beads and the resulting mixture was boiled for 10 min. 10 μL of before pull-down sample, 10 μL of supernatant sample and 20 μL of 'pull-down' sample were loaded on to a hand cast 12 % Bis-Tris gel, which was run using the same conditions as described for the streptavidin blotting experiments. Proteins were transferred to a nitrocellulose membrane using the same procedure as above. After blocking with 5 % milk in TBS-T for 30 min, the membrane was incubated with the appropriate KLK antibody (1:1000, 5 % milk in TBS-T) overnight at 4 °C, washed with TBS-T (3 x), incubated with the appropriate HRP-conjugated secondary antibody (1:10,000, 5 % milk in TBS-T) for 1 h at RT, washed with TBS-T (3 x) and developed as described above. For a list of antibodies used throughout this study see Table S1.

**Table S1**: Antibodies used throughout this project

| Protein | Supplier | Cat. No. | Host | /clonal | Immunogen |
|---|---|---|---|---|---|
| KLK2 | Abcam | Ab124899 | Rabbit | Mono | Synthetic peptide within amino acids 200-300 of human KLK2 |
| KLK3 | SCBT | sc-7316 | Mouse | Mono | Full length KLK3 of human origin |
| KLK14 | Abcam | Ab128957 | Rabbit | Mono | Synthetic peptide within the C-terminus of human KLK14 |
| Transferrin | Abcam | Ab66952 | Rabbit | Poly | Full length Transferrin of human origin |
| HSP90 | SCBT | sc-69703 | Mouse | Mono | Full length HSP90 of human origin |

**In gel-fluorescence**: LNCaP conditioned media (50 µg) was treated with a fluorescent activity-based probe for 1 h. After 1 h, proteins were precipitated using the aforementioned procedure and resuspended in a 3:1 ratio of PBS and sample loading buffer to a concentration of 1 mg/mL. After boiling for 5 min, 10 µL of sample was loaded on to a hand cast 12 % Bis-Tris gel and proteins were separated by SDS-PAGE as described above. Gels were then scanned for fluorescence at the appropriate wavelength.

**Immunoprecipitation**: LNCaP conditioned media (100 µg) was treated with the appropriate ABP for 1h. Proteins were precipitated using the $CHCl_3$/MeOH procedure outlined above and resuspended in RIPA buffer (20 mM Tris, 150 mM NaCl, 1 mM EDTA, 1mM EGTA, 1 % NP-40, 1 % sodium deoxycholate, 2.5 mM sodium pyrophosphate, 1 mM glycerophosphate, 1 mM $Na_3VO_4$ pH 7.5) to a concentration of 1 mg/mL. After reserving 10 µL of solution for the before pull-down sample, 1 µg of KLK, Transferrin or HSP90 antibody was added to the remaining protein followed by gentle agitation at RT for 2 h. After 2 h, 50 µL of protein G-coupled sepharose beads (Thermo Fisher Scientific) were added to the sample followed by gentle agitation at 4 °C for 12 h. After 12 h, the supernatant was removed (retaining 10 µL for the 'supernatant' sample) and the beads were washed with RIPA buffer (5 x 200 µL). 15 µL of PBS and 5 µL of sample loading buffer were added to the beads and the resulting mixture was boiled for 10 min. Samples were then processed as described above.

#### 2.4 Enzyme Assays

**Buffers**: KLK2 - 50 mM Tris, 10 mM $CaCl_2$, 150 mM NaCl, 0.05 % Brij-35, pH 7.5; KLK14 - 50 mM Tris, 150 mM NaCl, 0.05 % Brij-35, pH 8.0.

**Positional Scanning Substrate Library Screening**: Screening was carried out in the appropriate KLK buffer using 5 nM KLK2 or 1 nM KLK14. The final concentration used for each library member was 50 μM and the total reaction volume was 50 μL. The excitation wavelength was 355 nm, and the emission wavelength was 460 nm. An amount of 1 μL of each HyCoSuL peptide was added into a well in a 96-well plate containing 24 μL of KLK buffer. Next, 25 μL of the active KLK enzyme was added to each well and substrate hydrolysis was recorded in kinetic mode at RT for at least 10 min. The linear portion of each progress curve was used to calculate relative fluorescence units per second ($RFUs^{-1}$). All experiments were repeated at least three times and the results are presented as mean values ± SEM. Specificity was calculated by normalizing $RFUs^{-1}$ to the highest value, which was set at 100 %. Calculations were made in GraphPad Prism 5.

**Determination of $k_{cat}/K_M$**: The KLK2 substrate and KLK14 substrate were serially diluted in the appropriate KLK assay buffer and 25 μL was added to the wells of a 96-well plate. Next, 25 μL of KLK2 (1.5 nM) or KLK14 (75 pM) was added to each well containing the substrate and the enzyme-substrate reaction was monitored for at least 10 min. The linear portion of each progress curve was used to calculate relative fluorescence units per second ($RFUs^{-1}$). $K_M$, $k_{cat}$, and $k_{cat}/K_M$ parameters were determined by nonlinear regression in GraphPad Prism 5 (GraphPad Software). Each experiment was repeated three times and the results are presented as an average ± SEM.

**Determination of $k_{inact}/K_I$**: Under pseudo-first order conditions, a fixed amount of KLK2 (1.5 nM) or KLK14 (37 pM) was incubated with peptidyl-DPP analogue at different concentrations in the presence of 5 μM of substrate. The pseudo first-order rate constant, $k_{obs}$, was calculated for each probe concentration using non-linear regression analysis in GraphPad Prism 5. The obtained values were plotted against probe concentration to afford the apparent second order rate constant $k_{inact(app)}/K_I$. The absolute value of the second order rate constant was then obtained by adjusting for the substrate competition factor (1 + [S]/Km), where S is the concentration of substrate in the assay and $K_M$ is the Michaelis Menten constant for that substrate/enzyme reaction.

#### 2.5 Cell Migration Assays

**Cell migration**: The assay was performed as per the manufacturer's protocol (https://www.corning.com/catalog/cls/documents/protocols/an_Chemotaxis_protocol.pdf). Briefly, LNCaP-WT, LNCaP-mK14 and LNCaP-K14 cells were grown to 80% confluency in RPMI media supplemented with 5 % FBS. The media was then aspirated, the cells were washed with 2 mL PBS (3*) and incubated with serum-free RPMI (without phenol red) supplemented with 10 nM DHT ± 100 ng/mL Dox for 24 h. Cells were trypsinized, washed twice with serum-free RPMI and resuspended in serum-free RPMI. 100,000 cells in 100 μL of serum-free RPMI was then added to the transwell inserts placed in a 24-well plate (Corning

Cat. No. 3422) with receiver wells filled with 500 μL of FBS and incubated for 4 h. 1 μM of the appropriate ABP was then added and the cultures were incubated for a further 24 hours before medium was aspirated from the receiver plate and transwell inserts. Inserts were washed once and receiver wells twice with serum-free RPMI. Next, inserts were added back into the same receiver plate filled with 350 μL of 1 x cell dissociation solution and Calcein AM and incubated for 60 mins at 37 °C in a 5% $CO_2$ incubator with gentle tapping of the 24-well plate every 15 mins. Inserts were then removed from the 24-well plate and 100 μL of solution from each well was transferred to a black 96-well plate (in triplicate) and fluorescence was read with filters set to 485 nm for excitation and 520 nm for emission. In parallel, a standard curve was generated to enable the percentage of migrating cells to be assessed. Different cell numbers (3125-100,000) were incubated in 24-well receiver plates filled with 350 μL of 1 x cell dissociation solution and Calcein AM. After 60 mins at 37 °C in a 5% $CO_2$ incubator 100 μL of solution from each well was transferred to a black 96-well plate and fluorescence was read as above.

#### 3 Docking analysis with Molecular Operating Environment (MOE)

Molecular modelling studies were conducted using the software Molecular Operating Environment 2019.[3]

For KLK2 (PDB code 4NFE[4]) docking studies, solvent molecules as well as the extra benzamidine molecule lying outside the S1 pocket were deleted. The protein was then prepared using default parameters (quickprep) and the ligand site was used to perform a 'General Dock' of truncated compound **31** (lacking the biotin chain), which was imported as a sdf file from chemdraw, placing 50 poses with 'Triangle matcher', 'London' scoring, and refining 15 poses with 'Induced fit', 'GBVI/WSA' scoring. A pose (GBVI/WSA dG: –10.3020) correctly placing the amidine group in the S1 pocket was chosen and used to generate a pharmacophore query, which was used to perform a second 'General Dock' (50 placements, 15 pose refinements) on the obtained docked ligand following another protein preparation step (quickprep). One obtained pose (GBVI/WSA dG: –11.0790) was refined in 2 further rounds of 'General Dock' using the amidine pharmacophore and the setting "Ligand>Conformation>None". The final pose (GBVI/WSA dG: –12.0139) was minimized and exported as a PDB file. PyMOL was used to generate figure **S20B & D**.

For KLK3 (PDB code 2ZCK[5]) docking studies, chains H and L as well as solvent molecules were deleted. The protein was then prepared using default parameters (quickprep) and the ligand site was used to perform a 'General Dock' of compound **15**, which was imported as a sdf file from chemdraw, placing 200 poses with 'Triangle matcher', 'London' scoring, and refining 50 poses with 'Induced fit', 'GBVI/WSA' scoring. A pose (GBVI/WSA dG: –11.4271) correctly placing the P1 group in the S1 pocket was chosen and used to generate a pharmacophore query, which was used to perform a second 'General Dock' (50 placements, 15 pose refinements) on the obtained docked ligand following another protein preparation step (quickprep). One obtained pose (GBVI/WSA dG: –13.1373) was refined in 1 further round of 'General Dock' and the setting "Ligand>Conformation>None" (50 placements, 15 pose refinements) on the obtained docked ligand following another protein preparation step (quickprep). The final pose (GBVI/WSA dG: –12.9309) was minimized and exported as a PDB file. PyMOL was used to generate figure **S20A & C**.

### 4 Chemical Synthesis

Unless stated otherwise - all reactions were performed using anhydrous solvents, under an atmosphere of argon, monitored by thin layer chromatography (TLC) and allowed to stir at RT until completion.

#### 4.1 Equipment List

**Chromatography**: Thin Layer Chromatography was performed on Merck pre-coated Silica plates (Aluminum oxide 60 F254, Merck). Compounds were visualized by UV light (254 nm) or the appropriate stain (potassium permanganate, ninhydrin or *p*-anisaldehyde stain). Silica gel column chromatography was carried out using an Isolera (Biotage, UK) automated fraction collector apparatus equipped with SNAP cartridge columns (Biotage, UK).

**Analytical and Preparative LC-MS**: Analysis and purification of individual substrates and ABPs was carried out on a Waters RP-HPLC system consisting of (**i**) A Waters 2767 auto-sampler for sample injection and collection (**ii**) A Waters 515 HPLC pump for delivery of the mobile phase to the source (**iii**) X-Bridge C18 columns with dimensions 4.6 mm x 100 mm for analytical runs and 19 mm x 100 mm for preparative runs (**iv**) A Waters 3100 mass spectrometer with positive and negative mode ESI (**v**) A Waters 2998 Photodiode Array with detection between 200-600 nm. The following elution methods were used: **Method A** – a gradient of $H_2O$ and $CH_3CN$, supplemented with 0.1% formic acid: 0-10 min 5-98% $CH_3CN$, 10-12 min 98% $CH_3CN$, 12-13 min 98-5% $CH_3CN$, 13-18 min 5% $CH_3CN$. **Method B** – a gradient of $H_2O$ and $CH_3CN$, supplemented with 0.1% formic acid: 0-10 min 20-98% $CH_3CN$, 10-12 min 98% $CH_3CN$, 12-13 min 98 to 20% $CH_3CN$, 13-18 min 20% $CH_3CN$. A flow rate of 1.2 mL/min was used for analytical runs and 20 mL/min for preparative runs.

**Sample Drying**: Fractions obtained from preparative LC-MS purification were first concentrated using an EZ-2 series Genevac (SP Scientific, Pennsylvania, USA) and then combined and lyophilized using an Alpha 2-4 LD plus, Christ freeze-dryer (Osterode am Harz, Germany).

**NMR**: $^{1}H$, $^{13}C$, $^{19}F$ and $^{31}P$ NMR spectra were recorded on a Bruker AV-400 (400 Hz) instrument with TopSpin software using deuterated solvents for internal deuterium lock. Data is presented as follows: chemical shift, multiplicity (s = singlet, d = doublet, t = triplet, q = quartet, m = multiplet, b = broad), coupling constant(s) in Hz and integration. All NMR spectroscopy was carried out at RT unless otherwise indicated.

**Mass-Spectrometry**: High-resolution mass spectra were obtained by Dr Lisa Haigh, Imperial Mass spectrometry service using a Waters Premier instrument operating in either ES+ or ES- mode. M/Z values are reported in daltons (Da).

#### 4.2 Peptidic Diphenyl Phosphonate Synthesis

The synthesis of peptidyl DPP compounds consisted of three steps. **Step 1**: Solution-phase synthesis of diphenyl α-amino alkylphosphonate derivatives. **Step 2**: Solid-phase synthesis of N-terminal capped peptide specificity sequences. **Step 3**: Solution-phase coupling of the diphenyl α-amino alkylphosphonate warhead and the peptide specificity sequence, followed by TFA treatment to remove amino acid protecting groups.

**Diphenyl Phosphonate Warhead**

**Oleksyszyn reaction**. Benzyl carbamate (4 mmol, 1 eq.) was combined with triphenylphosphite (4 mmol, 1 eq.), the corresponding aldehyde (6 mmol, 1.5 eq.) and acetic acid (0.2 mL per mmol of aldehyde) and heated to 90 °C for 2 h. The reaction was monitored by TLC with completion confirmed by benzyl carbamate consumption. The reaction mixture was concentrated *in vacuo* and methanol was added to allow crystallization at -20 °C overnight. Crystals were collected by vacuum filtration, washed with cold MeOH and $Et_2O$ and dried. The resulting crystalline solids were of sufficient purity for use in subsequent reactions.

**Benzyl ((diphenoxyphosphoryl)(4-hydroxyphenyl)methyl)carbamate (S1)**

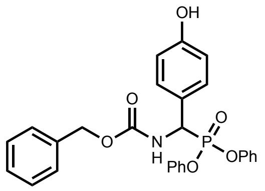

The title compound was isolated as a white solid in 52% yield. $^1$H NMR (400 MHz, DMSO-$d_6$) δ 9.58 (s, 1H), 8.80 (d, *J* = 10.2 Hz, 1H), 7.46 – 7.40 (m, 2H), 7.41 – 7.26 (m, 8H), 7.19 (m, 2H), 7.06 (d, *J* = 8.0 Hz, 2H), 6.96 (d, *J* = 8.1 Hz, 2H), 6.77 (dd, *J* = 8.1, 4.4 Hz, 2H), 5.47 (dd, *J* = 21.5, 10.0 Hz, 1H), 5.21 – 5.00 (m, 2H); $^{13}$C NMR (101 MHz, DMSO-$d_6$) δ 157.88, 156.47, 150.58, 150.39, 137.18, 130.29, 130.26, 129.84, 128.84, 128.42, 125.73, 125.64, 124.77, 120.82, 120.77, 119.27, 115.65, 66.70, 66.57; $^{31}$P NMR (162 MHz, DMSO-$d_6$) δ 15.31; HRMS (ESI, positive mode) found 490.1422 ($[C_{27}H_{24}NO_6P+H]^+$ requires 490.1420).

**Benzyl ((diphenoxyphosphoryl)(phenyl)methyl)carbamate (S3)**

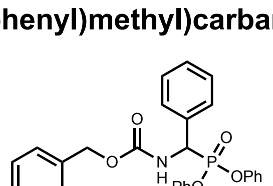

The title compound was isolated as a white solid in 80% yield. $^1$H NMR (400 MHz, $CDCl_3$) δ 7.56 – 7.48 (m, 2H), 7.43 – 7.32 (m, 8H), 7.32 – 7.25 (m, 2H), 7.25 – 7.17 (m, 3H), 7.12 (t, *J* = 7.7 Hz, 3H), 6.90 – 6.82 (m, 2H), 6.07 (dd, *J* = 10.2, 4.1 Hz, 1H), 5.62 (dd, *J* = 22.3, 10.0 Hz, 1H), 5.18 (d, *J* = 12.2 Hz, 1H), 5.08 (d, *J* = 12.2 Hz, 1H); $^{13}$C NMR (101 MHz, $CDCl_3$) δ 155.48, 150.04, 149.94, 135.91, 134.08, 129.77, 129.66, 129.53, 128.92, 128.70, 128.57, 128.32, 128.22, 125.44, 125.36, 120.51, 120.47, 120.36, 120.32, 115.36, 67.57, 53.62, 52.05.

$^{31}$P NMR (162 MHz, DMSO-$d_6$) δ 14.79; HRMS (ESI, positive mode) found 474.1470 ([$C_{27}H_{24}NO_5P$+H]$^+$ requires 474.1483).

**4-((((Benzyloxy)carbonyl)amino)(diphenoxyphosphoryl)methyl)benzoic acid (S5)**

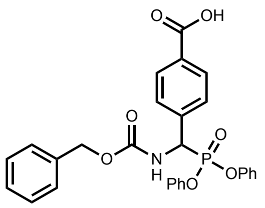

The title compound was isolated as a white solid in 57% yield. $^1$H NMR (400 MHz, DMSO-$d_6$) δ 9.04 (d, *J* = 10.0 Hz, 1H), 7.97 (d, *J* = 8.1 Hz, 2H), 7.78 (dd, *J* = 8.4, 2.1 Hz, 2H), 7.42 – 7.29 (m, 9H), 7.25 – 7.13 (m, 2H), 7.09 – 7.04 (m, 2H), 7.01 (d, *J* = 8.0 Hz, 2H), 5.73 (dd, *J* = 23.1, 10.1 Hz, 1H), 5.16 (d, *J* = 12.5 Hz, 1H), 5.07 (d, *J* = 12.4 Hz, 1H); $^{13}$C NMR (101 MHz, DMSO-$d_6$) δ 172.48, 167.40, 150.49, 139.72, 137.05, 131.06, 130.38, 130.31, 129.82, 129.13, 129.08, 128.84, 128.43, 125.89, 125.78, 120.80, 120.76, 120.73, 120.69, 115.67, 66.74, 52.45, 49.07; $^{31}$P NMR (162 MHz, DMSO-$d_6$) δ 13.99; HRMS (ESI, positive mode) found 518.1369 ([$C_{28}H_{24}NO_7P$+H]$^+$ requires 518.1328).

**Benzyl ((diphenoxyphosphoryl)(4-nitrophenyl)methyl)carbamate (S8)**

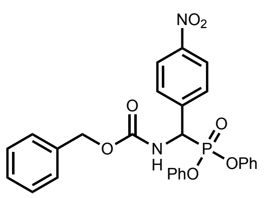

The title compound was isolated as a white solid in 58% yield. $^1$H NMR (400 MHz, DMSO-$d_6$) δ 9.13 (d, *J* = 9.9 Hz, 1H), 8.29 (d, *J* = 8.4 Hz, 2H), 7.96 (d, *J* = 8.1 Hz, 2H), 7.50 – 7.27 (m, 9H), 7.22 (dt, *J* = 10.3, 5.2 Hz, 2H), 7.06 (dd, *J* = 13.0, 8.1 Hz, 4H), 5.89 (dd, *J* = 23.7, 10.1 Hz, 1H), 5.17 (d, *J* = 12.5 Hz, 1H), 5.08 (d, *J* = 12.4 Hz, 1H); $^{13}$C NMR (101 MHz, DMSO-$d_6$) δ 156.51, 150.46, 150.17, 147.83, 142.58, 137.00, 130.43, 130.35, 130.23, 130.18, 128.87, 128.48, 125.99, 125.88, 124.03, 120.82, 120.70, 66.85, 53.82, 52.26, 40.64, 40.43, 40.22, 40.01, 39.80, 39.59, 39.39; $^{31}$P NMR (162 MHz, DMSO-$d_6$) δ 13.24; HRMS (ESI, negative mode) 517.1171 ([$C_{27}H_{23}N_2O_7P$-H]$^-$ requires 517.1165).

**Benzyl ((4-carbamoylphenyl)(diphenoxyphosphoryl)methyl)carbamate (S6)**

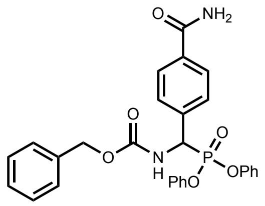

Compound **S5** (0.50 g, 0.97 mmol, 1 eq.) and di-tert-butyldicarbonate (0.63 g, 2.91 mmol, 3 eq.) were suspended in 12 mL THF/ 1mL pyridine and stirred at 50 °C for 30 min. Ammonium carbonate (559 mg, 5.82 mmol, 6 eq.) was added to the reaction mixture and the solution was stirred for an additional 6 h under the same conditions. The reaction was monitored by TLC and after completion solvents were removed under rotatory evaporation and the solid residue was suspended in MeOH. The solution was stirred with gentle heating until dissolution to remove excess ammonia. The reaction vessel was then cooled to -20 °C and the resulting white crystalline precipitate was filtered and dried to afford compound **S6** in 56 % yield. $^{1}$H NMR (400 MHz, DMSO- $d_6$) δ 8.97 (d, *J* = 10.1 Hz, 1H), 8.00 (s, 1H), 7.89 (d, *J* = 8.1 Hz, 2H), 7.72 (dd, *J* = 8.4, 2.1 Hz, 2H), 7.45 – 7.27 (m, 10H), 7.26 – 7.16 (m, 2H), 7.09 – 7.03 (m, 2H), 7.00 (d, *J* = 8.0 Hz, 2H), 5.69 (dd, *J* = 22.8, 10.1 Hz, 1H), 5.15 (d, *J* = 12.5 Hz, 1H), 5.07 (d, *J* = 12.4 Hz, 1H); $^{13}$C NMR (101 MHz, DMSO-$d_6$) δ 167.90, 156.57, 137.92, 137.07, 133.86, 132.41, 130.36, 130.30, 128.84, 128.42, 128.00, 125.87, 125.76, 120.82, 120.78, 120.73, 120.69, 118.79, 117.42, 66.71, 53.92, 50.81, 43.84, 40.62; $^{31}$P NMR (162 MHz, DMSO-$d_6$) δ 14.23; HRMS (ESI, positive mode) found 517.1523 ($[C_{28}H_{25}N_2O_6P+H]^+$ requires 517.1529).

**Benzyl((diphenoxyphosphoryl)(4-(N, N' di-Boc)guanidinophenyl)methyl)carbamate (S9)**

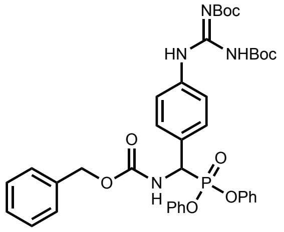

Acetic acid (4 mL mmol$^{-1}$ **S8**) was added to a mixture of **S8** (5.00 g, 9.63 mmol, 1 eq.) and Fe powder (4.84 g, 86.7 mmol, 9 eq.). The reaction mixture was heated to 70 °C for 2 h. The acetic acid was then removed, and the crude residue dissolved in EtOAc. The sample was centrifuged for 5 min at 3000 rpm to remove the $Fe_2O_3$ by-product and the supernatant was concentrated. N,N'-bis-Boc-1-guanylpyrazole (2.98 g, 9.63 mmol, 1eq.) and triethylamine (2.68 mL, 19.26 mmol, 2 eq.) were added to the crude solid (4.71 g, 9.63 mmol, 1 eq.) in DCM (50 mL). The solution was allowed to stir overnight at RT. Solvent was removed by rotatory evaporation and the resulting residue was dissolved in EtOAc and washed with 1M HCl, sat. $NaHCO_3$ and Brine. The organic layer was dried over anhydrous $MgSO_4$, filtered and concentrated *in vacuo*. The crude oil was purified by flash silica chromatography using a gradient of 10%-100% EtOAc in hexane. After concentration of pure fractions **S9** was obtained as a white solid in 47% yield. $^{1}$H NMR (400 MHz, DMSO-$d_6$) δ 11.41 (s, 1H), 10.03 (s, 1H), 8.90 (d, *J* = 10.1 Hz, 1H), 7.65 – 7.54 (m, 4H), 7.41 – 7.30 (m, 9H), 7.21 (t, *J* = 7.4 Hz, 2H), 7.06 (d, *J* = 8.1 Hz, 2H), 6.99 (d, *J* = 8.2 Hz, 2H), 5.60 (dd, *J* = 22.2, 10.1 Hz, 1H), 5.15 (d, *J* = 12.5 Hz, 1H), 5.07 (d, *J* = 12.5 Hz, 1H), 1.52 (s, 9H), 1.41 (s, 9H); $^{13}$C NMR (101 MHz, DMSO-$d_6$) δ 170.98, 163.10, 156.70, 156.49, 153.28, 152.56, 151.13, 150.31, 137.11, 131.16, 130.33, 130.26, 129.33, 128.82, 128.42, 125.78, 125.68, 123.14, 120.86, 120.75, 83.86, 82.31, 80.92, 79.35, 77.49, 66.68, 40.60, 40.39, 40.18, 39.97, 39.77, 39.56, 39.35, 28.67,

28.33, 28.20, 28.10, 24.72. $^{31}P$ NMR (162 MHz, DMSO-$d_6$) δ 14.67; HRMS (ESI, positive mode) found 731.1806 ($[C_{38}H_{43}N_4O_9P+H]^+$ requires 731.1797).

**Cbz Deprotection.** Cbz-protected DPP compounds (1 mmol) were treated with 33% HBr-AcOH solution (5 mL) for 2 h at RT. The solvent was removed by rotatory evaporation and the residue was dissolved in a minimal amount of MeOH. An excess of diethyl ether was added and overnight storage at -20 °C led to crystallization. Crystalline solids were filtered, washed with cold diethyl ether and dried to afford α-amino DPP products as HBr salts. Compound purity was sufficient for use in subsequent reactions.

**Diphenyl (amino(4-hydroxyphenyl)methyl)phosphonate (S2)**

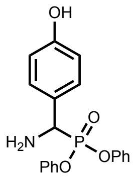

The title compound was isolated as a white solid in 94% yield. $^1H$ NMR (400 MHz, MeOD) δ 7.74 (m, 2H), 7.45 – 7.38 (m, 2H), 7.35 – 7.26 (m, 5H), 7.25 – 7.11 (m, 2H), 6.88 (dt, $J$ = 8.2, 1.2 Hz, 2H), 5.54 (d, $J$ = 18.4 Hz, 1H); $^{31}P$ NMR (162 MHz, MeOD); δ 13.44; LCMS – 18 min, 5 to 98% $CH_3CN$ in $H_2O$, Rt 8.95 min; HRMS (ESI, positive mode) found 356.1039 ($[C_{19}H_{18}NO_4P+H]^+$ requires 356.1052).

**Diphenyl (amino(phenyl)methyl)phosphonate (S4)**

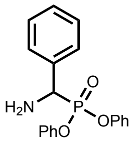

The title compound was isolated as a white solid in 98% yield. $^1H$ NMR (400 MHz, MeOD) δ 7.73 – 7.66 (m, 2H), 7.59 – 7.53 (m, 3H), 7.43 – 7.34 (m, 2H), 7.32 – 7.24 (m, 3H), 7.23 – 7.16 (m, 1H), 7.12 (m, 2H), 6.87 – 6.77 (m, 2H), 5.49 (d, $J$ = 18.3 Hz, 1H); $^{13}C$ NMR (101 MHz, MeOD) δ 149.94, 149.77, 130.62, 130.55, 130.40, 130.10, 129.52, 129.46, 129.43, 126.33, 126.21, 120.91, 120.87, 120.73, 120.70, 51.67; $^{31}P$ NMR (162 MHz, MeOD) δ 10.22; LCMS – 18 min, 5 to 98% $CH_3CN$ in $H_2O$, Rt 9.38 min; HRMS (ESI, positive mode) found 340.1116 ($[C_{19}H_{18}NO_3P+H]^+$ requires 340.1103).

**Diphenyl (amino(4-carbamoylphenyl)methyl)phosphonate (S7)**

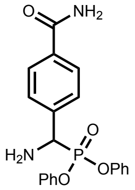

The title compound was isolated as a white solid in 87% yield. $^1H$ NMR (400 MHz, DMSO-$d_6$) δ 9.53 (s, 2H), 8.10 (d, $J$ = 4.1 Hz, 1H), 8.00 (d, $J$ = 7.9 Hz, 2H), 7.78 (m, 2H), 7.53 (s, 1H),

7.39 (m, 4H), 7.24 (m, 2H), 7.13 (d, $J$ = 7.9 Hz, 2H), 6.97 (m, 2H), 5.89 – 5.68 (m, 1H); $^{13}$C NMR (101 MHz, DMSO-$d_6$) δ 167.61, 149.80, 135.74, 133.59, 130.59, 130.46, 129.27, 128.43, 126.40, 126.30, 120.89, 120.84, 120.72, 51.47, 49.93; $^{31}$P NMR (162 MHz, DMSO-$d_6$) δ 11.06; LCMS – 18 min, 5 to 98% $CH_3CN$ in $H_2O$, Rt 8.53 min; HRMS (ESI, positive mode) found 383.1147 ($[C_{20}H_{19}N_2O_4P+H]^+$ requires 383.1161).

**Diphenyl (amino(4-guanidinophenyl)methyl)phosphonate (S10)**

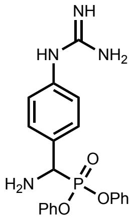

The title compound was isolated as a white solid in 94% yield. $^1$H NMR (400 MHz, DMSO-$d_6$) δ 9.90 (s, 1H), 9.47 (s, 3H), 7.74 (d, $J$ = 8.1 Hz, 2H), 7.57 (s, 4H), 7.47 – 7.32 (m, 5H), 7.26 (m, 2H), 7.15 (d, $J$ = 8.0 Hz, 2H), 7.03 (d, $J$ = 8.0 Hz, 2H), 5.74 (d, $J$ = 18.4 Hz, 1H); $^{13}$C NMR (101 MHz, DMSO); δ 155.82, 149.91, 149.76, 137.18, 130.89, 130.84, 130.55, 130.45, 127.91, 127.87, 126.33, 126.26, 123.76, 120.92, 49.04; $^{31}$P NMR (162 MHz, DMSO) δ 11.21; LCMS – 18 min, 5 to 98% $CH_3CN$ in $H_2O$, Rt 7.68 min; HRMS (ESI, positive mode) found 397.1415 ($[C_{20}H_{21}N_4O_3P+H]^+$ requires 397.1430).

#### Peptide Specificity Sequence

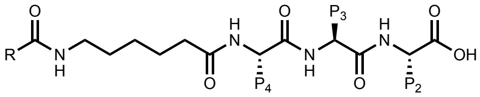

Specificity sequences were obtained by use of solid-phase peptide synthesis. 2-chlorotrityl chloride resin (100 mg, loading 1.31 mmol/g) was suspended in DCM and shaken for 10 min, followed by washing (3 x dry DCM). Fmoc-P2-OH (0.393 mmol, 3 eq.) and DIPEA (136 µL, 0.786 mmol, 6eq.) in dry DCM were then added to the resin and the mixture was agitated for 3 h at RT. After 3 h the resin was filtered and washed with 1 mL DCM (3 x) before addition of 2 mL MeOH and agitation for a further 30 min. The resin was again filtered and washed with 1 mL DMF (3 x), DCM (3 x) and DMF (3 x). At this stage it was assumed that the yield of the Fmoc-P2-OH amino acid coupling step was 100 % and thus a loading amount of 1.31 mmol/g was used to calculate molar equivalents for all remaining couplings. To remove the Fmoc protecting group the resin was treated with 20% PIP in DMF (3 x 2 mL) for 3 min per treatment. After the third treatment the resin was washed with 1 mL DMF (3 x), DCM (3 x) and DMF (3 x). Next, Fmoc-P3-OH (0.328 mmol, 2.5 eq) was pre-activated with HBTU (124.0 mg, 0.328 mmol, 2.5 eq.) and DIPEA (57.00 µL, 0.328 mmol, 2.5 eq.) in DMF for 3 min and added to the resin, which was shaken for 1 h at RT. To check that the coupling had proceeded to completion a 'ninhydrin test' was used. To a glass vial containing several resin beads was added 1 mL of ninhydrin solution (5 g of ninhydrin in 100 mL EtOH). The mixture was then heated for 3 min at 95 °C. A positive test result (free amine groups, incomplete reaction) is indicated by a dark blue color of resin, while a negative test (no free amine groups, completed reaction) is indicated by the resin remaining colorless/pale yellow. Following reaction completion, the resin was washed with 1 mL DMF (3 x), DCM (3 x) and DMF (3 x). After Fmoc removal, the peptide chain was elongated with Fmoc-P4-OH and Fmoc-Ahx-OH using the same procedure outlined above. Finally, following Fmoc deprotection the peptide chain was capped on the N-terminus with either 4-pentynoic acid, morpholin-4-yl acetic acid or biotin using the same procedure as for the P2-P4 amino acid couplings. Following reaction completion, the resin was washed with 1 mL DMF (3 x), DCM (3 x), MeOH (3 x) and diethyl ether (3 x) and dried over $P_2O_5$ for at least 3 h. The peptide was then cleaved from the 2-chlorotrityl chloride resin by treatment with 1 mL of a 1:3 mixture of HFIP:DCM for 2 h at RT. Filtrate was collected and the peptide was precipitated by addition of 13 mL of cold diethyl ether and storage at -20 °C for 30 min. The sample was then centrifuged (5 min, 3000 rpm), the supernatant decanted and the solid dissolved in 3 mL of a 2:1 mixture of $CH_3CN$:$H_2O$. Crude peptide products were lyophilized and used for coupling to DPP warheads without further purification.

**Peptide Warhead Coupling**

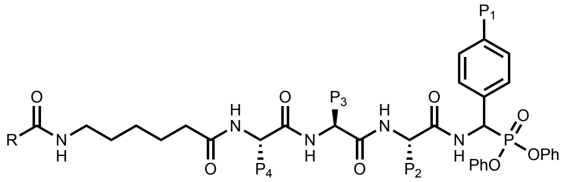

Crude peptide (0.13 mmol, 1 eq.) was dissolved in 1 mL of DMF and activated with HATU (50.0 mg, 0.13 mmol, 1 eq.) and 2,4,6-trimethylpyridine (86.0 µL, 0.66 mmol, 5 eq.) for 3 min before being added to a round-bottom flask (RBF) containing the appropriate α-amino DPP warhead (0.16 mmol, 1.2 eq.). The mixture was stirred at RT and reaction progress was monitored by LCMS (Method B). Typically, the reaction was complete after 2 h. After completion DMF was removed under a stream of $N_2$ gas and the resulting residue was treated with 1 mL of TFA deprotection mixture (95 % TFA, 2.5 % TIS and 2.5 % $H_2O$) for 2 h at RT. Volatiles were removed under a stream of $N_2$ gas and the residue was dissolved in DMSO:$CH_3CN$:$H_2O$ (1:4.5:4.5) and purified by preparative LCMS. Pure fractions were combined and lyophilized to afford peptidyl DPP compounds.

#### 4.3 Fluorescent Diphenyl Phosphonate Activity-based Probes

Alkyne-DPP (5 µmol, 1 eq.) and Fluorophore-azide (5 µmol, 1 eq.) were dissolved in 800 µL of DMF and the solution was deoxygenated for 10 min with a flow of argon gas. Next, sodium ascorbate (0.74 mg, 3.75 µmol, 0.75 eq.) in 100 µL of $H_2O$ and $CuSO_4.5H_2O$ (0.62 mg, 2.5

μmol, 0.5 eq.) in 100 μL were added to the reaction vessel and the reaction mixture was stirred overnight at RT under an argon atmosphere. DMF was removed under a flow of $N_2$ gas, and the resulting residue was dissolved in DMSO:$CH_3CN$:$H_2O$ (1:4.5:4.5) and purified by preparative LCMS. Pure fractions were combined and lyophilized.

**TAMRA-SQL-Ty'-DPP (3)**

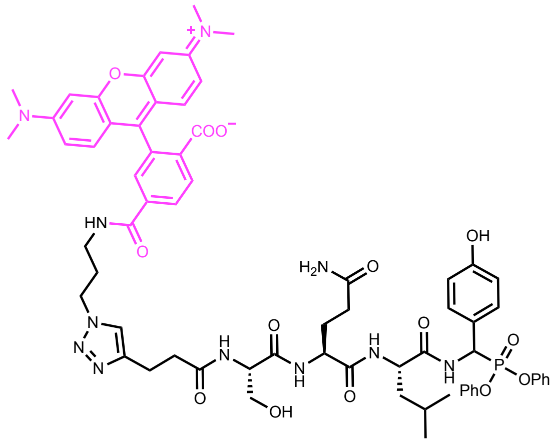

The title compound was isolated as a pink solid in 48 % yield. LCMS – 18 min, 20 to 98% $CH_3CN$ in $H_2O$, Rt 7.10 & 7.30 min; HRMS (ESI, positive mode) found 1276.5201 ($[C_{66}H_{74}N_{11}O_{14}P+H]^+$ requires 1276.5233).

**TAMRA-NQ-NVa-Gln'-DPP (20, KLK3_fABP)**

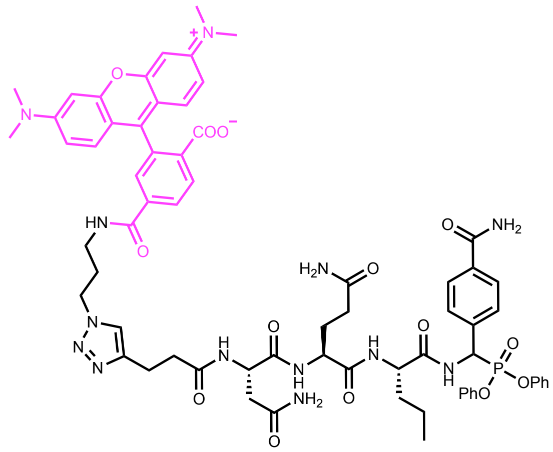

The title compound was isolated as a pink solid in 41 % yield. LCMS – 18 min, 20 to 98% $CH_3CN$ in $H_2O$, Rt 8.05 & 8.15 min; HRMS (ESI, positive mode) found 1316.5315 ($[C_{67}H_{74}N_{13}O_{14}P+H]^+$ requires 1316.5205).

**Cy5-Tle-Dab-Y-Arg'-DPP (42, KLK2_fABP)**

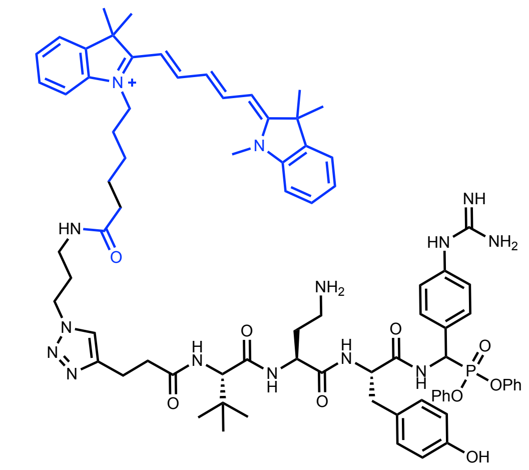

The title compound was isolated as a blue solid in 57 % yield. LCMS – 18 min, 20 to 98% $CH_3CN$ in $H_2O$, Rt 6.25 & 6.55 min; HRMS (ESI, positive mode) found 1417.7391 ($[C_{79}H_{98}N_{14}O_9P+H]^+$ requires 1417.7233).

**Fluorescein-2Nal-Dab-Abu-Arg'-DPP (43, KLK14_fABP)**

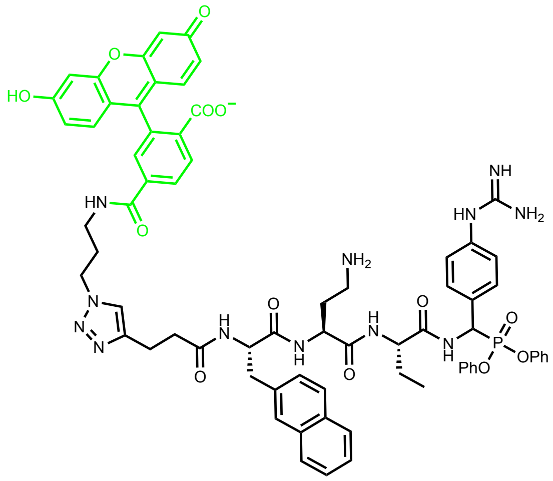

The title compound was isolated as an orange solid in 32 % yield. LCMS – 18 min, 20 to 98% $CH_3CN$ in $H_2O$, Rt 8.55 & 8.90 min; HRMS (ESI, positive mode) found 1314.4706 ($[C_{70}H_{68}N_{12}O_{13}P-H]^-$ requires 1314.4837).

#### 4.4 Positional Scanning Substrate Library Synthesis

The synthesis of the library was divided into five key parts. **Part 1**: Large-scale synthesis of the coumarin **S14**. For each sub-library 7 g of material was required and thus synthesis was carried out on a scale that would provide at least 25 g of pure product. **Part 2:** Synthesis of Arginine-ACC resin **S17**. For each sub-library 12 g of rink amide resin was used. **Part 3-5:** Synthesis of individual sub-libraries.

#### Fmoc-ACC-OH

##### Ethyl (3-hydroxyphenyl)carbamate (S11)

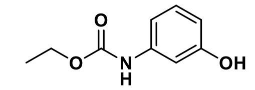

To a 2 L RBF was added 3-aminophenol (150 g, 1.37 mol, 2 eq.) and EtOAc (500 mL). The mixture was heated under reflux for 30 min before dropwise addition of ethyl chloroformate (74.6 g, 0.687 mol, 1 eq.) over a 1 h period. The reaction mixture was allowed to cool to RT and the resulting precipitate was filtered and washed with EtOAc (3 x 300 mL) and hexane (3 x 300 mL). The combined filtrate was concentrated to afford 123 g of **S11** as a white solid. Yield = 99%. The product was sufficiently pure to use in subsequent reactions. $^{1}$H NMR (400 MHz, DMSO-$d_6$) δ 9.49 (s, 1H), 9.33 (s, 1H), 7.03-6.98 (m, 2H), 6.86-6.79 (m, 1H), 6.39-6.35 (m, 1H), 4.11 (q, $J$ = 7.1 Hz, 2H), 1.24 (t, 7.1 Hz, 3H); $^{13}$C NMR (101 MHz, DMSO-$d_6$) δ 158.14, 153.88, 140.75, 129.79, 109.88, 109.44, 105.75, 60.46, 15.00.

##### 2-(7-((ethoxycarbonyl)amino)-2-oxo-2H-chromen-4-yl)acetic acid (S12)

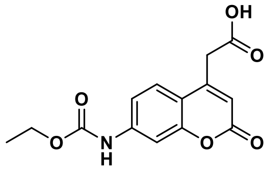

To a 5 L RBF were added S11 (100 g, 0.55 mol, 1eq.) and 70% H2SO4 (2.5 L). The mixture was cooled in an ice bath and vigorously stirred before portion-wise addition of 1,3-acetonedicarboxylic acid (88.7 g, 0.61 mol, 1.1 eq.). The reaction was allowed to warm to RT and was stirred for an additional 8 h. After 8 h, the reaction mixture was poured onto ice (4 kg) and stirred for 30 min. The resulting white precipitate was filtered and washed with diethyl ether (3 x 1 L). Crude material was suspended in hot CH3CN (700 mL) and the precipitate filtered to afford 90 g of S12 as a white solid. Yield = 56 %. 1H NMR (400 MHz, DMSO-d6) δ 10.18 (s, 1H), 7.63 (dd, J = 8.7, 1.9 Hz, 1H), 7.58 (d, J = 2.0 Hz, 1H), 7.39-7.35 (m, 1H), 6.34 (s, 1H), 4.18 (q, J = 7.1 Hz, 2H), 3.87 (s, 2H), 2.09 (d, J = 1.3 Hz, 1H), 1.27 (t, J = 7.7, 3H); 13C NMR (101 MHz, DMSO-d6) δ 171.08, 160.48, 154.50, 153.80, 150.36, 143.36, 126.60, 114.78, 114.10, 104.94, 61.21, 37.49, 31.17, 14.86; HRMS (ESI, positive mode) found 292.0824 ([C14H13NO6+H]+ requires 292.0821).

##### 2-(7-((ethoxycarbonyl)amino)-2-oxo-2H-chromen-4-yl)acetic acid (S13)

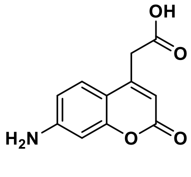

To a 2 L RBF was added **S12** (90.0 g, 0.31 mol, 1 eq.), NaOH (123g, 3.10 mol, 10 eq.) and $H_20$ (800 mL). The reaction mixture was stirred at reflux for 16 h. After cooling to RT, the pH of the reaction was adjusted to 2 by dropwise addition of $H_2SO_4$. The resulting precipitate was filtered and washed with diethyl ether (3 x 200 mL) to afford 45 g of **S13** as a yellow solid. Yield = 66 %. The product was sufficiently pure by NMR analysis to be used in subsequent steps without further purification. $^1$H NMR (400 MHz, DMSO-$d_6$) δ 7.34 (d, $J$ = 8.7 Hz, 1H), 6.56 (dd, $J$ = 8.7, 2.3 Hz, 1H), 6.43 (d, $J$ = 2.1 Hz, 1H), 6.17 (s, 2H), 5.99 (s, 1H), 3.74 (s, 2H); $^{13}$C NMR (101 MHz, DMSO-$d_6$) δ 171.29, 161.16, 156.16, 153.60, 150.84, 126.76, 111.74, 109.43, 108.66, 99.02, 37.73; HRMS (ESI, positive mode) found 220.0609 ($[C_{11}H_9NO_4+H]^+$ requires 220.0610).

**2-(7-((((9H-fluoren-9-yl)methoxy)carbonyl)amino)-2-oxo-2H-chromen-4-yl)acetic acid (S14)**

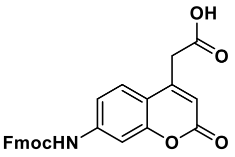

To a vigorously stirring suspension of **S13** (20.0 g, 91.2 mmol) in DCM (150 mL) was added TMSCl (21.8 g, 201 mmol, 2.2 eq.) and DIPEA (25.9 g, 201 mmol, 2.2 eq.). The reaction mixture was heated to reflux for 3 h, followed by cooling in an ice bath. Fmoc-Cl (26.0 g, 100 mmol, 1.1 eq.) was added portion wise and the reaction was allowed to warm to RT and stirred for overnight. Next, the reaction was stirred vigorously and MeOH (500 mL) was added. The resulting precipitate was collected by filtration and washed with MeOH (2 x 250 mL) and diethyl ether (2 x 250 mL) to afford 33.1 g of **S14** as an off-white solid. Yield = 82.3 %. $^1$H NMR (400 MHz, DMSO-$d_6$) δ 12.84 (s, 1H), 10.22 (s, 1H), 7.92 (d, $J$ = 7.4 Hz, 2H), 7.77 (d, $J$ = 7.5 Hz, 2H), 7.63 (d, $J$ = 8.7 Hz, 1H), 7.56 (s, 1H), 7.44-7.33 (m, 5H), 6.35 (s, 1H), 4.58 (d, $J$ = 6.4 Hz, 2H), 4.35 (t, $J$ = 6.4 Hz, 1H), 3.87 (s, 2H); $^{13}$C NMR (101 MHz, DMSO-$d_6$) δ 171.08, 160.45, 154.46, 153.70, 150.32, 144.12, 143.13, 141.31, 128.20, 127.63, 126.59, 125.56, 120.70, 114.91, 114.23, 105.16, 66.41, 47.02, 37.52; HRMS (ESI, positive mode) found 442.1259 ($[C_{26}H_{19}NO_6+H]^+$ requires 442.1291).

**NH2-Arg(Pbf)-ACC-Rink Amide Resin (S17)**

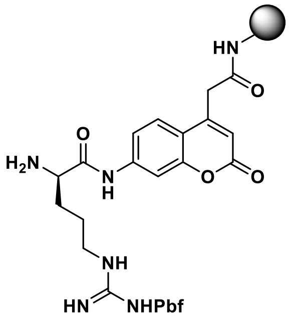

The Rink amide resin (12.0 g, 5.76 mmol, 1 eq.) was charged in an RBF with 100 mL of DCM and agitated for 30 min. After 30 min the resin was filtered and washed with DMF (3 x 60 mL). Next, the Fmoc-protecting group was removed from the resin by treatment with 20% PIP in DMF (3 x 80 mL) for 5 min, 5 min and 25 min. The resin was then washed with 60 mL of DMF (3 x), DCM (3 x) and DMF (3 x). At this stage a ninhydrin test was performed to confirm the Fmoc deprotection reaction had proceeded to completion. Next 6.40 g of **S14** (14.4 mmol, 2.5 eq.) and 2.16 g of HOBt (14.4 mmol, 2.5 eq.) were added to a falcon tube and dissolved in a minimal amount of DMF. Then, 1.90 mL of DICI (14.4 mmol, 2.5 eq.) was added and the mixture was gently shaken for 5 min. After 5 min the mixture was poured onto the resin and shaken for 24 h at RT. The resin was then filtered and washed with 60 ml of DMF (3 x), DCM (3 x) and DMF (3 x). A ninhydrin test was performed to check that the reaction had proceeded to completion. The Fmoc group was then removed using the same procedure described previously. Fmoc-Arg(Pbf)-OH (9.33 g, 14.4 mmol, 2.5 eq.) and HATU (5.47 g, 14.4 mmol, 2.5 eq.) were dissolved in a minimal amount of DMF before addition of 2,4,6-trimethylpyridine (1.90 mL, 14.4 mmol, 2.5 eq.) and gentle agitation for 3 min. The mixture was then poured on the resin and the vessel was shaken for 24 h at RT. The resin was filtered and washed with 60 mL of DMF (3 x). The coupling was then repeated using half the amount of reagents. Next, 3-nitro-1,2,4-triazole (6.57 g, 57.6 mmol, 10 eq.), AcOH (3.29 mL, 57.6 mmol, 10 eq.) and DICI (8.90 mL, 57.6 mmol, 10 eq.) were dissolved in DMF (40 mL) and gently shaken for 5 min. The reaction mixture was poured onto the resin and the vessel was agitated for 12 h at RT. After 12 h, DIPEA (2 mL, 11.52 mmol, 2 eq.) was added and the vessel was shaken for an additional 3 h. The resin was filtered and washed with 60 mL of DMF (3 x), DCM (3 x) and DMF (3 x). Finally, the Fmoc protecting group was removed from Arg by using the same protocol outlined previously. The resin was washed with 60 mL of DMF (3 x), DCM (3 x), DMF (3 x), MeOH (3x) and diethyl ether (3 x) and dried over $P_2O_5$ for at least 3 h.

#### P2 Sublibrary

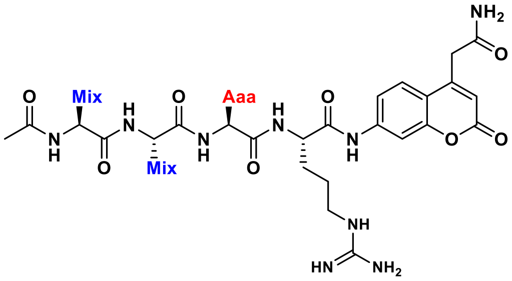

**General.** Each library consists of 105 peptides. Each of the natural amino acids (excluding methionine and cysteine, including norleucine) and 86 unnatural amino acids were coupled at the P2 position and an isokinetic mixture of 19 amino acids (excluding cysteine and methionine but including norleucine) was coupled at P3 and P4. Equivalent ratios of amino acids in the isokinetic mixture were created based on previously reported coupling rates (**Table S2**). For coupling of the various amino acids in the P2 position a 2.5 equimolar excess of amino acid was used. For coupling of the isokinetic mixture in P3 and P4 a 5-fold excess of mixture was used. All reactions were carried out using DIC and HOBt as the coupling reagents.

**Table S2:** Composition of the isokinetic mixture used during synthesis of the scanning library

| Amino acid | mol % | Amino acid | mol % |
|---|---|---|---|
| Fmoc-Ala-OH | 3.4 | Fmoc-Lys(Boc)-OH | 6.2 |
| Fmoc-Arg(Pbf)-OH | 6.5 | Fmoc-Nle-OH | 3.8 |
| Fmoc-Asn(Trt)-OH | 5.3 | Fmoc-Phe-OH | 2.5 |
| Fmoc-Asp(tBu)-OH | 3.5 | Fmoc-Pro-OH | 4.3 |
| Fmoc-Glu(tBu)-OH | 3.6 | Fmoc-Ser(tBu)-OH | 2.8 |
| Fmoc-Gln(Trt)-OH | 5.3 | Fmoc-Thr(tBu)-OH | 4.8 |
| Fmoc-Gly-OH | 2.9 | Fmoc-Trp(Boc)-OH | 3.8 |
| Fmoc-His(Trt)-OH | 3.5 | Fmoc-Tyr(tBu)-OH | 4.1 |
| Fmoc-Ile-OH | 17.4 | Fmoc-Val-OH | 11.3 |
| Fmoc-Leu-OH | 4.9 | | |

**P2 Coupling.** To 105 individual peptide syringe filters was added 1 eq. of **S17** (0.04 mmol, 80.0 mg). Next 1 mL of DCM was added to each syringe and the resin was gently agitated for 1 h. The resin was filtered and washed with 1 mL DMF (4 x). In separate 1.5 mL Eppendorf tubes, 2.5 eq. of Fmoc-P2-OH (0.1 mmol) was pre-activated with 2.5 eq. HOBt (0.1 mmol, 15 mg) and 2.5 eq. DICI (0.1 mmol, 14 µL) in 1 mL DMF. The pre-activated mixture was added to the resin followed by 3 h of agitation. After 3 h, the resin was filtered and washed with 1 mL of DMF (3 x), DCM (3 x) and DMF (3 x). A ninhydrin test was then carried out on all 105 resin mixtures to confirm that each reaction had proceeded to completion. Next, the Fmoc protecting group was removed by shaking in 1 mL of 20 % PIP in DMF (2 x 5 min and 1 x 25 min).

**P3 and P4 Coupling.** An isokinetic mixture was prepared in the amount required for 105 individual coupling reactions. 525 eq. of isokinetic mixture (22.2 mmol), 525 eq. HOBt (22.2 mmol, 3.00 g) and 525 eq. DICI (22.2 mmol, 3.43 mL) were dissolved in DMF up to a volume of 105 mL and pre-activated for 3 min. To each of the 105 syringes was added 1 mL of activated isokinetic mixture. After 3h agitation, the resin was filtered and washed with DMF (3 x), DCM (3 x) and DMF (3x). A ninhydrin test was performed on all 105 batches of resin to confirm that each reaction had proceeded to completion. The Fmoc protecting group was removed by shaking in 1 mL of 20 % PIP in DMF (2 x 5 min and 1 x 25 min). The exact same procedure was then carried out for coupling of the isokinetic mixture in the P4 position.

**N-terminus Acetylation.** An 'acetylation mixture' was prepared in the amount required for 105 individual capping reactions. 525 eq. AcOH (22.2 mmol, 1.27 mL), 525 eq. HBTU (22.2 mmol, 8.41 g) and 525 eq. DIPEA (22.2 mmol, 3.86 mL) were dissolved in DMF up to a volume of 105 mL and left to pre-activate for 3 min. 1 mL of acetylation mixture was added to each syringe and the suspension was agitated for 1 h. After 1 h, the resin was washed with 1 mL DMF (3 x), DCM (3 x), DMF (3 x), MeOH (3 x) and diethyl ether (3 x) and dried over $P_2O_5$ for at least 3 h.

**Cleavage from the resin.** Peptides were cleaved from the resin by addition of 2 mL of TFA deprotection mixture and gentle shaking for 2 h. The solution from each syringe was collected separately in individual 15 mL Falcon tubes and the resin was washed once with 1 mL of TFA deprotection mixture. To the 3 mL of deprotection mixture was added 12 mL of cold diethyl ether. After precipitation (1 h, - 20 °C) each mixture was centrifuged (5 min, 3000 rpm), washed with cold diethyl ether (4 mL), centrifuged again and left to dry at RT for 3 h. The resulting off-white precipitates were dissolved in 3 mL of a 2:1 ratio of $CH_3CN$:$H_2O$ and lyophilized. The final products were dissolved in DMSO to a concentration of 10 mM and used without further purification.

**P3 and P4 Sublibraries**

In the same manner as described for the P2 sublibrary, the P3 and P4 sub-libraries were synthesized by coupling fixed amino acid residues to the P3 position (isokinetic mixture to P2 and P4) and the P4 position (isokinetic mixture to P2 and P3), respectively.

#### 4.5 KLK Substrates

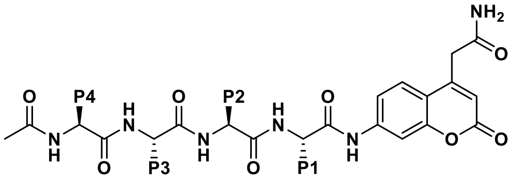

2.5 eq. Fmoc-P2-OH (0.1 mmol) was pre-activated with 2.5 eq. HOBt (0.1 mmol, 15 mg) and 2.5 eq. DICI (0.1 mmol, 14 µL) in 1 mL DMF and added to a syringe containing 1 eq. of **S17** (0.04 mmol, 80 mg), followed by gentle agitation for 1 h. A ninhydrin test was carried out after each coupling to check for reaction completion. The resin was filtered and washed with 1 mL of DMF (3 x), DCM (3 x) and DMF (3 x). The Fmoc-protecting group was removed using 20% PIP in DMF (3 x 2 mL) for 3 min per treatment. After the third treatment the resin was washed with 1 mL of DMF (3 x), DCM (3 x) and DMF (3 x). The same procedure was carried out for coupling of Fmoc-P3-OH and Fmoc-P4-OH. The N-terminus was then acetyl protected with 1 mL of acetylation mixture and gentle agitation for 1 h. The resin was filtered and washed with 1 mL of DMF (3 x), DCM (3 x), DMF (3 x), MeOH (3 x) and diethyl ether (3 x) and dried over $P_2O_5$ for at least 3 h. Peptides were cleaved from the resin by addition of 2 mL of TFA deprotection mixture and gentle shaking for 2 h. The solution from the syringe was collected separately in a 15 mL Falcon tube and the resin was washed once with 1 mL of TFA deprotection mixture. To the 3 mL of deprotection mixture was added 12 mL of cold diethyl ether. After precipitation (1 h, - 20 °C) the solid was centrifuged (5 min, 3000 rpm), washed with cold diethyl ether (4 mL), centrifuged again and left to dry at RT for 3 h. The solid was dissolved in DMSO:$CH_3CN$:$H_2O$ (1:4.5:4.5) and purified by preparative LCMS. Pure fractions were combined and lyophilized to afford ACC substrates. Substrate **S18**: LCMS – 18 min, 5 to 98% $CH_3CN$ in $H_2O$, Rt 1.14 min; HRMS (ESI, positive mode) found 764.3867 ($[C_{36}H_{49}N_{11}O_8+H]^+$ requires 764.3844). Substrate **S19**: LCMS – 18 min, 5 to 98% $CH_3CN$ in $H_2O$, Rt 7.62 min; HRMS (ESI, positive mode) found 963.4859 ($[C_{49}H_{62}N_{12}O_9+H]^+$ requires 963.4841).

1. Bock, N. *et al.* Engineering osteoblastic metastases to delineate the adaptive response of androgen-deprived prostate cancer in the bone metastatic microenvironment. *Bone Res.* **7**, 13 (2019).
2. Shokoohmand, A. *et al.* Microenvironment engineering of osteoblastic bone metastases reveals osteomimicry of patient-derived prostate cancer xenografts. *Biomaterials* **220**, 119402 (2019).
3. Chemical Computing Group ULC. Molecular Operating Environment (MOE). 1010 Sherbooke St. West, Suite #910, Montreal, QC, (2020).
4. Skala, W. *et al.* Structure-function analyses of human kallikrein-related peptidase 2

establish the 99-loop as master regulator of activity. *J. Biol. Chem.* **289**, 34267–34283 (2014).

5. Ménez, R. *et al.* Crystal Structure of a Ternary Complex between Human Prostate-specific Antigen, Its Substrate Acyl Intermediate and an Activating Antibody. *J. Mol. Biol.* **376**, 1021–1033 (2008).

### 5 Supplementary Figures

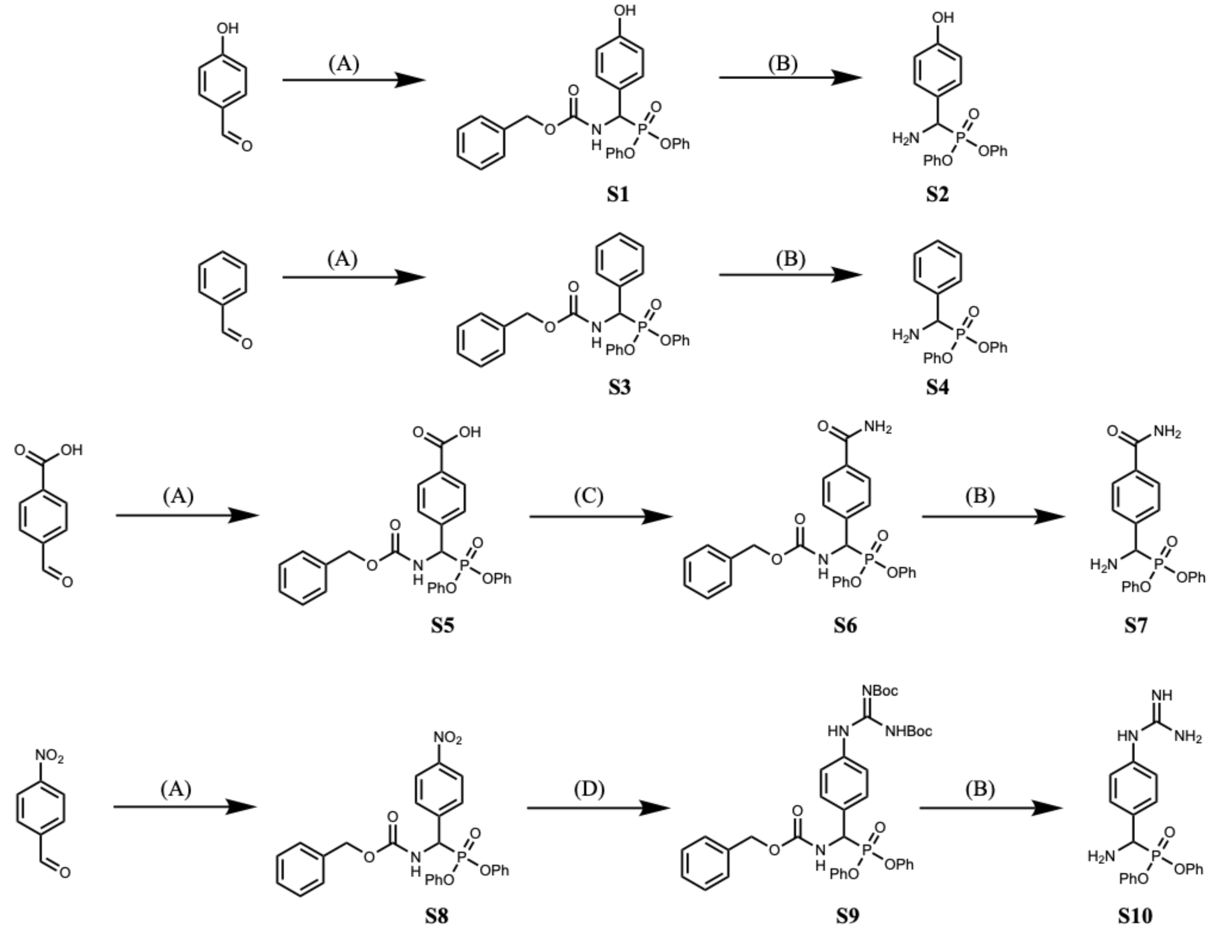

**Figure S1**: Synthesis of amino-diphenyl phosphonate warheads. (**A**) Benzyl carbamate, $P(OPh)_3$, AcOH, 90 °C, 2 h (**B**) 33 % HBr in AcOH, RT, 2 h (**C**) i. Boc anhydride, pyridine, THF, 50 °C, 30 min; ii. $(NH_4)_2CO_3$, 6 h, 50 °C (**D**) i. Fe Powder, AcOH, 70 °C, 2 h; ii. *N*,*N'*-Di-Boc-1H-pyrazole-1-carboxamidine, $NEt_3$, DCM, RT, O/N.

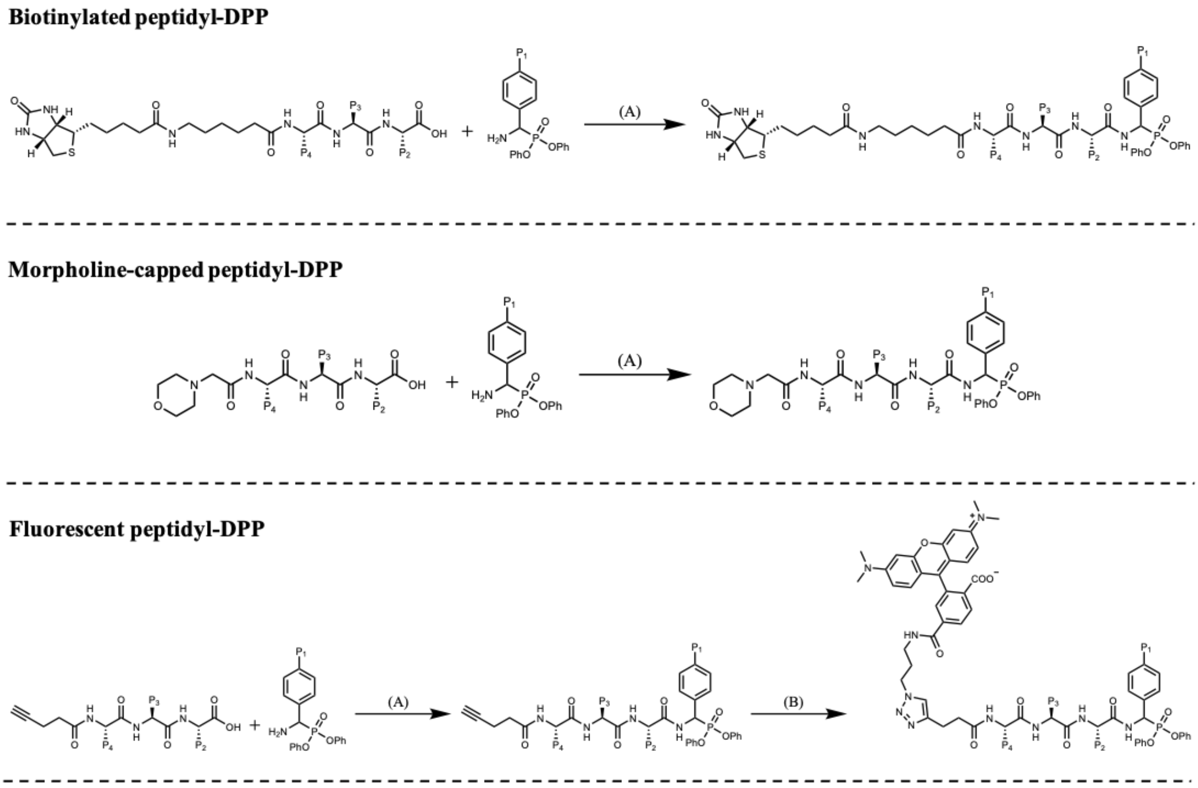

**Figure S2**: Synthesis of morpholine-capped, biotinylated and fluorescent peptidyl-diphenyl phosphonate analogues. Peptides were synthesized by standard solid-phase peptide synthesis using 2-chlorotrityl resin (**A**) HATU, 2,4,6-trimethylpyridine, DMF, RT, 2h (**B**) Cy5-/Fluorescein-/TAMRA-azide, $CuSO_4$, sodium ascorbate, DMF:$H_2O$ (4:1), RT, O/N.

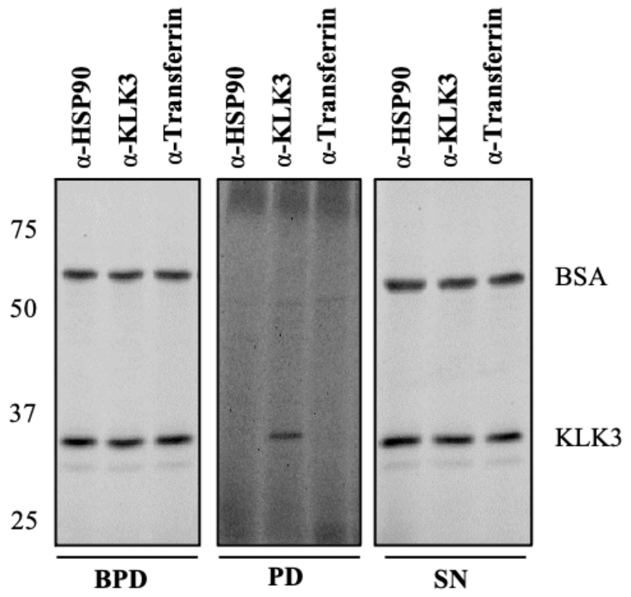

**Figure S3**: Immuno-enrichment of active KLK3 with a KLK3 antibody after labelling with 20 µM ABP **3** for 1 h. ABP **3** sticks to residual bovine serum albumin (BSA), which can be prevented by precipitating protein samples after probe treatment. BPD = Before pull-down, PD = Pull-down and SN = Supernatant.

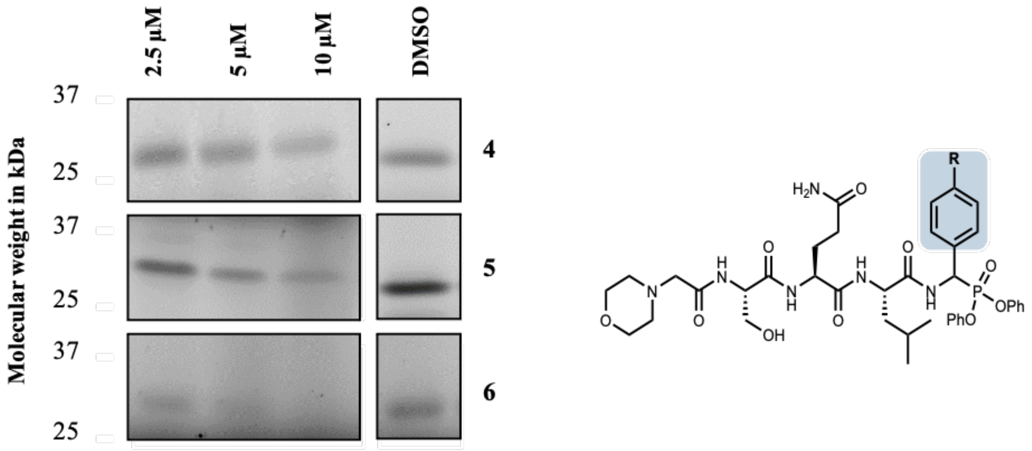

**P1 Optimization**

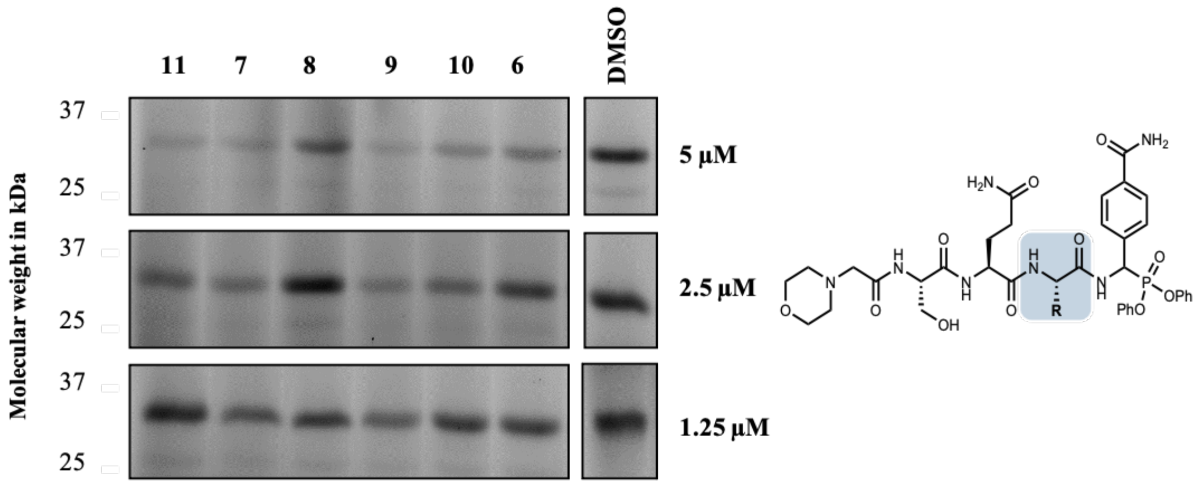

**P2 Optimization**

**Figure S4**: Typical gel obtained from assessment of the potency of P1 (top) and P2 (bottom) morpholine-DPP analogues against KLK3 using competitive ABPP. LNCaP CM was incubated with different concentrations of each analogue for 1 h prior to treatment with 20 µM **3** for 1 h. Residual KLK3 activity was calculated using densitometry (ImageQuant) by comparing to a DMSO treated sample.

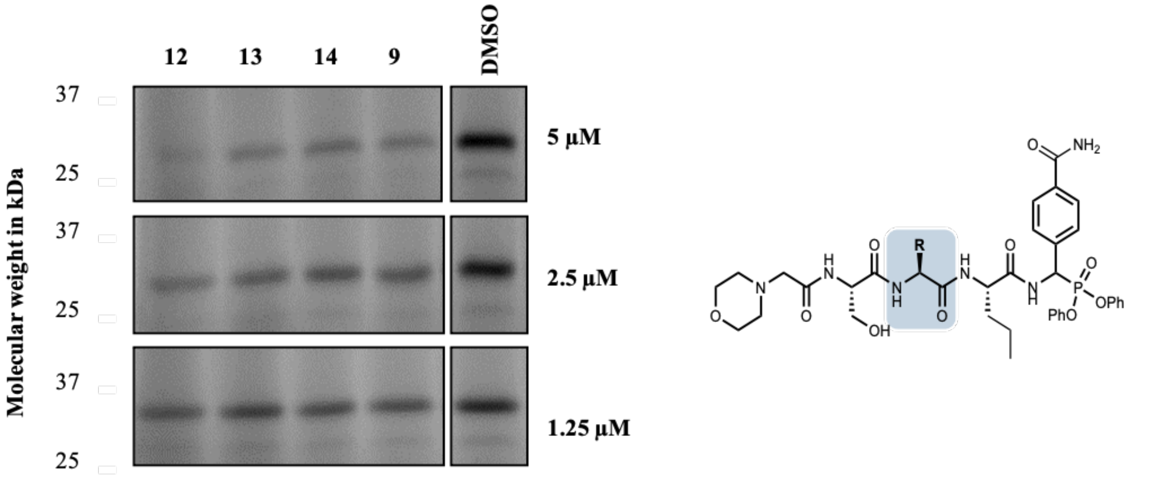

P3 Optimization

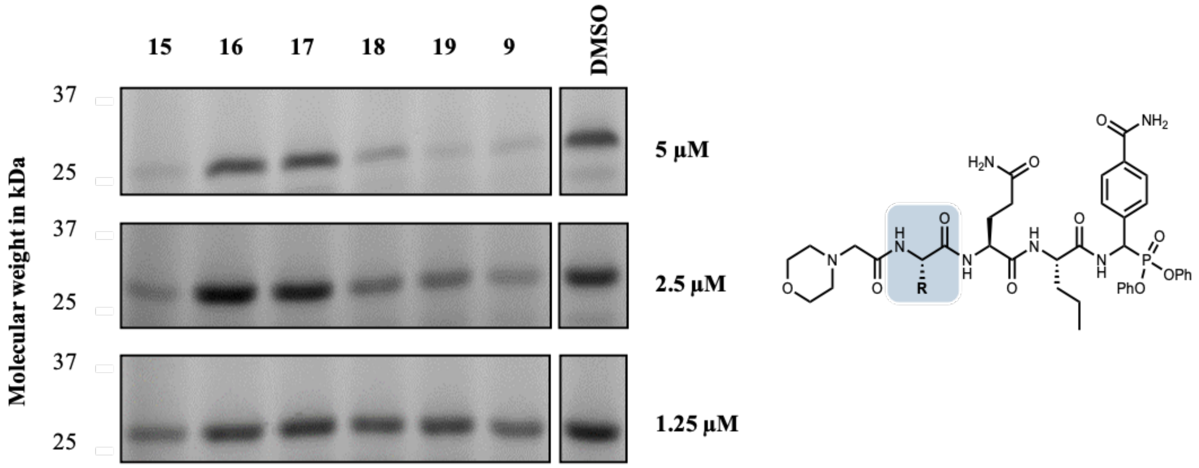

P4 Optimization

**Figure S5**: Typical gels obtained from assessment of the potency of P3 (top) and P4 (bottom) morpholine-DPP analogues against KLK3 using competitive ABPP. LNCaP CM was incubated with different concentrations of each analogue for 1 h prior to treatment with 20 µM **3** for 1 h. Residual KLK3 activity was calculated using densitometry (ImageQuant) by comparing to a DMSO treated sample.

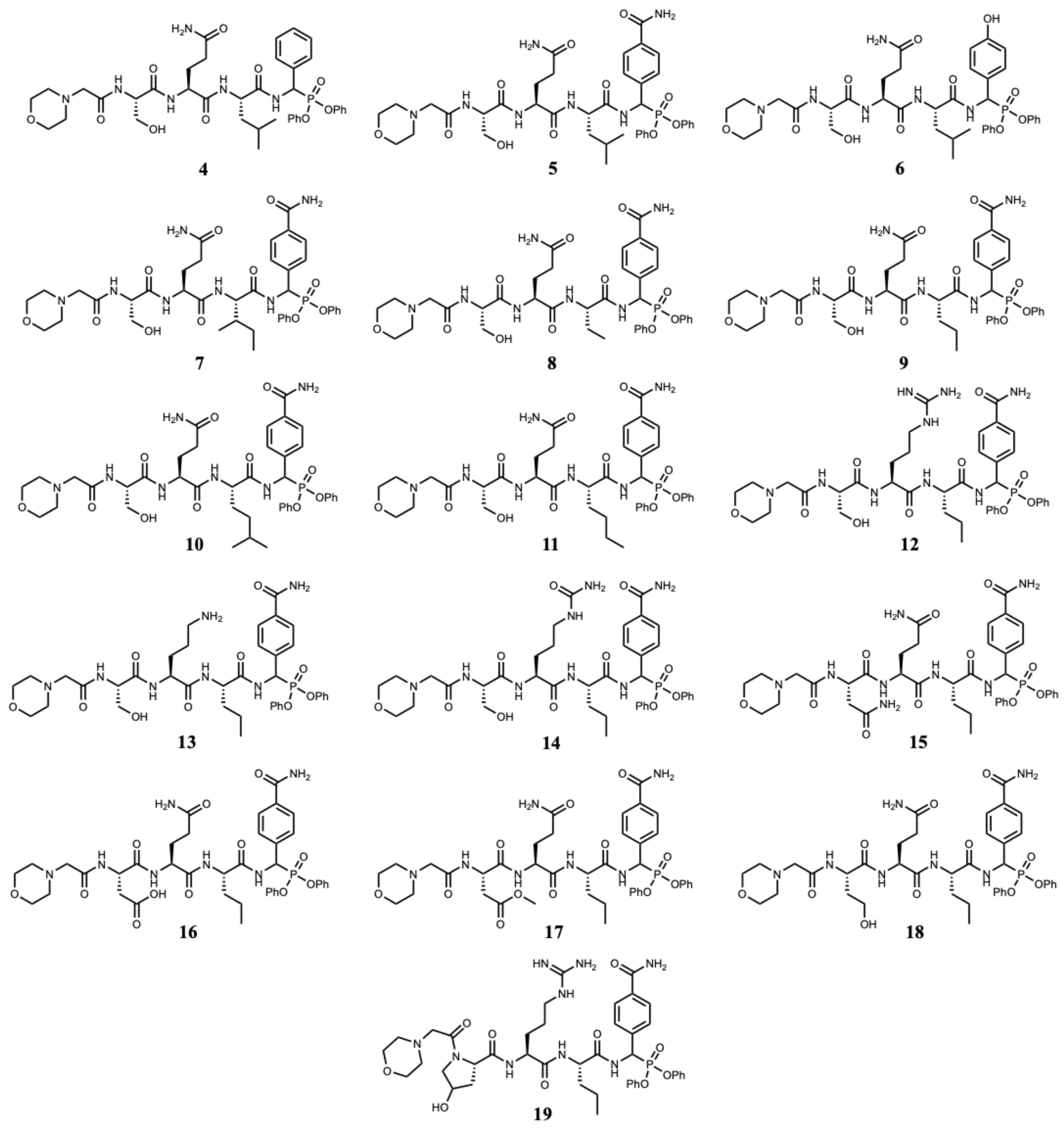

**Figure S6**: Structures of the KLK3 morpholine-DPP analogues **4-19**.

| Lane | 1 | 2 | 3 | 4 | 5 | 6 | 7 |
|---|---|---|---|---|---|---|---|
| PMSF (1mM) | + | - | - | - | - | - | - |
| 3 (μM) | 1 | .125 | .25 | .5 | 1 | 5 | 10 |

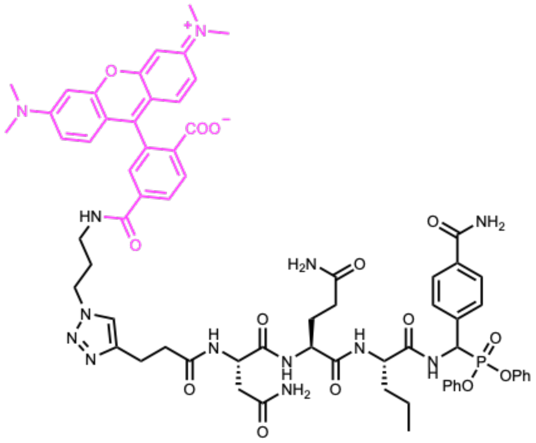

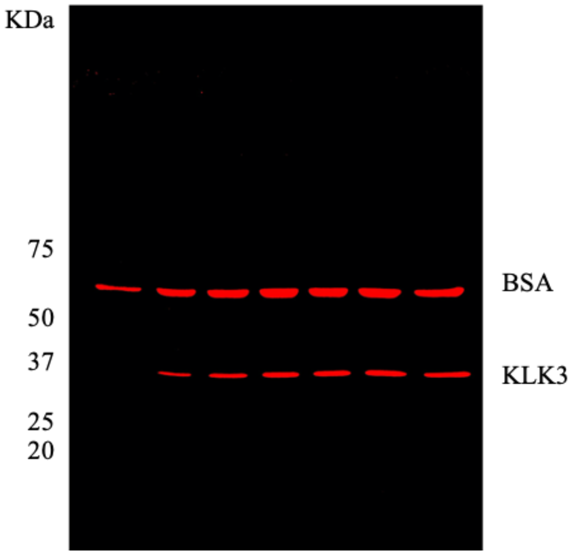

**Figure S7**: Structure (left) and labeling profile (right) in LNCaP CM of **21**. LNCaP CM was labeled with different concentrations of **21** for 1 h. Pre-treatment of CM with PMSF reduces KLK3 labelling but not BSA labelling. Non-specific BSA labelling can be prevented by precipitating protein samples after probe treatment as shown in **Fig. 2B** and **Fig. 6B**.

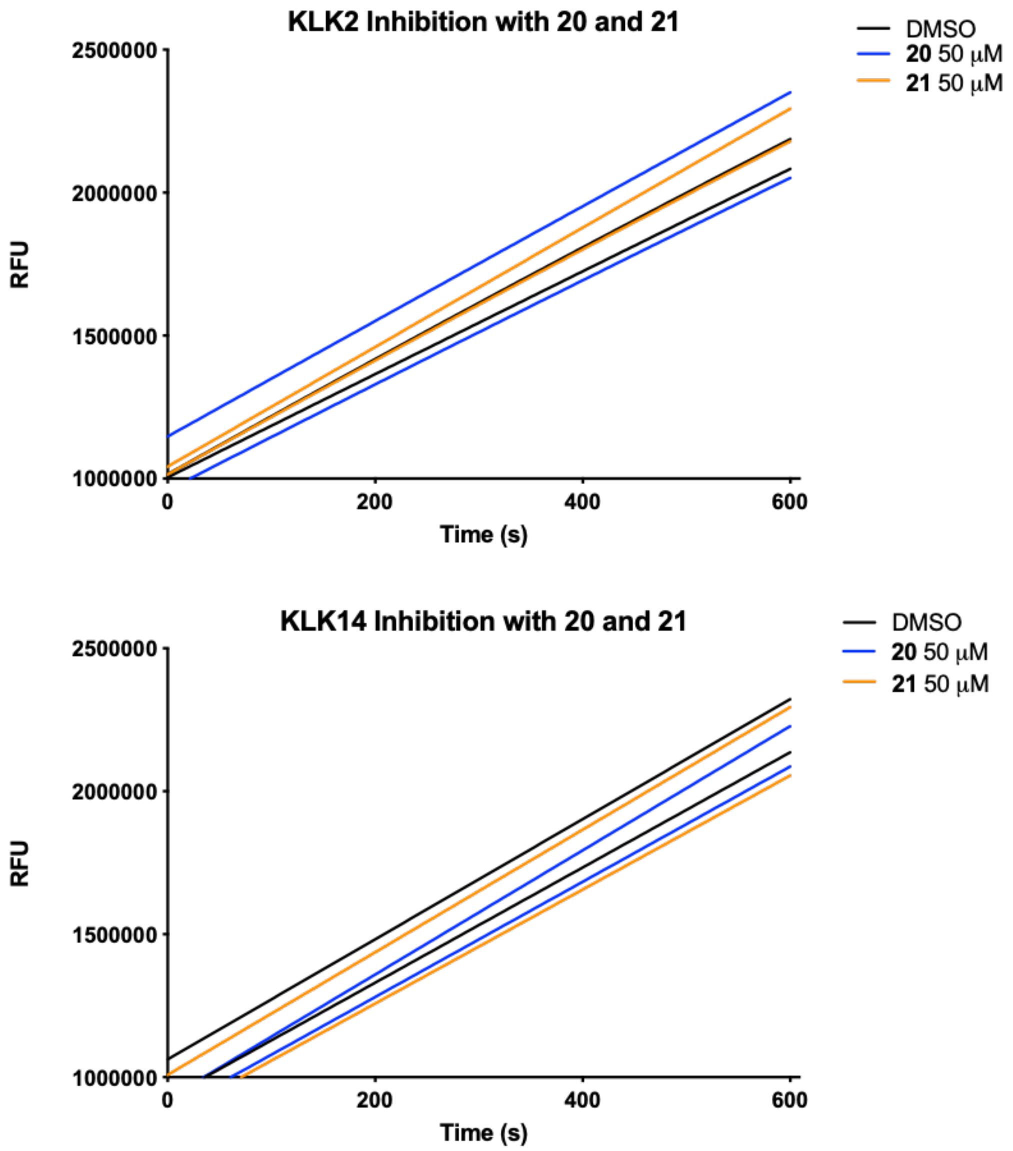

**Figure S8**: Inhibition of KLK2 (top) or KLK14 (bottom) after incubation with 50 μM of **20** or **21** for 30 minutes

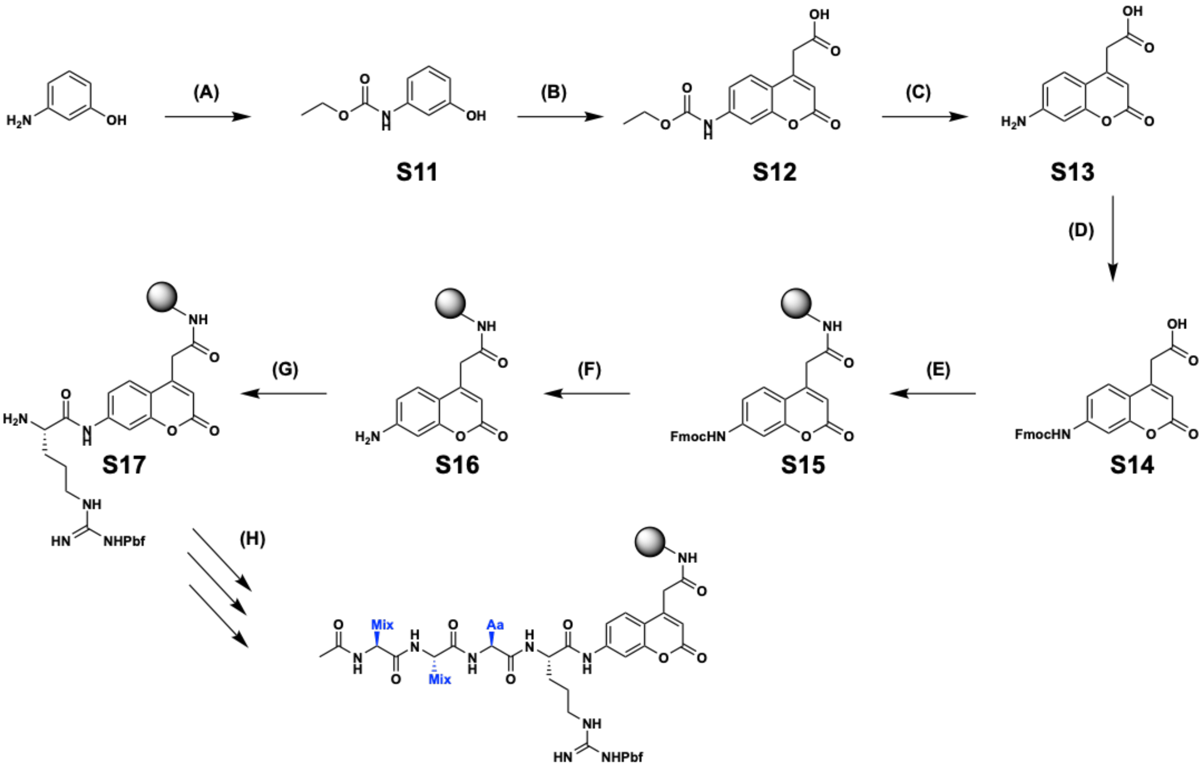

**Figure S9**: Synthesis of Positional Scanning Libraries. (**A**) Ethyl chloroformate, EtOAc, reflux; 99 % (**B**) 1,3-acetonedicarboxylicacid, 70% $H_2SO_4$, RT; 56 % (**C**) 10 M NaOH, reflux; 66 % (**D**) (i) TMSCl, DIPEA, DCM, reflux (ii) Fmoc-Cl, RT; 82 % (**E**) Rink amide AM resin, HOBt, DICI, DMF, RT; quantitative (**F**) 20 % PIP/DMF, RT; quantitative (**G**) (i) Fmoc-Arg(Pbf)-OH, HATU, 2,4,6-trimethylpyridine, DMF, RT (ii) AcOH, 3-nitro-1,2,4-triazole, DICI, DMF, RT; 48 % (**H**) P2 amino acid coupling and isokinetic mixture coupling: DICI, HOBt, RT; Acetylation: Acetic acid, HBTU, DIPEA, RT

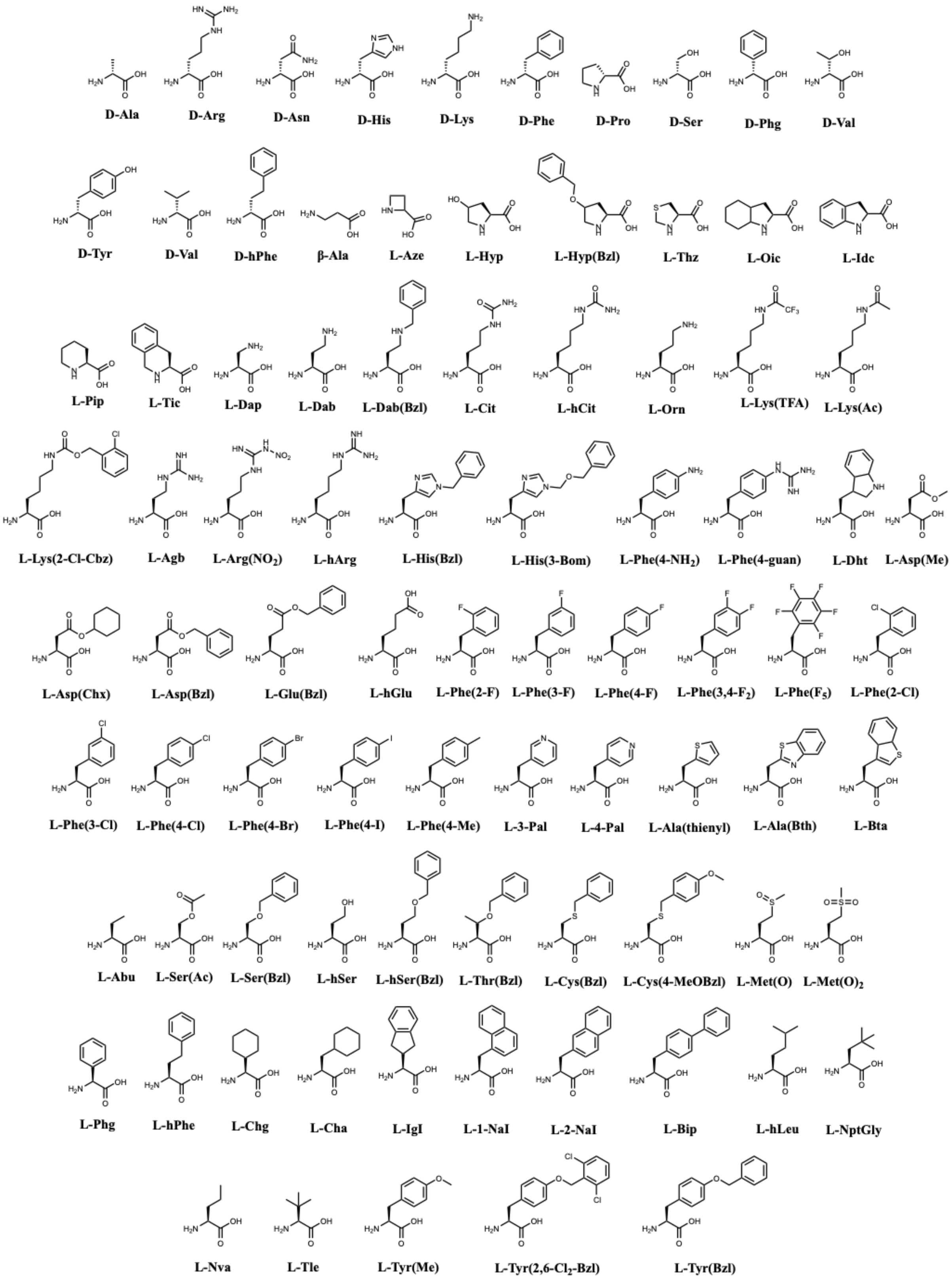

**Figure S10**: The amino acids used to generate the Positional Scanning Library.

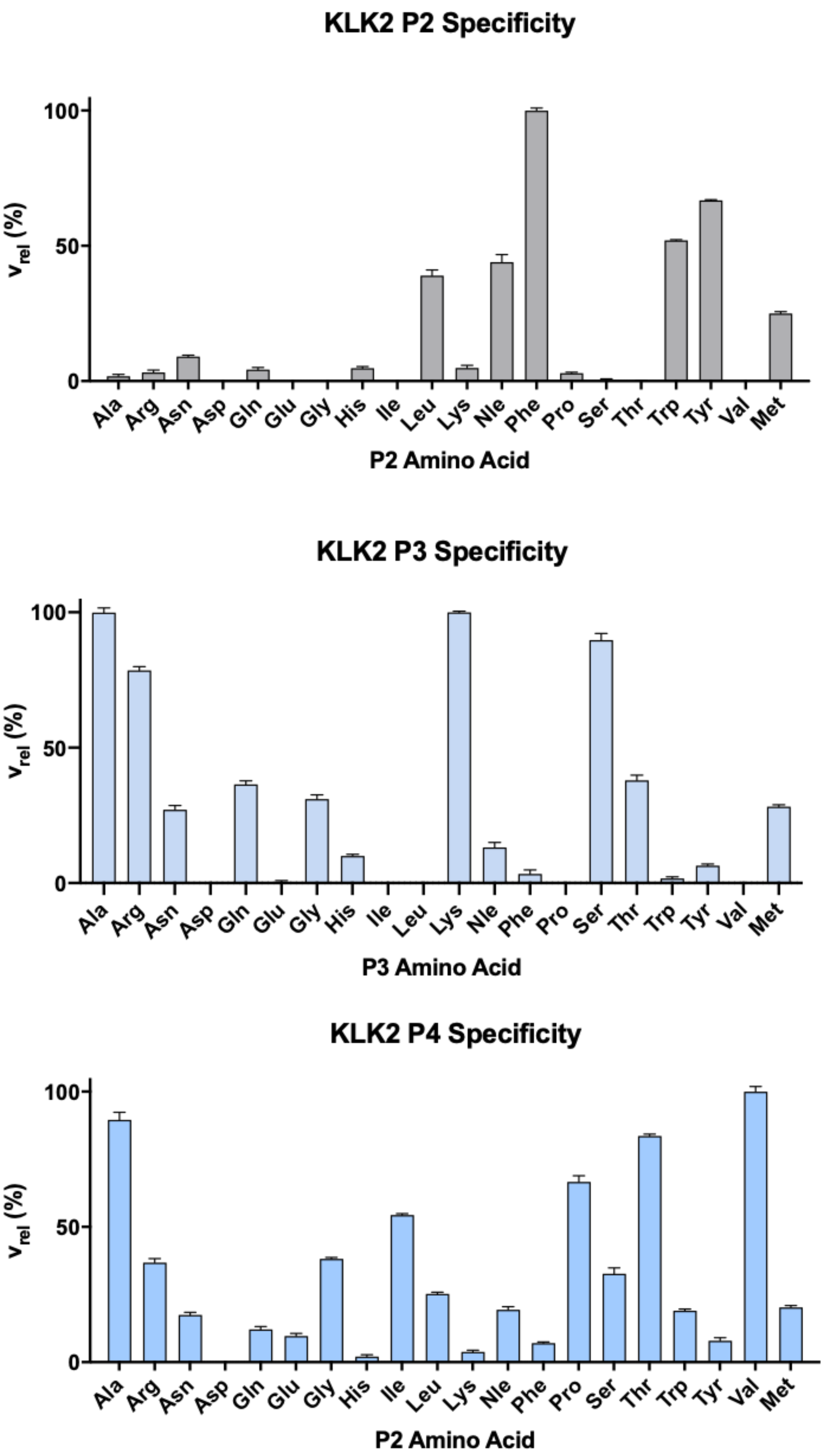

**Figure S11**: S2-S4 specificity profiles of KLK2 using natural amino acids.

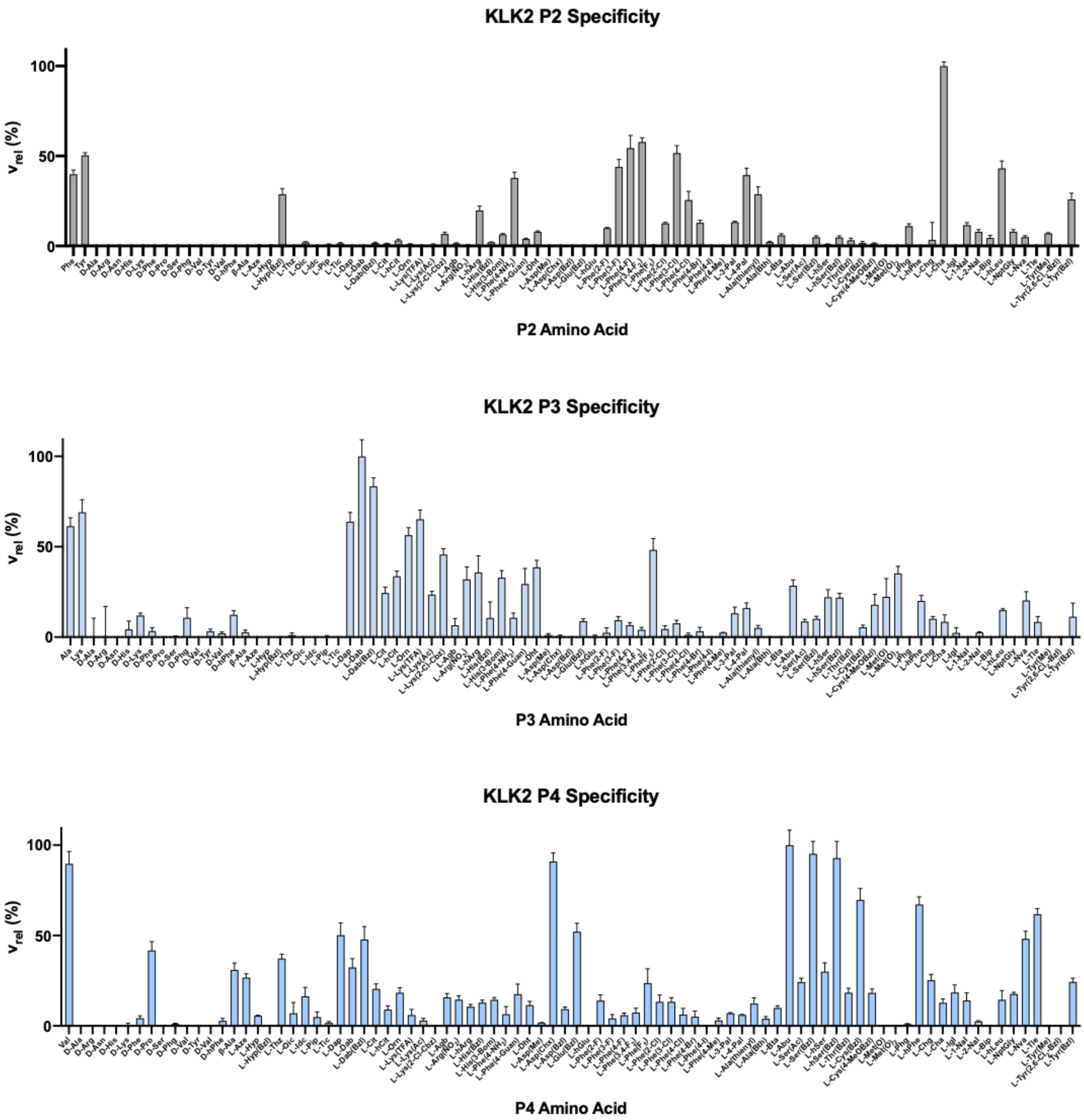

**Figure S12**: S2-S4 specificity profiles of KLK2 using non-natural amino acids.

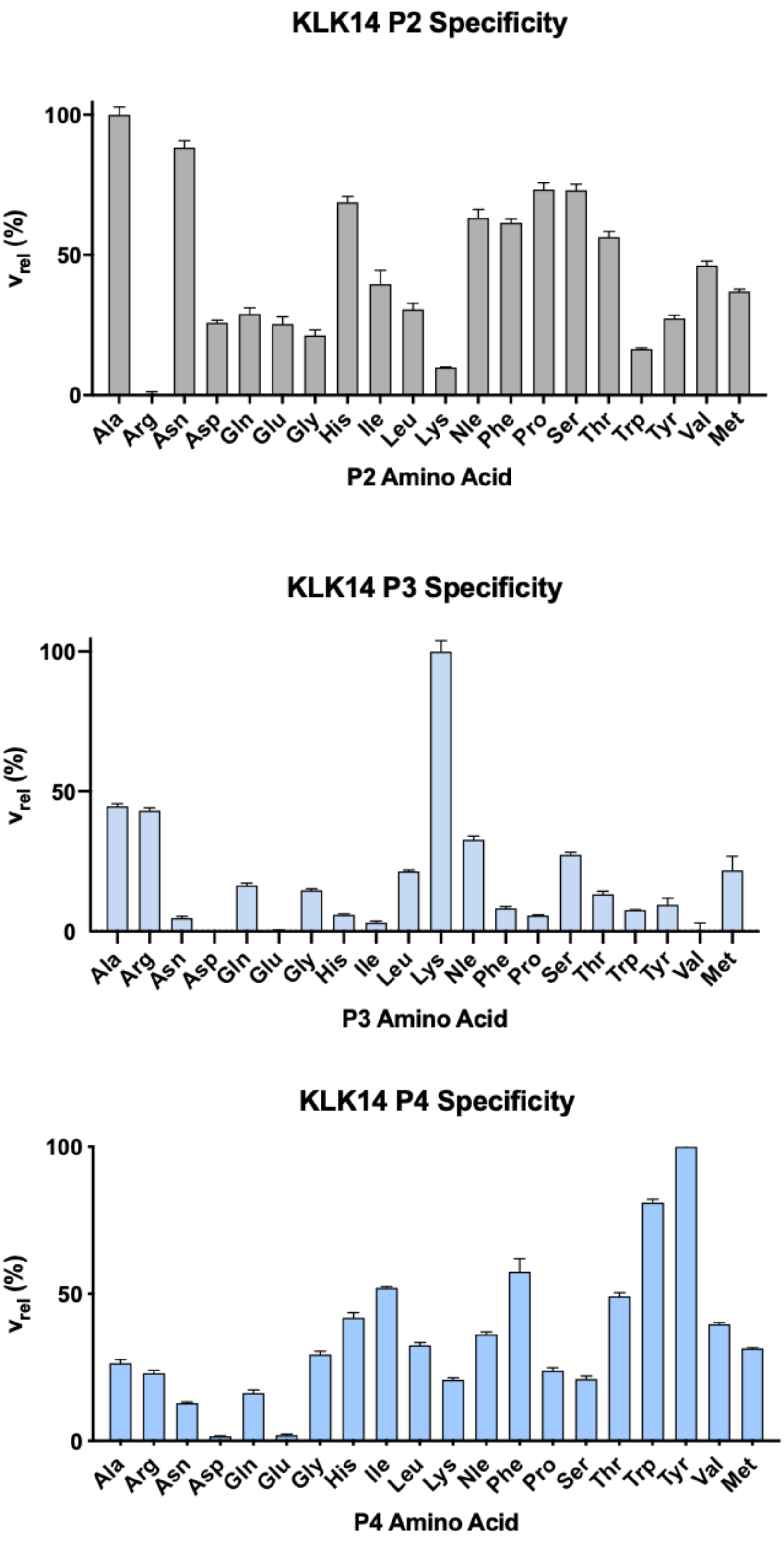

**Figure S13**: S2-S4 specificity profiles of KLK14 using natural amino acids.

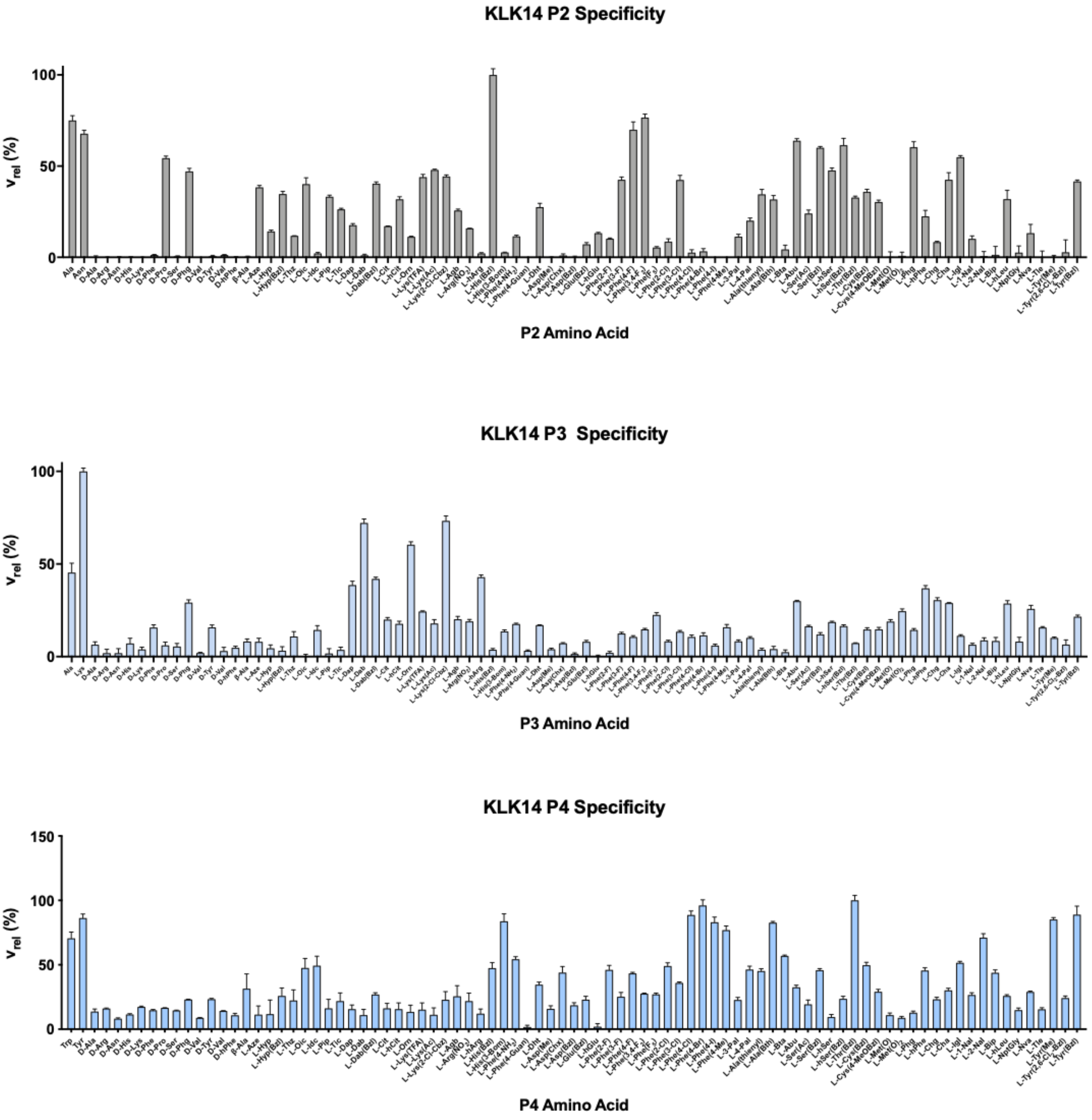

**Figure S14**: S2-S4 specificity profiles of KLK14 using non-natural amino acids.

**A**

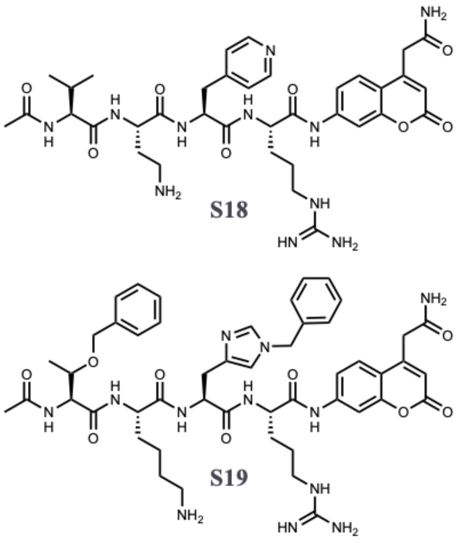

**B**

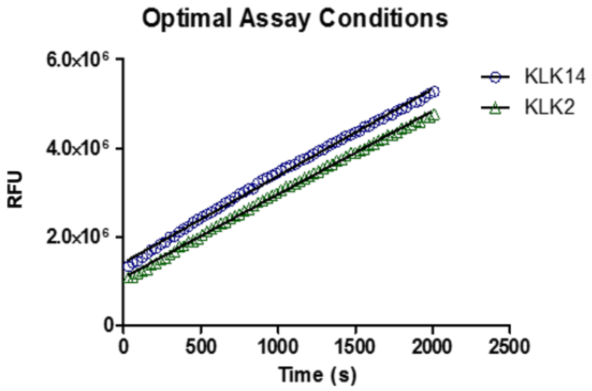

**C**

| KLK | Compound | $K_M$ (µM) | Kcat ($s^{-1}$) | Kcat/Km ( x$10^6$ $M^{-1}s^{-1}$) |
|---|---|---|---|---|
| **2** | Ac-Val-Dab-4Pal-Arg-ACC (S18) | 42 ± 2.5 | 870 ± 11.3 | 162.7 |
| **14** | Ac-Thr(Bz)-Lys-His(Bz)-Arg-ACC (S19) | 65 ± 4.2 | 10667 ± 11.3 | 20.6 |

**Figure S15**: KLK2 and KLK14 substrates. (**A**) Structures (**B**) Optimal assay conditions for kinetic analyses of KLK2 (0.625 nM KLK2, 5 µM S18) and KLK14 (37.5 pM KLK14, 5 µM S19) inhibitors (**C**) Michaelis-Menten ($K_M$) and rate ($k_{cat}$) constants for KLK substrates

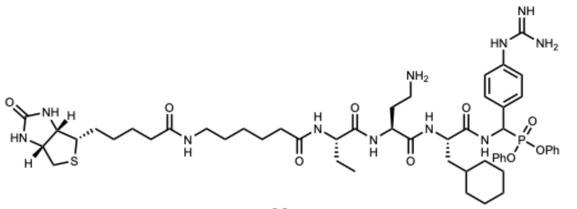

**22**

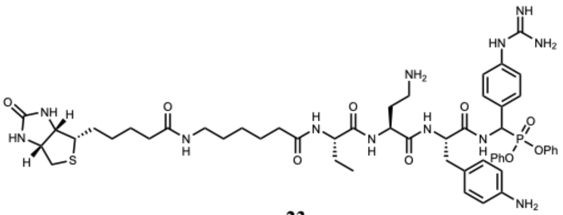

**23**

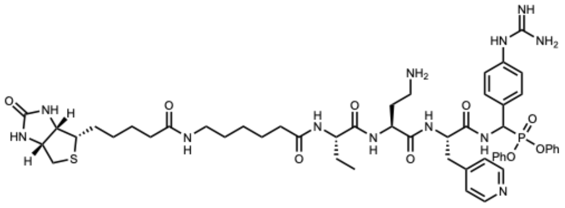

**24**

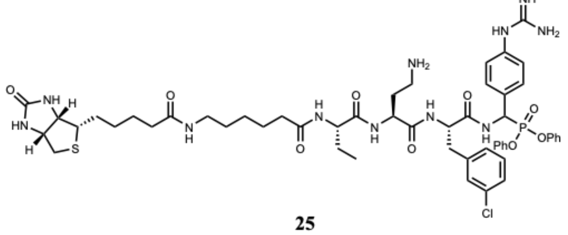

**25**

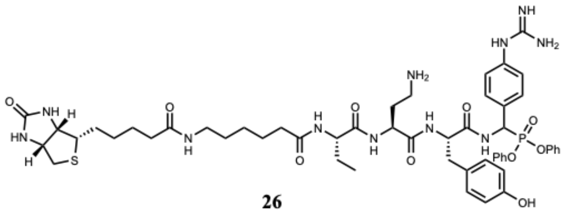

**26**

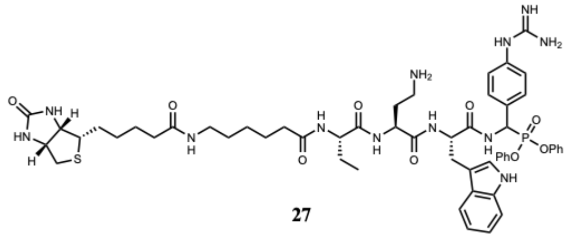

**27**

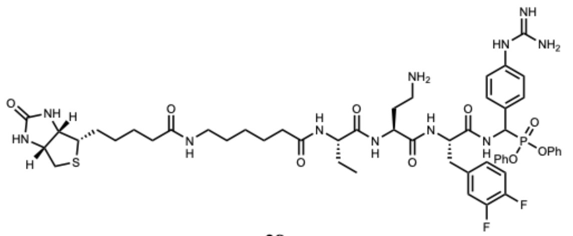

**28**

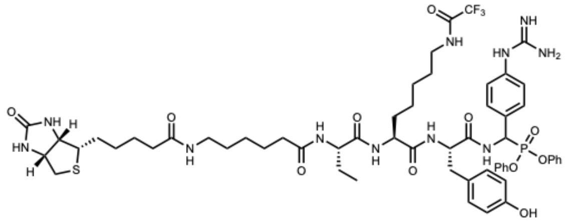

**29**

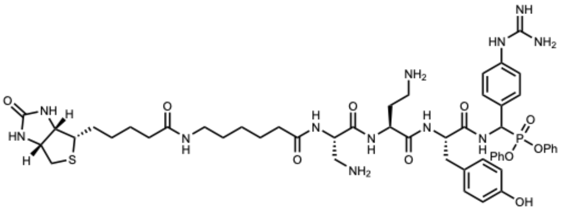

**30**

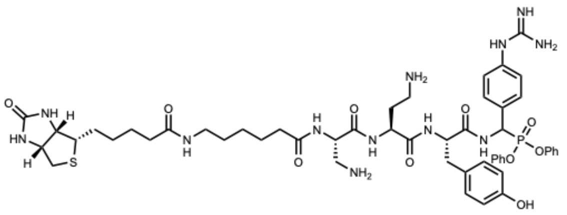

**31**

**Figure S16**: Structures of the KLK2 biotinylated peptidyl-DPP analogues **22-31**.

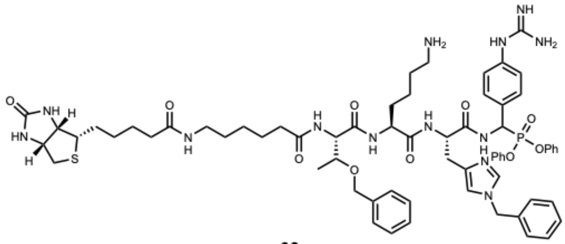

**32**

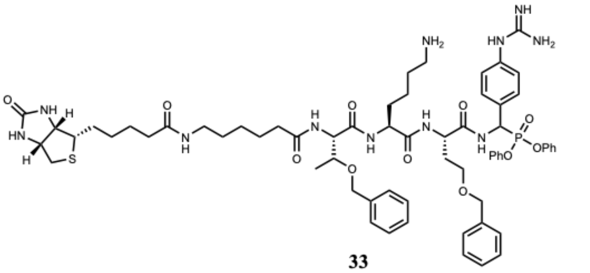

**33**

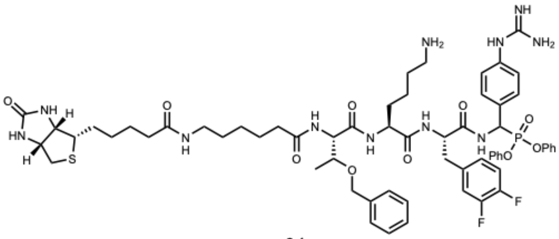

**34**

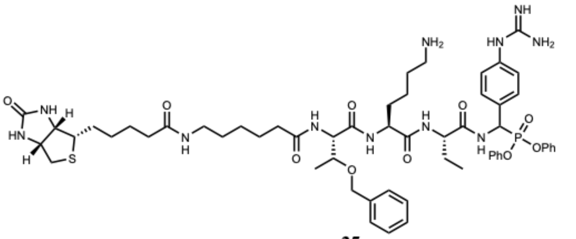

**35**

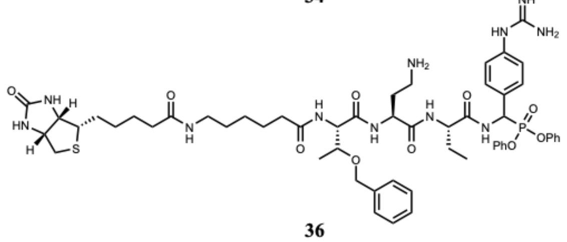

**36**

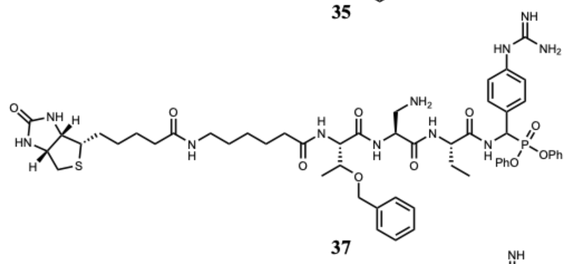

**37**

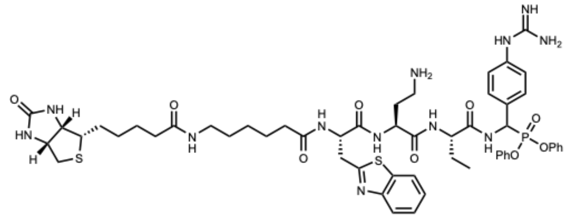

**38**

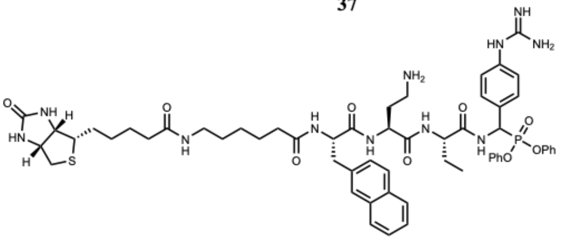

**39**

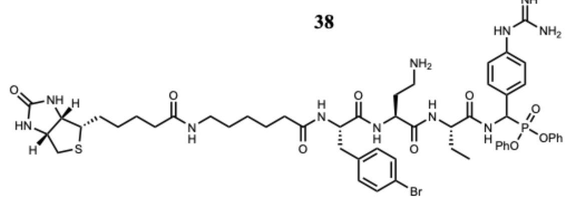

**40**

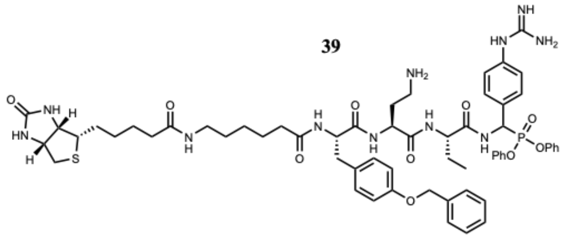

**41**

**Figure S17**: Structures of the KLK14 biotinylated peptidyl-DPP analogues **32-41**.

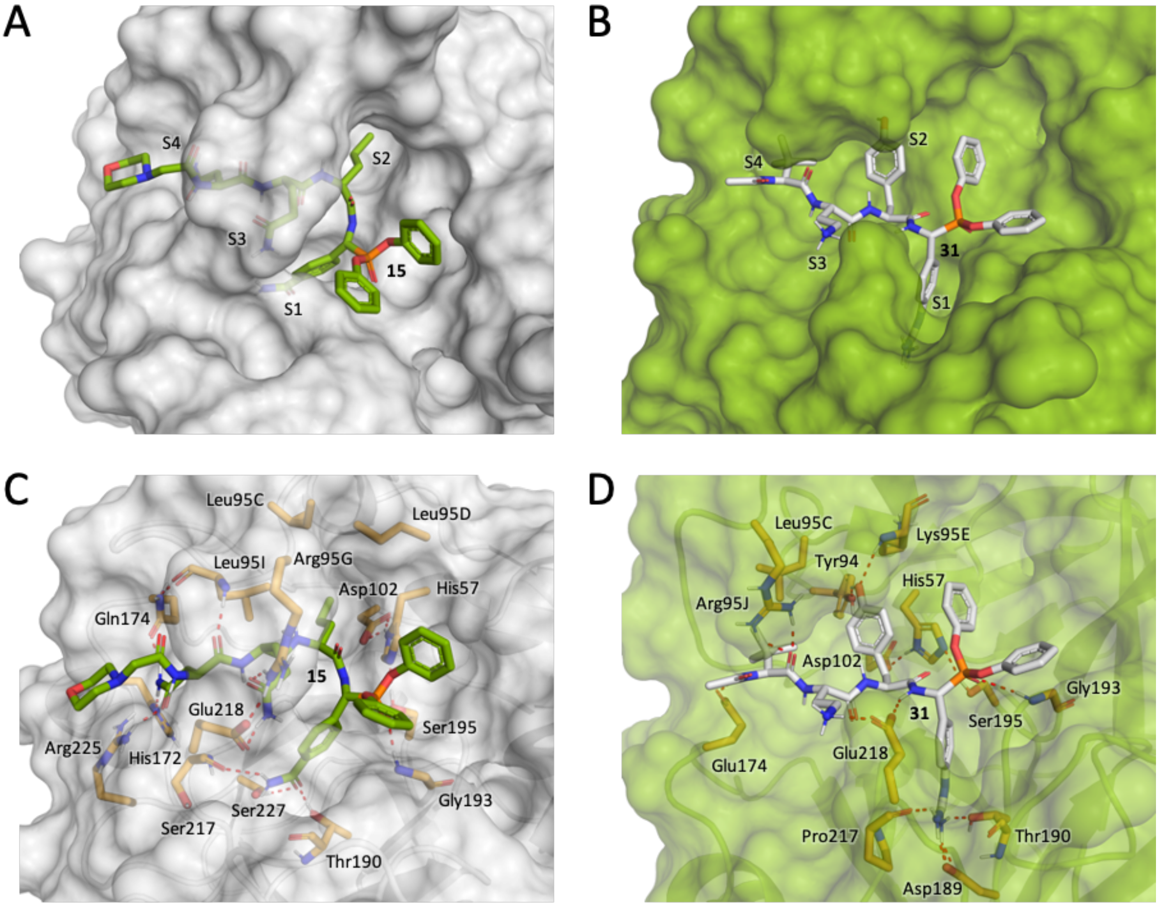

**Figure S18**: Binding of acyl KLK2_bABP and morpholino KLK3_bABP to the active site of KLK2 and KLK3, respectively, as proposed by molecular modelling. (**A**) Surface of KLK3 active site with docked probe morpholino KLK3_bABP. (**B**) Surface of KLK2 active site with docked probe acyl KLK2_bABP. (**C**) H-bond interactions of morpholino KLK3_bABP with the KLK3 active site. (**D**) H-bond interactions of acyl KLK2_bABP with the KLK2 active site.

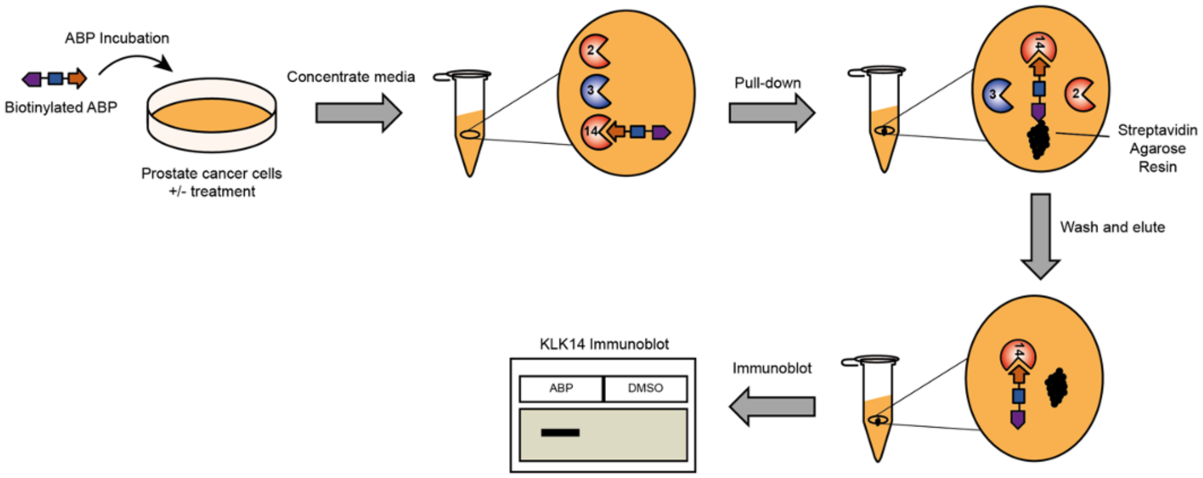

**Figure S19**: Activity-based protein profiling workflow using biotinylated ABPs and streptavidin enrichment. (**A**) A biotinylated ABP is incubated with a plate of prostate cancer cells previously subjected to a defined perturbation. Conditioned media is collected and concentrated using a molecular weight cut-off spin filter. (**B**) Probe-labeled proteins are immobilized on magnetic streptavidin beads. (**C**) Unlabeled proteins are discarded, and enriched proteins are eluted from beads by boiling in sample loading buffer (**D**) KLK enrichment is confirmed by immunoblotting with a KLK antibody.

| Lane | 1 | 2 | 3 |
|---|---|---|---|
| **22** (1 μM) | - | + | - |
| **31** (1 μM) | - | - | + |

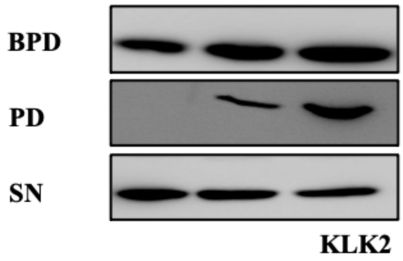

| Lane | 1 | 2 | 3 | 4 | 5 |
|---|---|---|---|---|---|
| **20** (μM) | 0 | .125 | .25 | .5 | 1 |

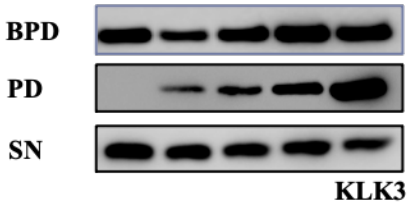

| Lane | 1 | 2 | 3 |
|---|---|---|---|
| **32** (1 μM) | - | + | - |
| **39** (1 μM) | - | - | + |

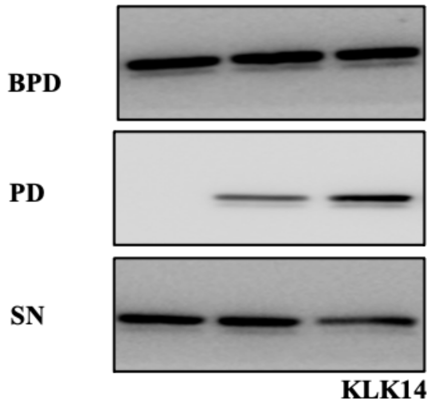

**Figure S20**: Enrichment from LNCaP-K14 CM of KLK2, KLK3 and KLK14 using the workflow outlined in Fig. S19 after 1 h treatment with **22**/**31**, **20** or **32**/**39**, respectively.

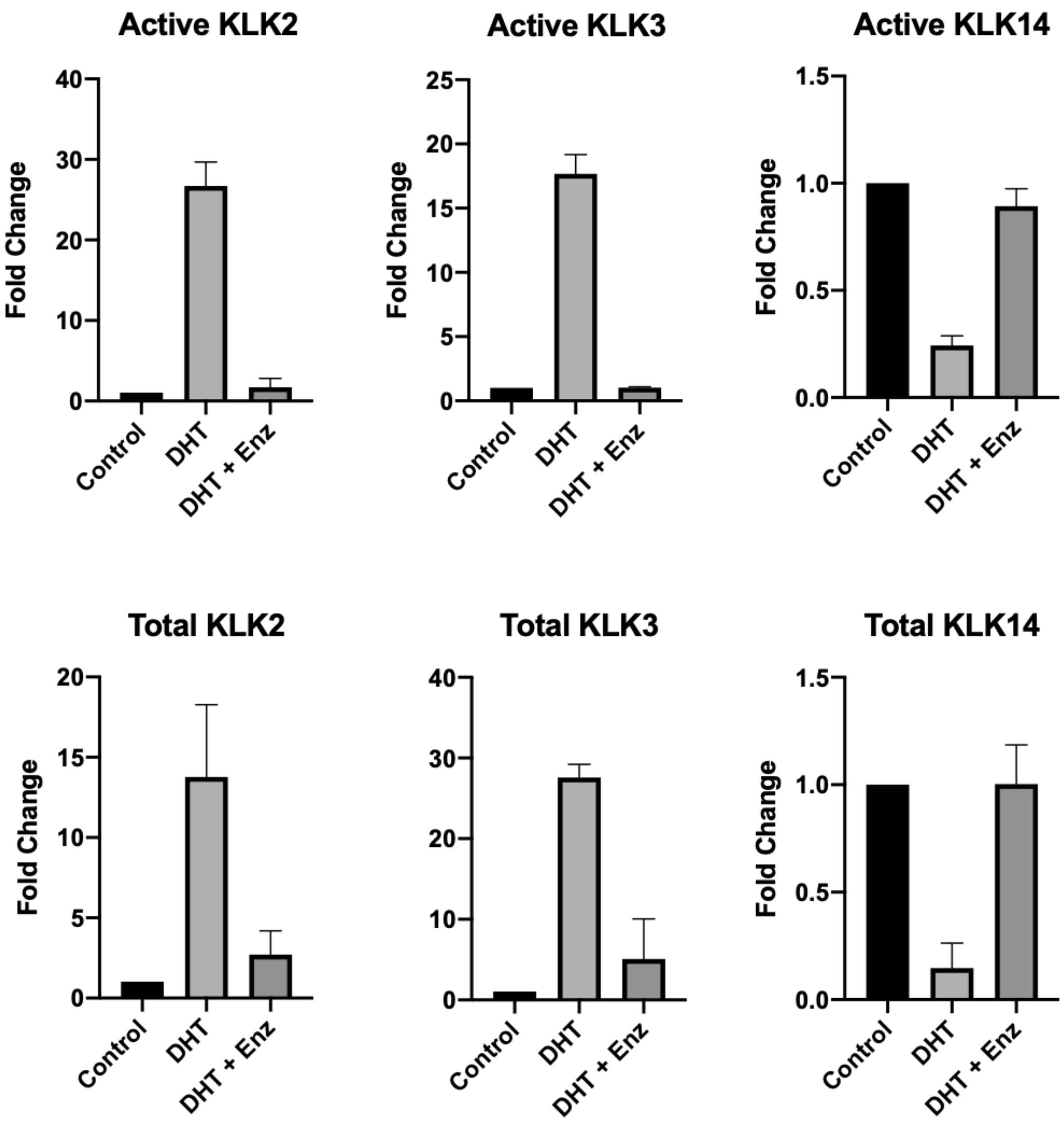

**Figure S21**: Quantification of KLK activity in figure 4B by densitometry (ImageQuant). Each data point is normalized to the control sample and is a mean value ± SEM (N=3).

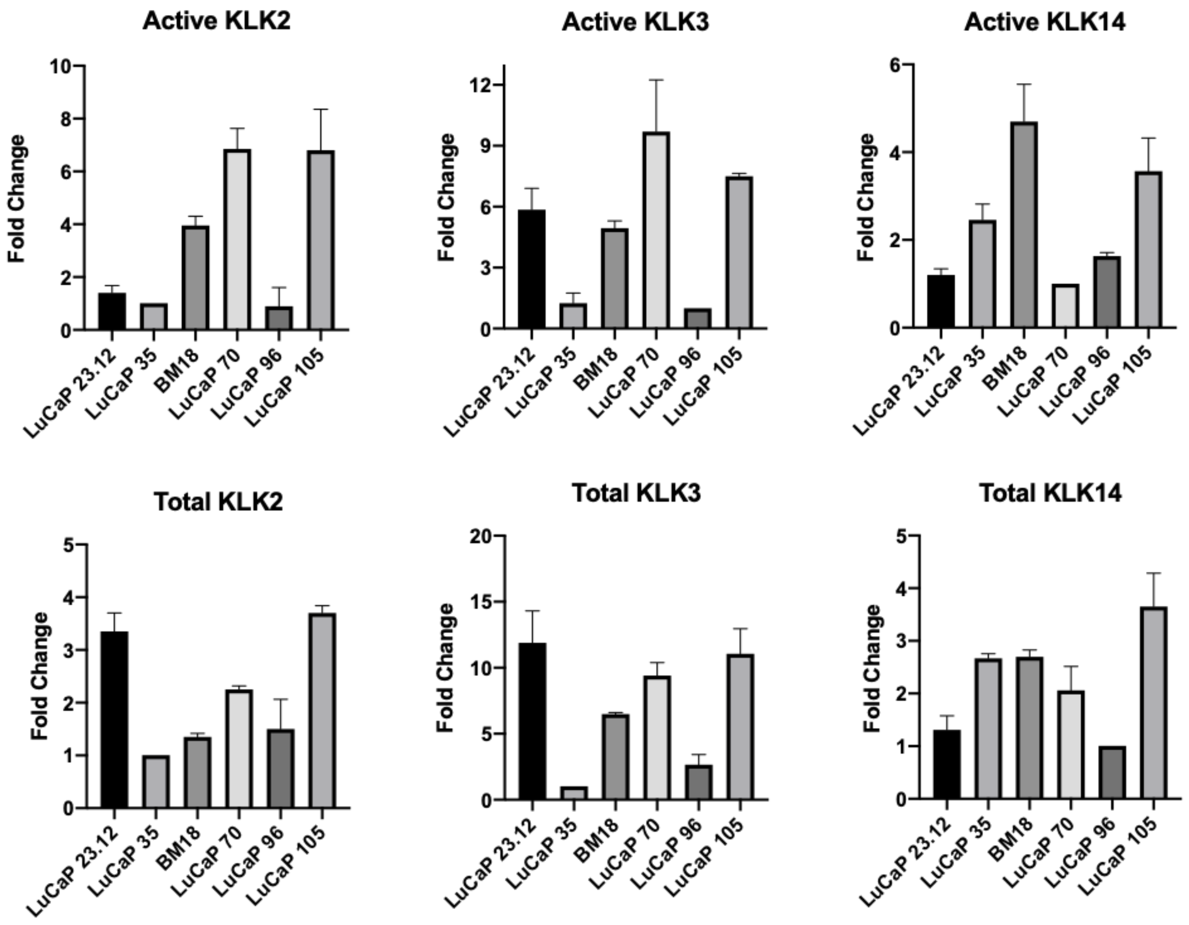

**Figure S22**: Quantification of KLK activity in figure 4C by densitometry (ImageQuant). Each data point is normalized to the PDX with the least intense band and is a mean value ± SEM (N=2).

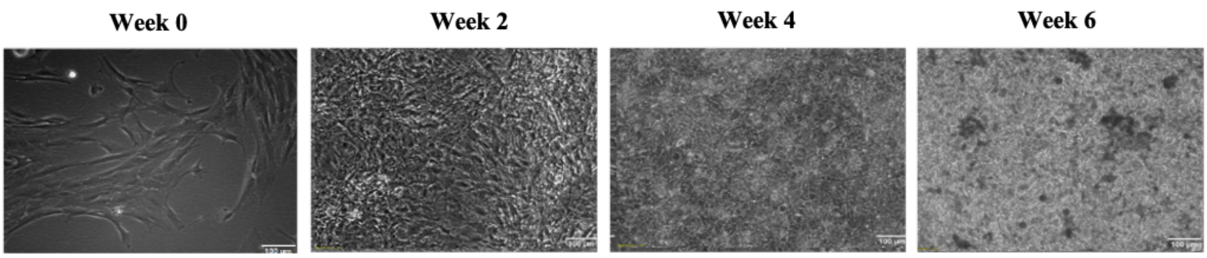

**Figure S23**: Osteoblast cell differentiation and mineralization by continued culturing in osteogenic media (OM). Visualized by phase contrast imaging.

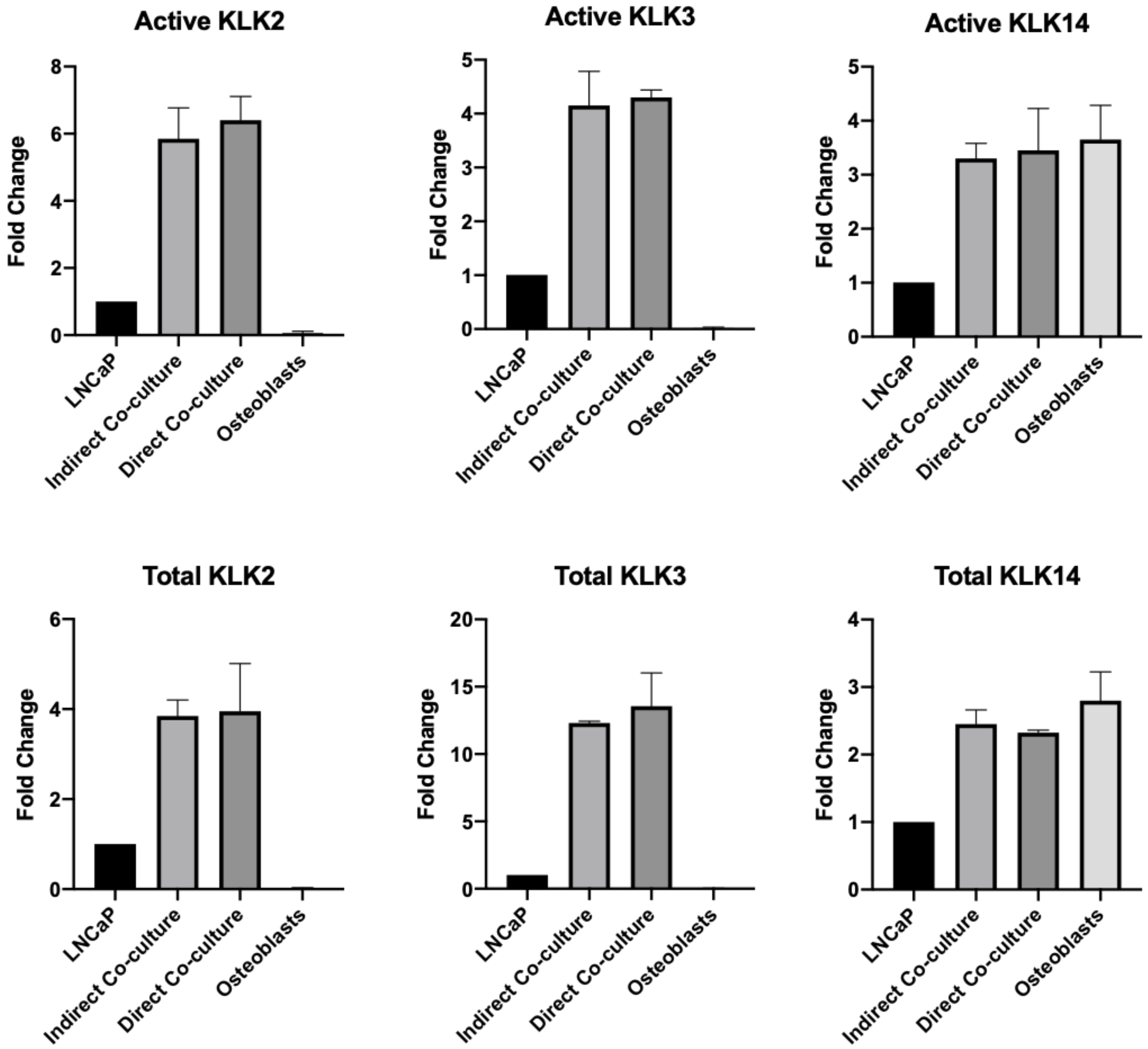

**Figure S24**: Quantification of KLK activity in figure 5B by densitometry (ImageQuant). Each data point is normalized to the LNCaP sample and is a mean value ± SEM (N=3).

| Fluorescent Probes | $K_{inact}/K_I$ ($M^{-1}s^{-1}$) | | |
|---|---|---|---|
| ABP Analogue | KLK14 | KLK2 | Selectivity (fold-change) |
| Cy5-Tle-Dab-Tyr-Arg-DPP (**42**) | 202 ± 9 | 4392 ± 356 | 21 |
| Fluorescein-NaI-Dab-Abu-Arg-DPP (**43**) | 47325 ± 2301 | 164 ± 13 | 288 |

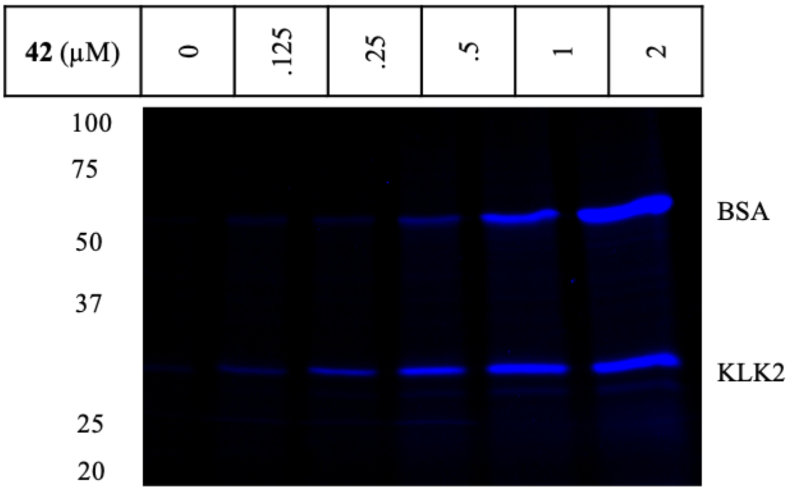

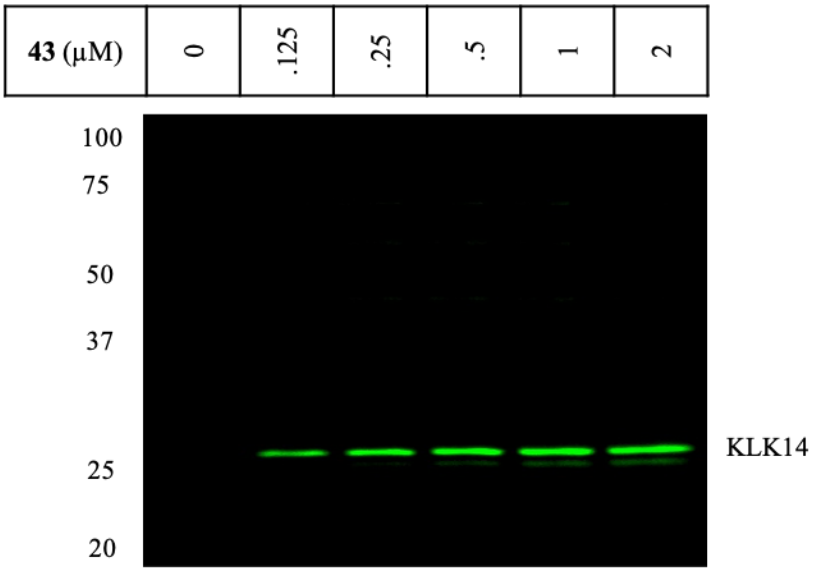

**Figure S25**: $K_{inact}/K_I$ values (top) and labeling profiles in LNCaP-K14 CM of **42** (middle) and **43** (bottom) after 1 h of treatment. Both probes show non-specific labelling to BSA, which can be prevented by precipitating proteins after treatment as seen in Fig. 6D.

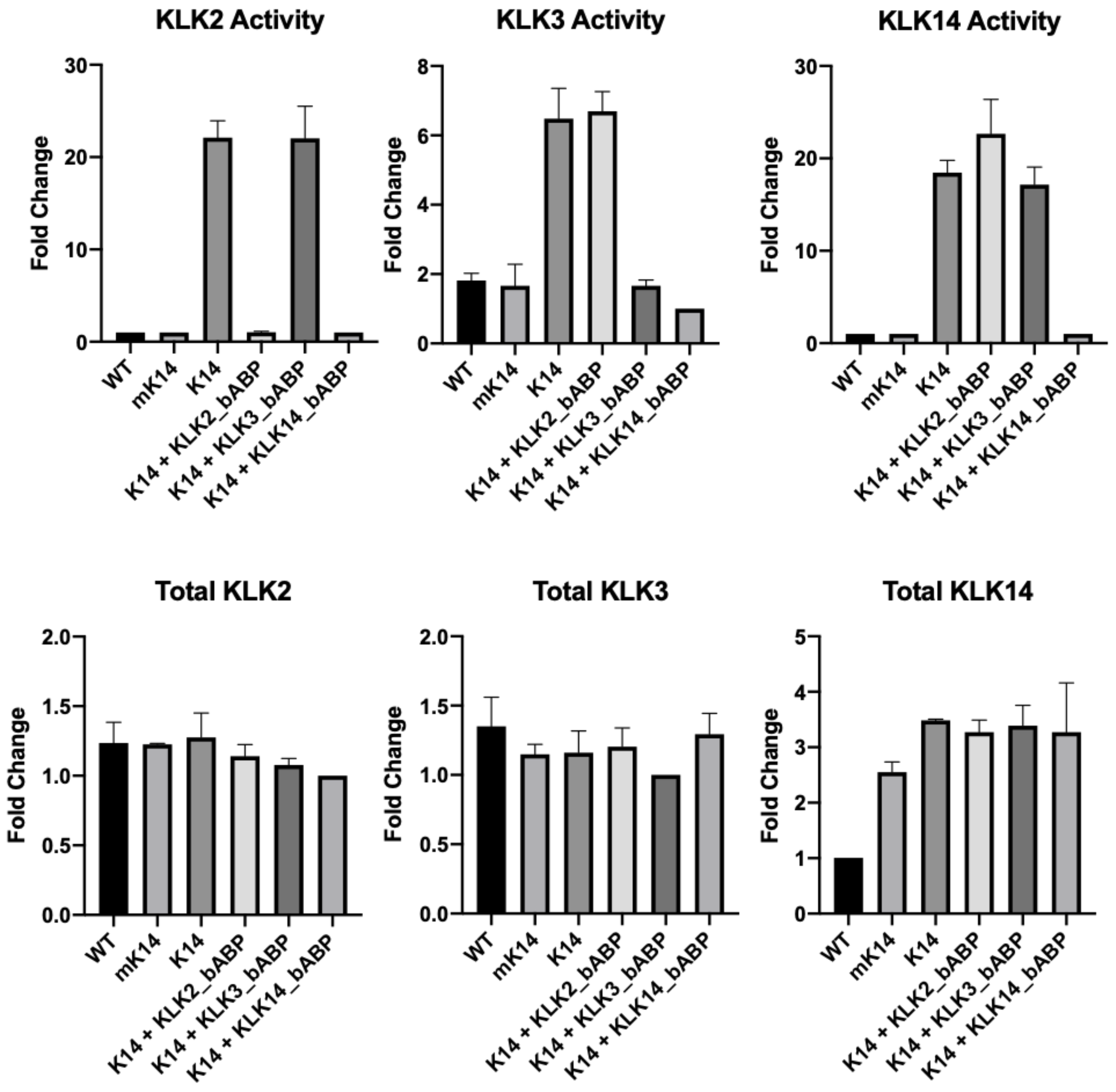

**Figure S26**: Quantification of KLK activity in figure 6B by densitometry (ImageQuant). Each data point is normalized to the least intense lane and is a mean value ± SEM (N=3)

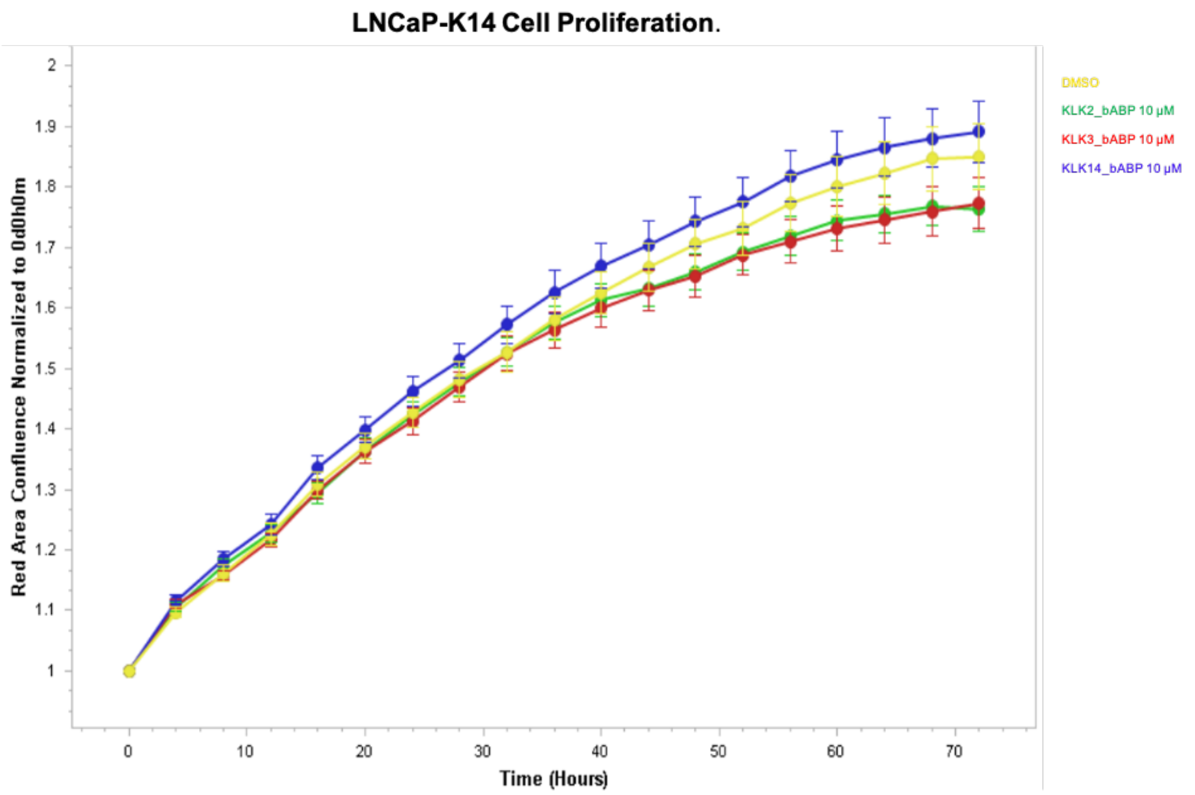

**Figure S27**: KLK2_bABP, KLK3_bABP and KLK14_bABP have no significant effect on the proliferation of LNCaP-K14 cells. Cell confluency was monitored by real-time imaging using an Incucyte S3 system (Essen Bioscience).

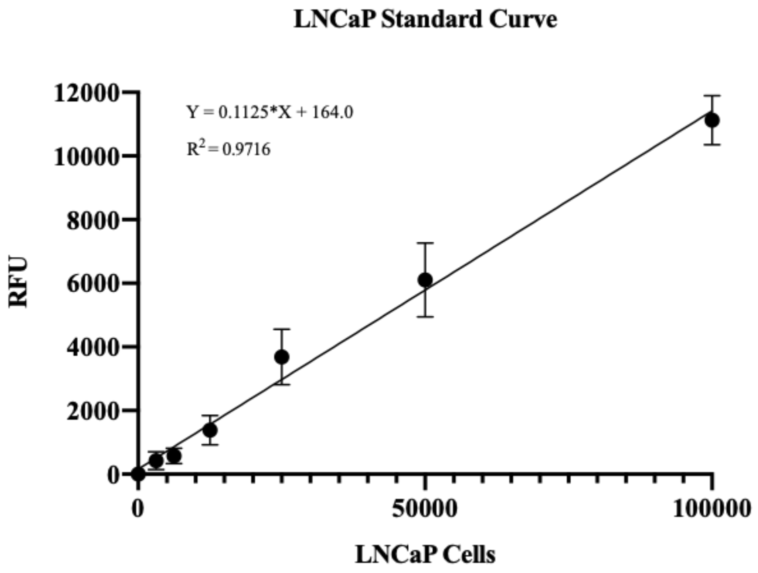

**Figure S28**: Standard curve generated from LNCaP cells stained with Calcein AM.

### 6 Uncropped Images of Western Blots and Fluorescent Gels

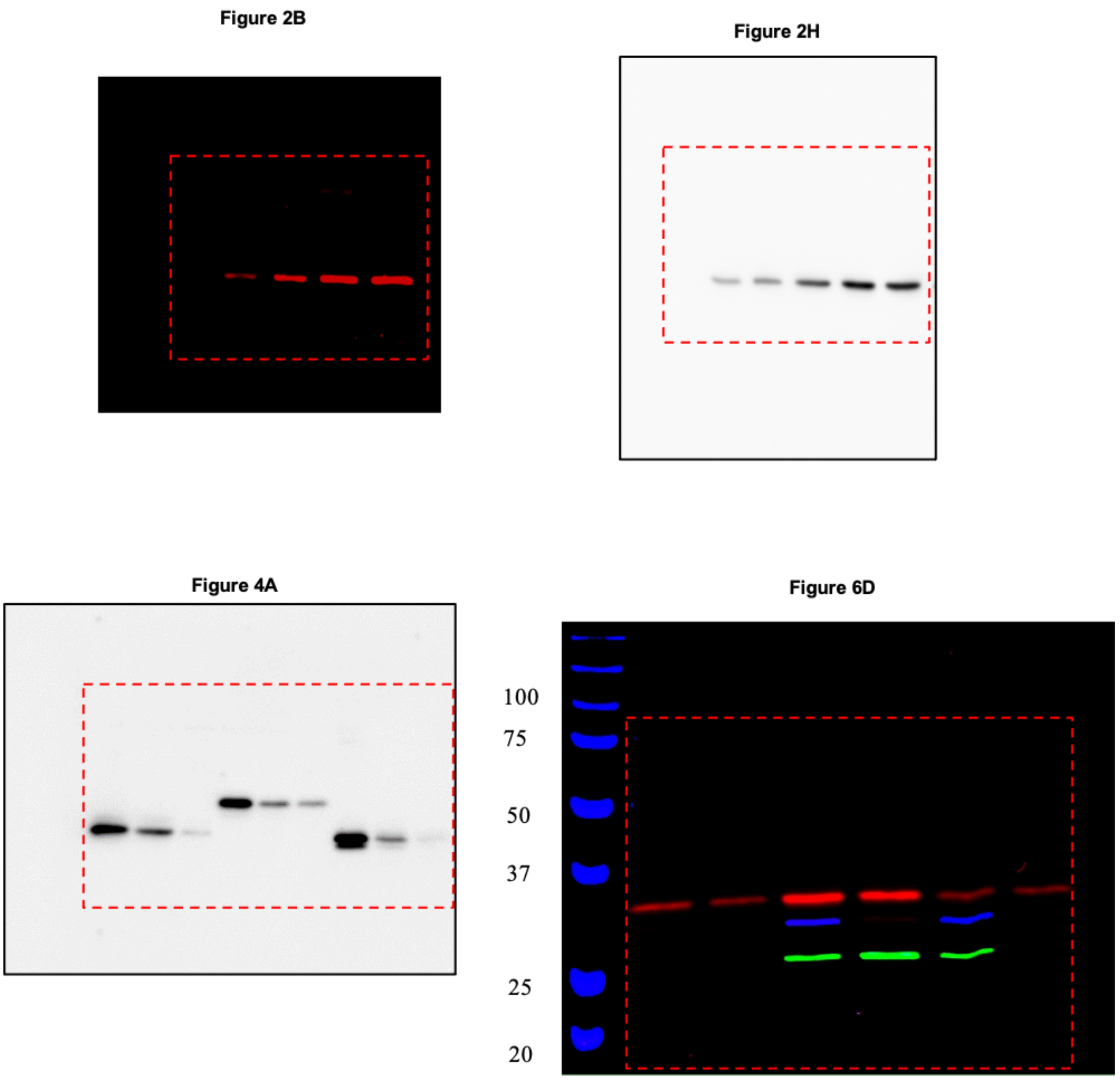

**Figure S29**: Uncropped images from fluorescent gels and streptavidin blots demonstrating probe specificities.

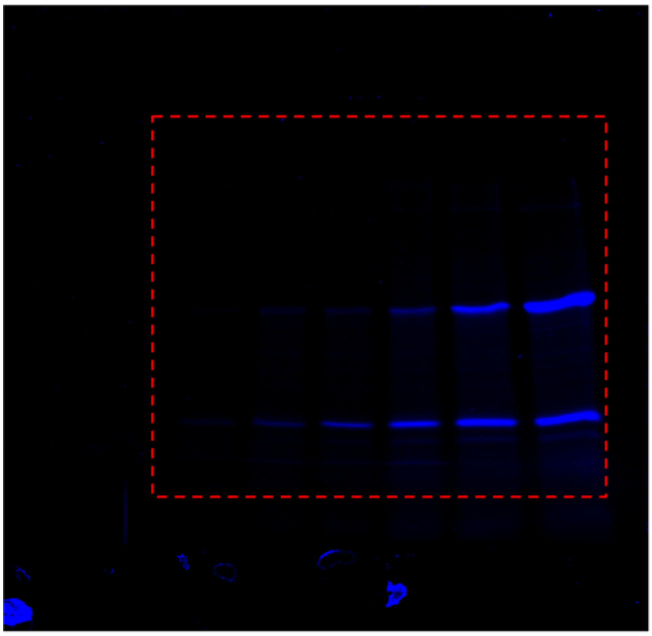

**Figure S19 middle**

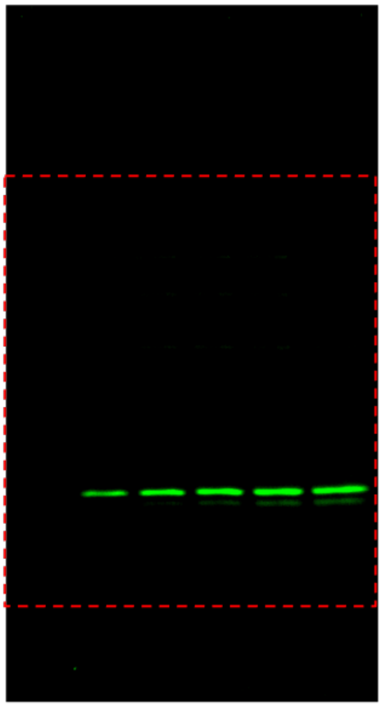

**Figure S19 bottom**

**Figure S30**: Uncropped images from fluorescent gels and streptavidin blots demonstrating probe specificities.

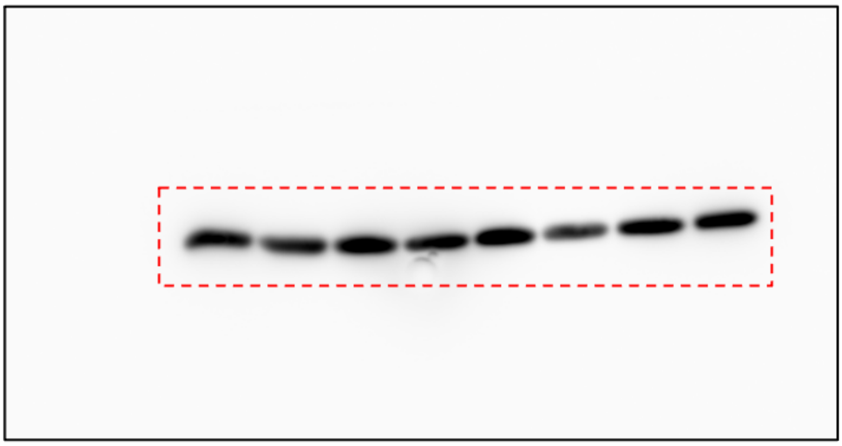

Figure 2H – Total KLK3

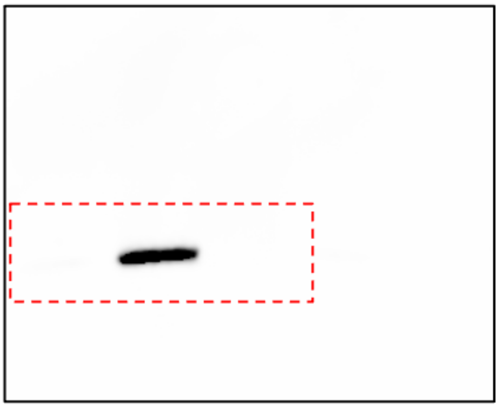

Figure 4B – Active KLK2

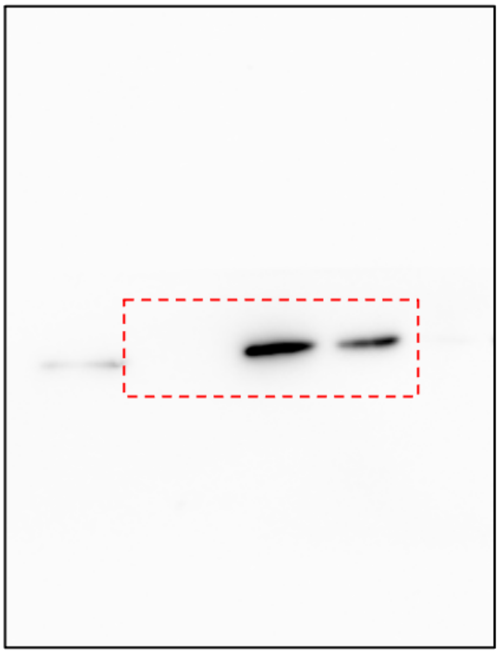

Figure 4B –Total KLK2

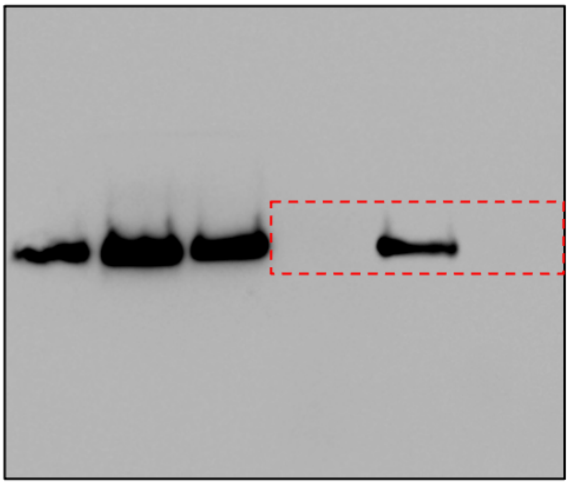

Figure 4B – Active KLK3

**Figure S31**: Uncropped western blots

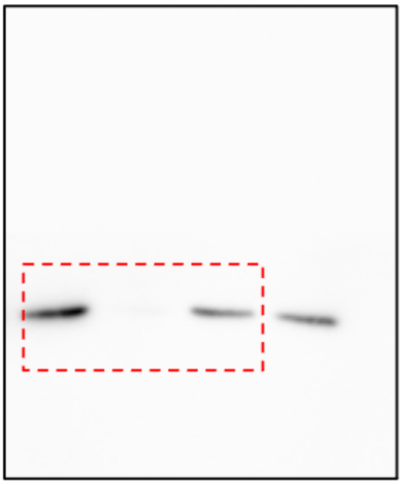

Figure 4B – Active KLK14

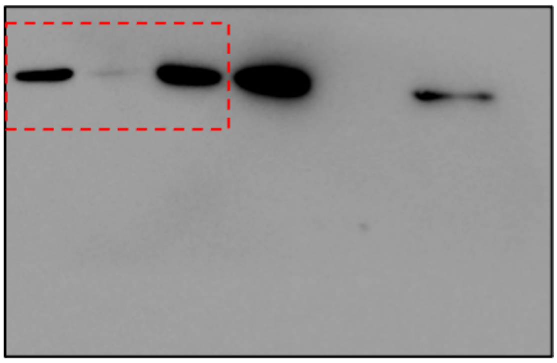

Figure 4B – Total KLK14

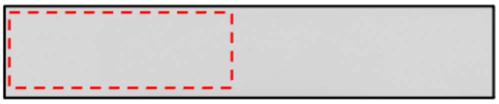

Figure 4B –Active Transferrin

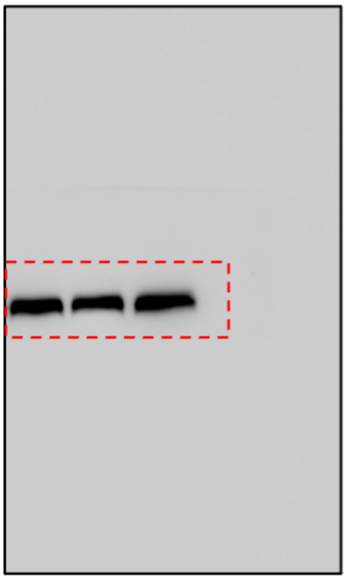

Figure 4B – Total Transferrin

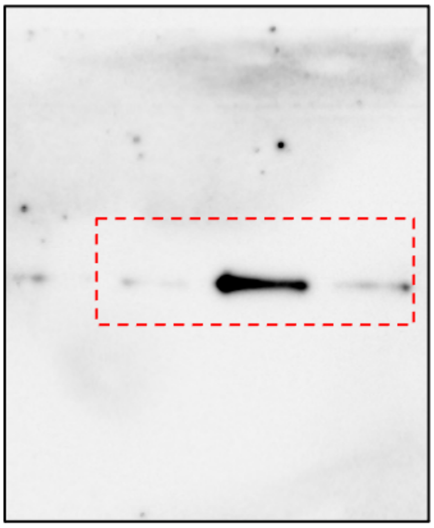

Figure 4B – Total KLK3

**Figure S32**: Uncropped western blots

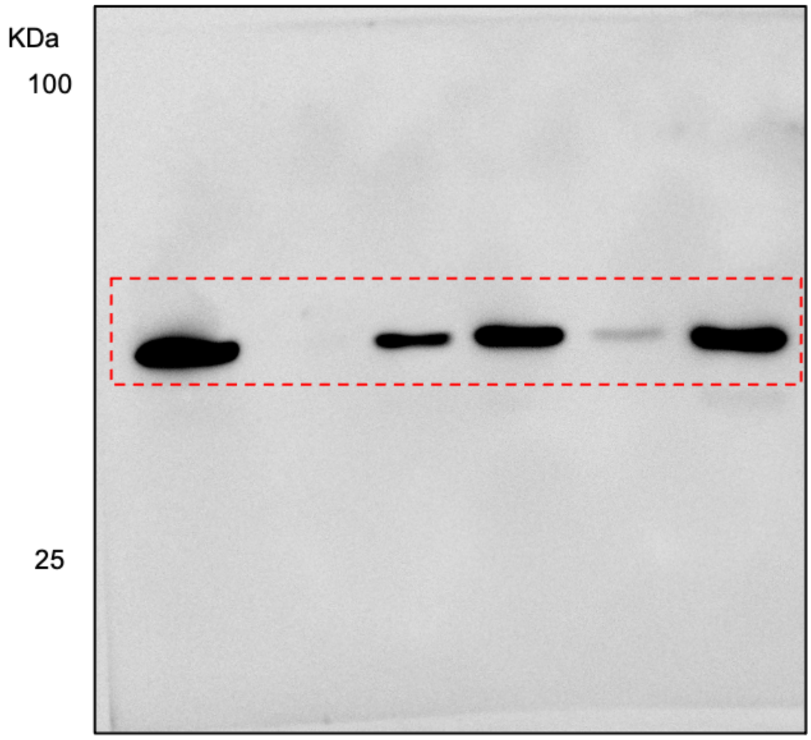

**Figure 4C – Total KLK3**

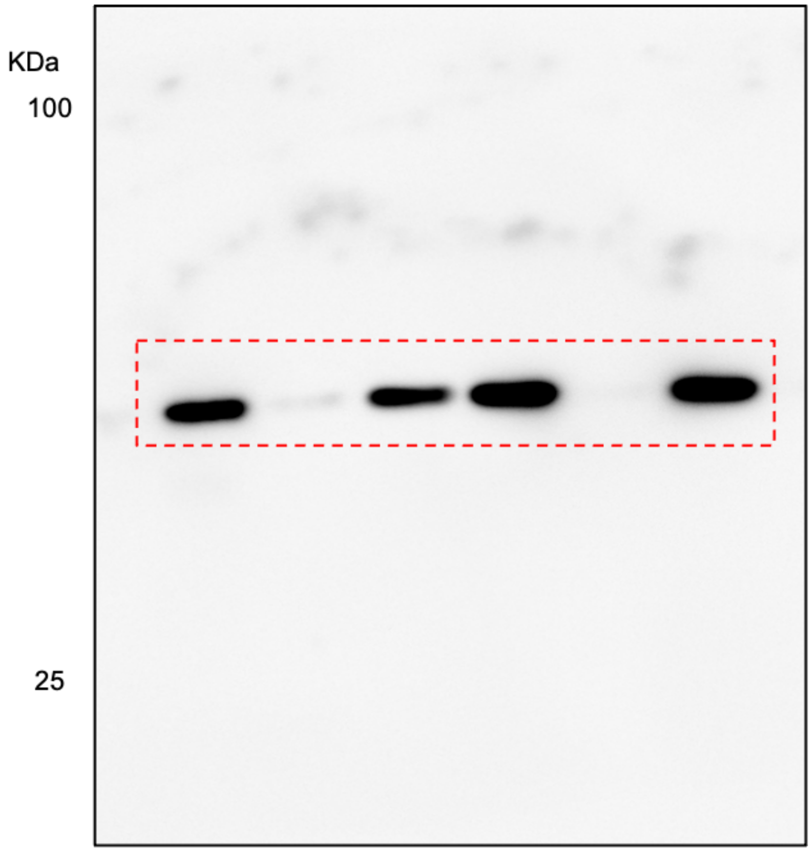

**Figure 4C – Active KLK3**

**Figure S33**: Uncropped western blots

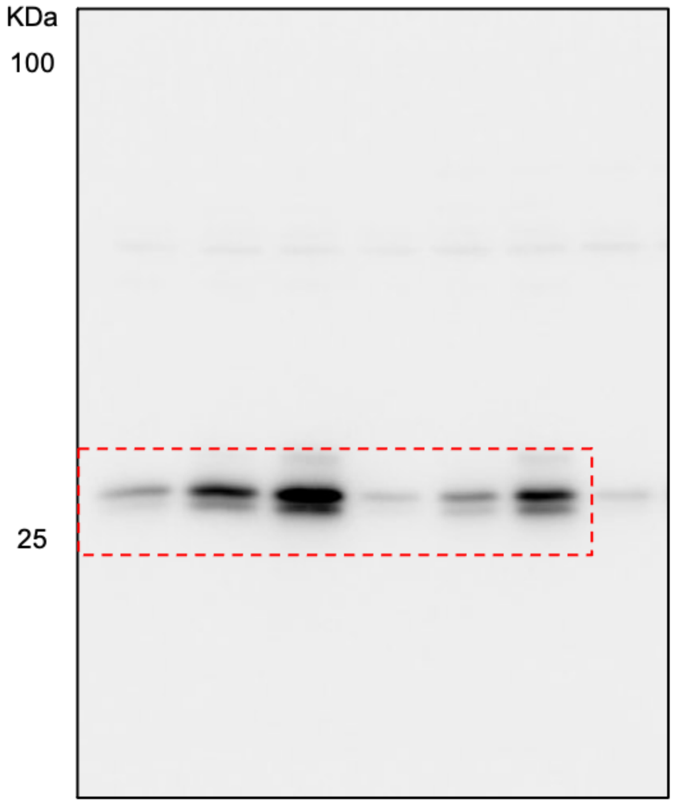

**Figure 4C – Active KLK14**

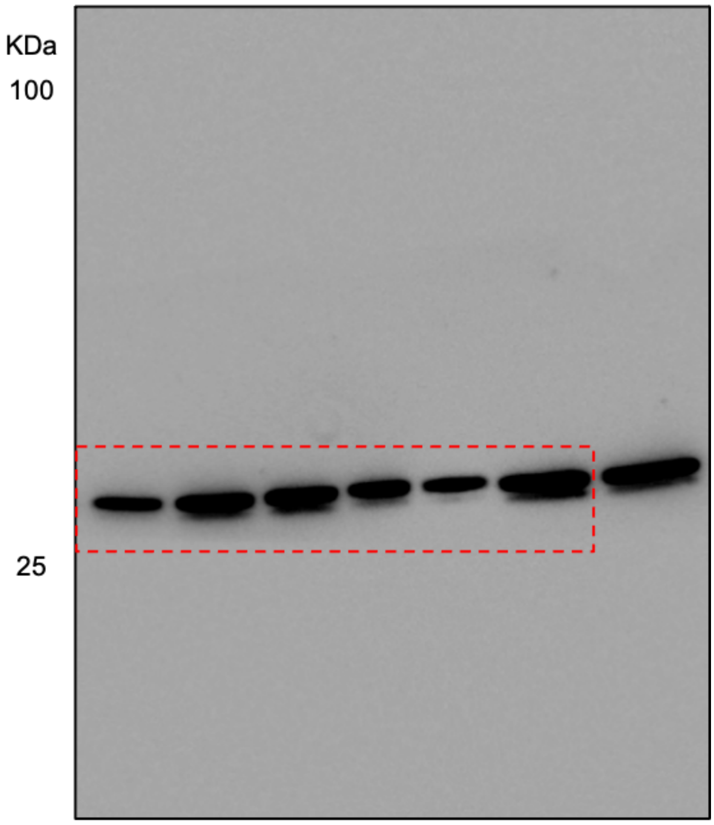

**Figure 4C – Total KLK14**

**Figure S34**: Uncropped western blots

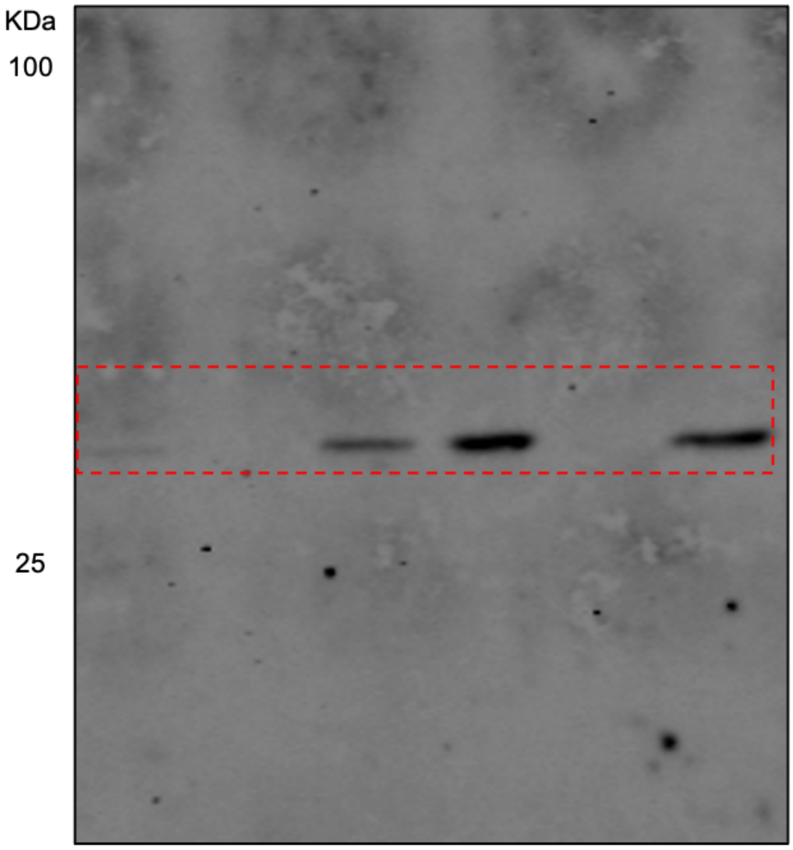

**Figure 4C – Active KLK2**

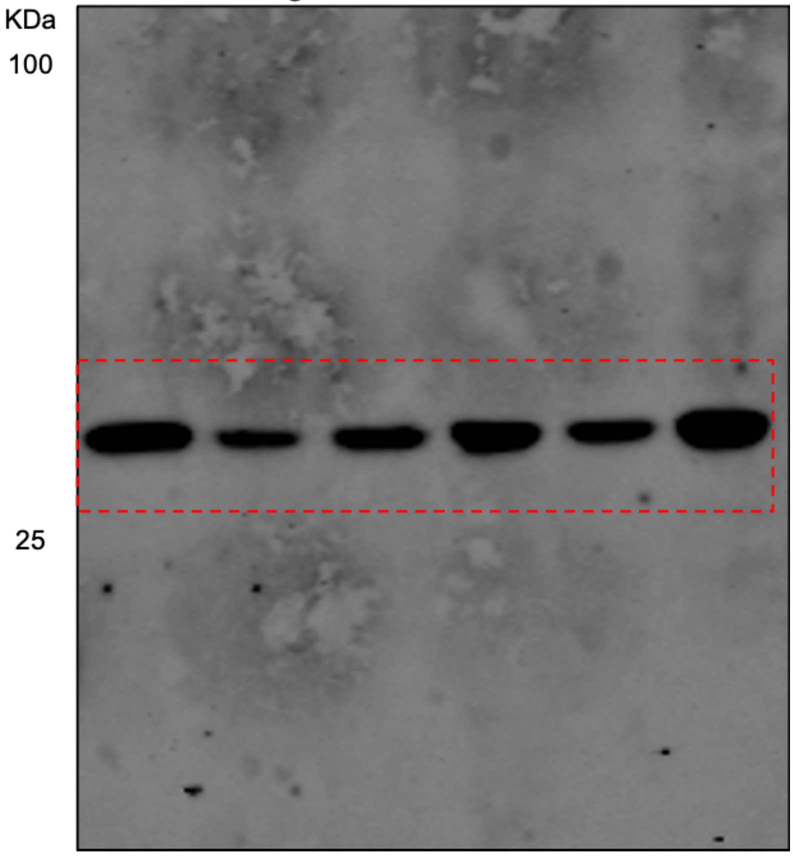

**Figure 4C – Total KLK2**

**Figure S35**: Uncropped western blots

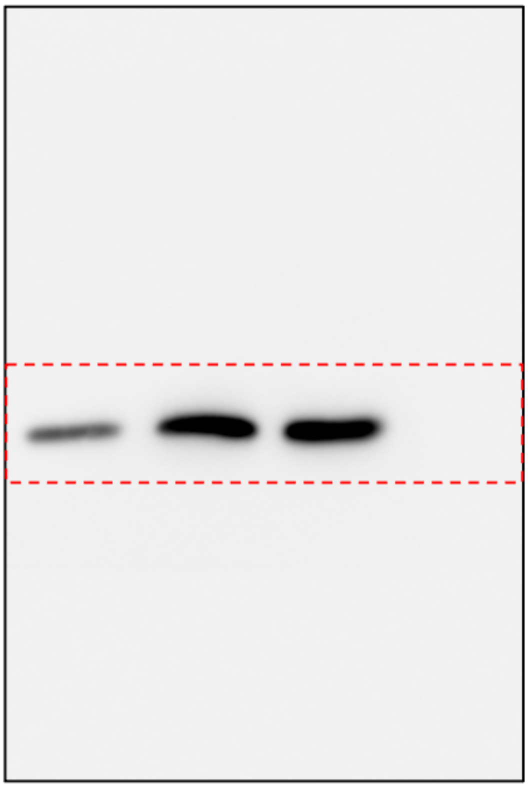

**Figure 5B – Total KLK2**

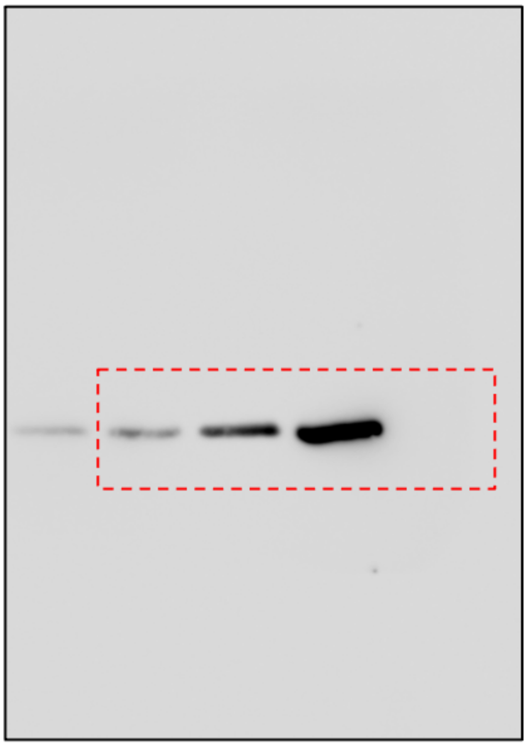

**Figure 5B – Active KLK2**

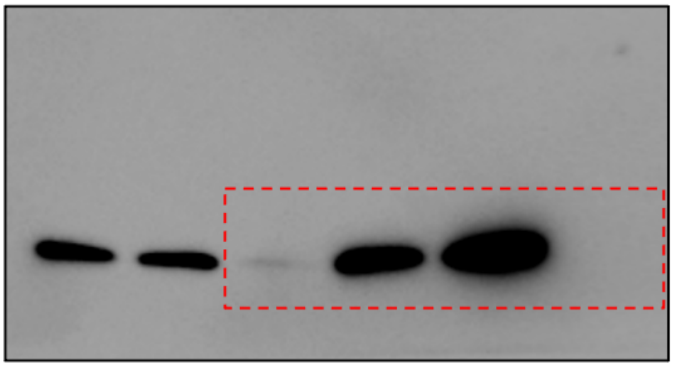

**Figure 5B – Total KLK3**

**Figure S36**: Uncropped western blots

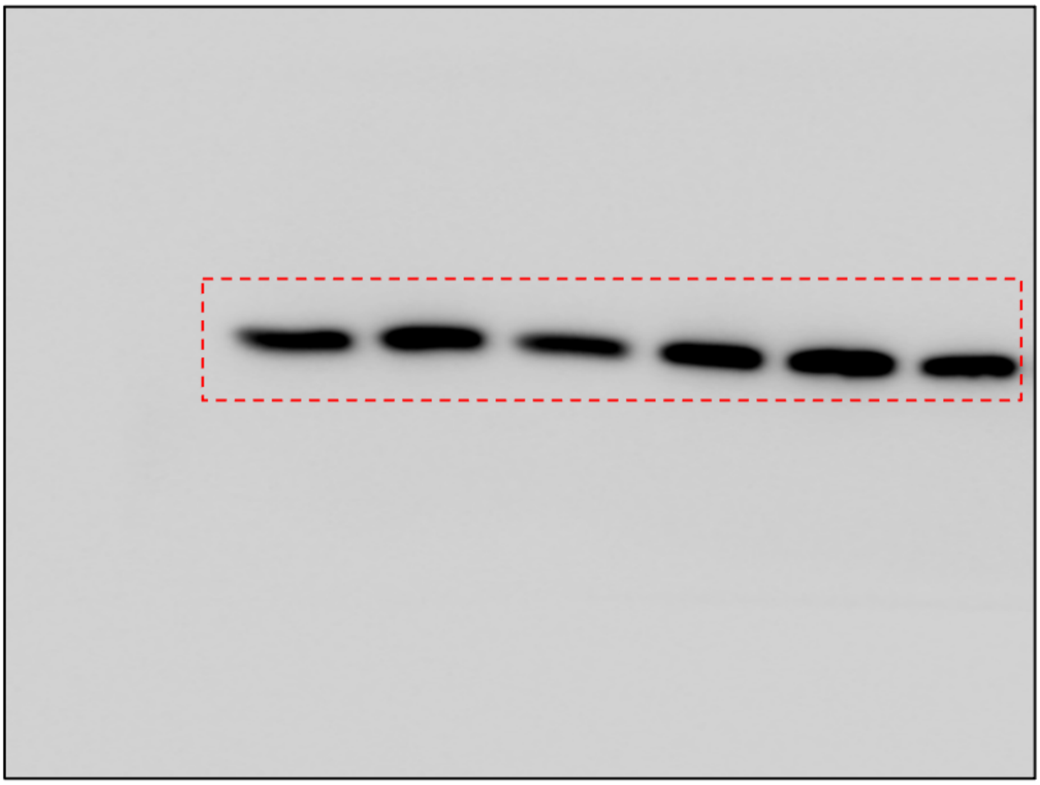

**Figure 6D – Total KLK3**

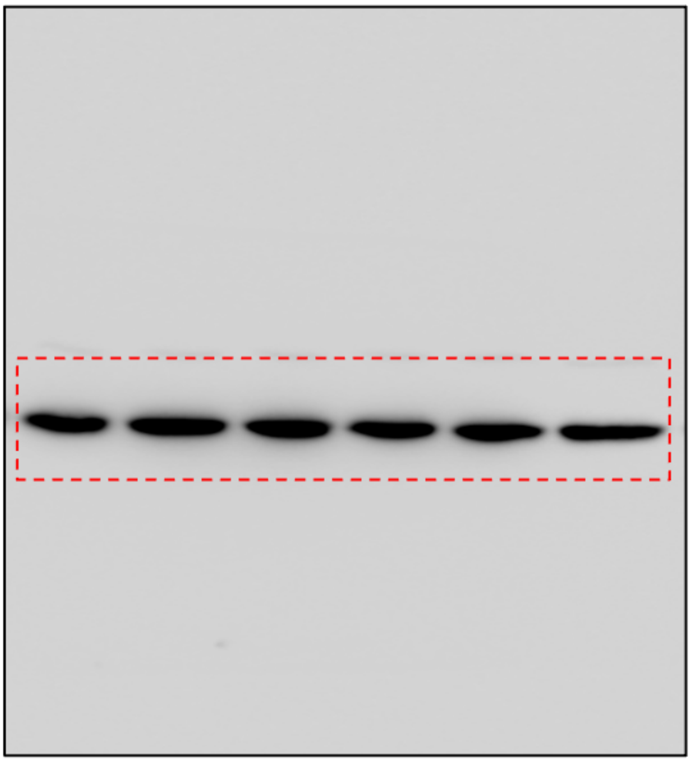

**Figure 6D – Total KLK2**

**Figure S37**: Uncropped western blots

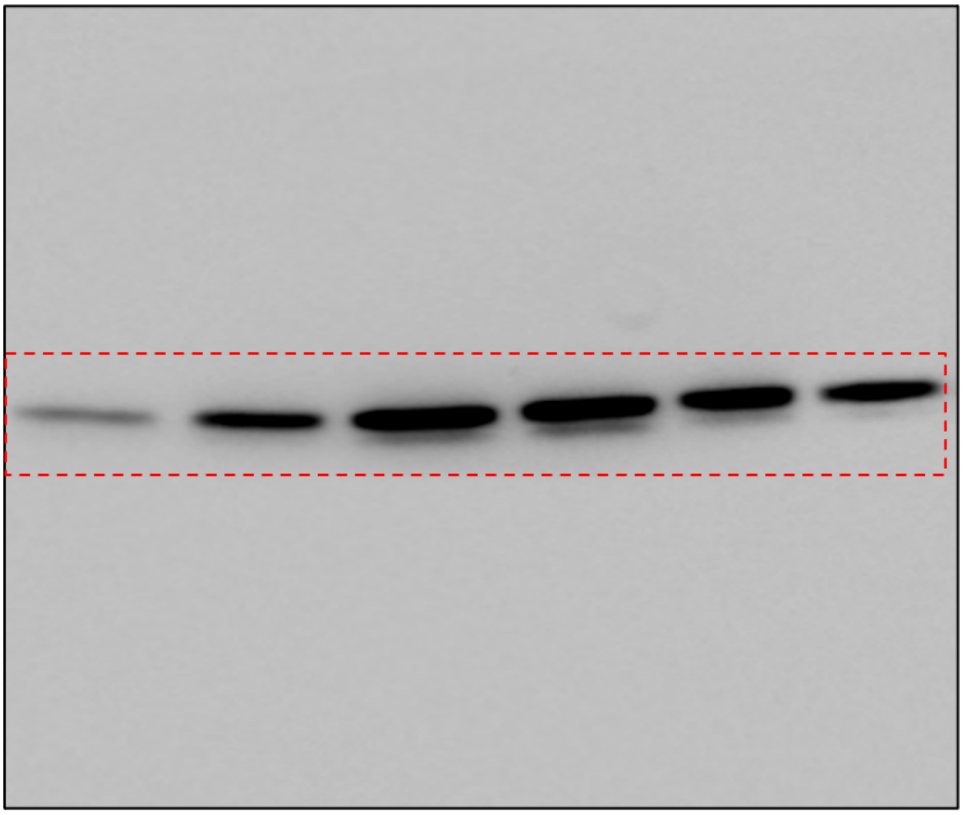

Figure 6D – Total KLK14

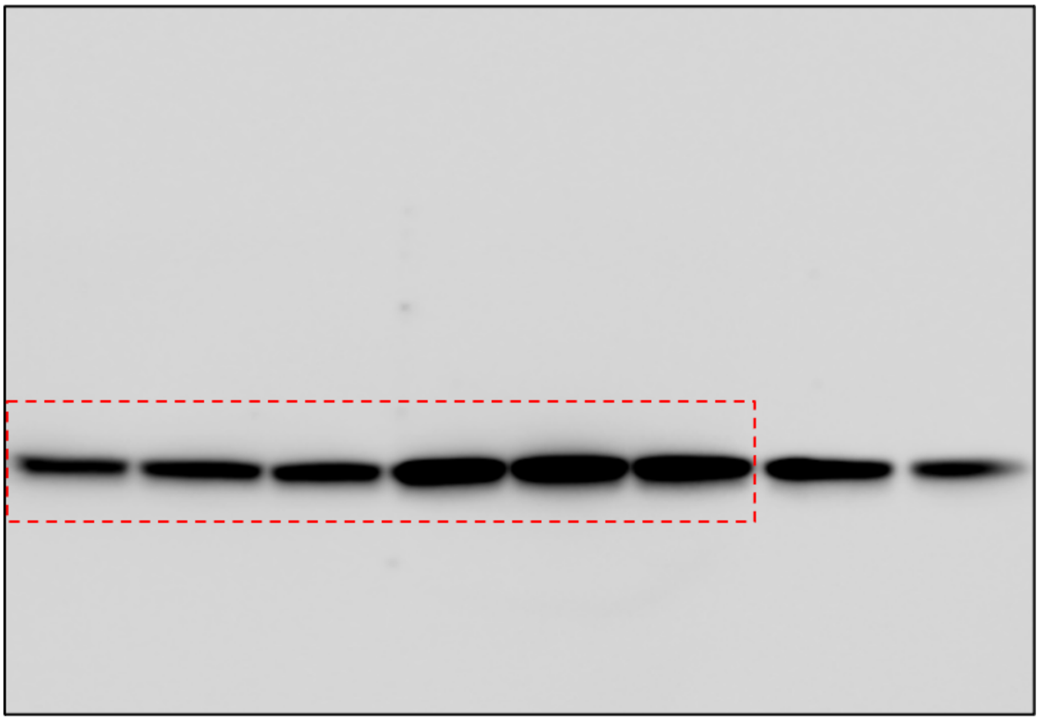

Figure 6D – Total Transferrin

**Figure S38**: Uncropped western blots

### 7 Characterization Data

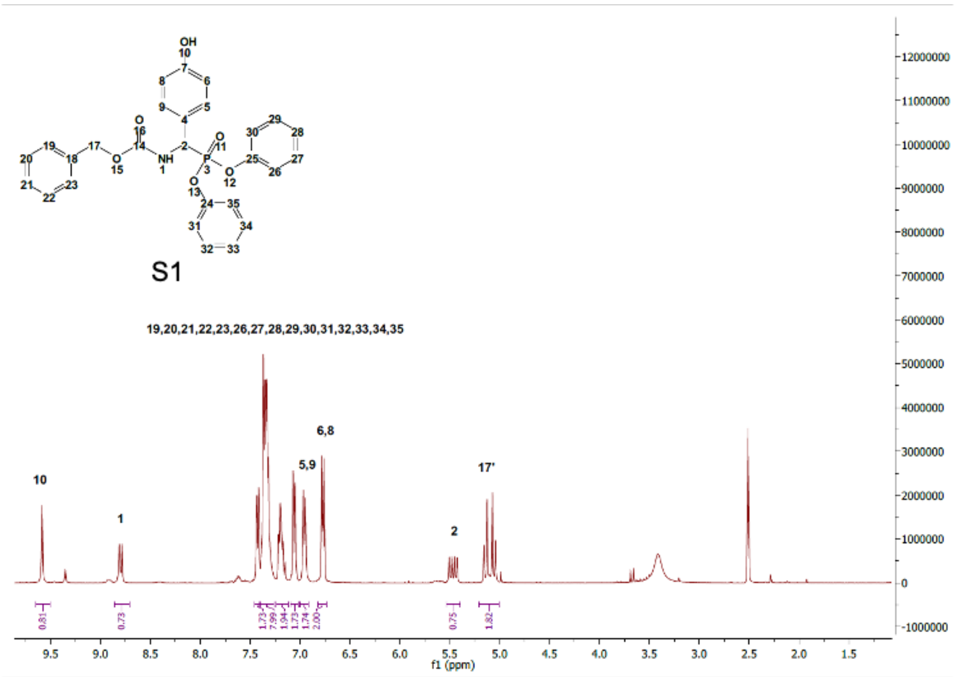

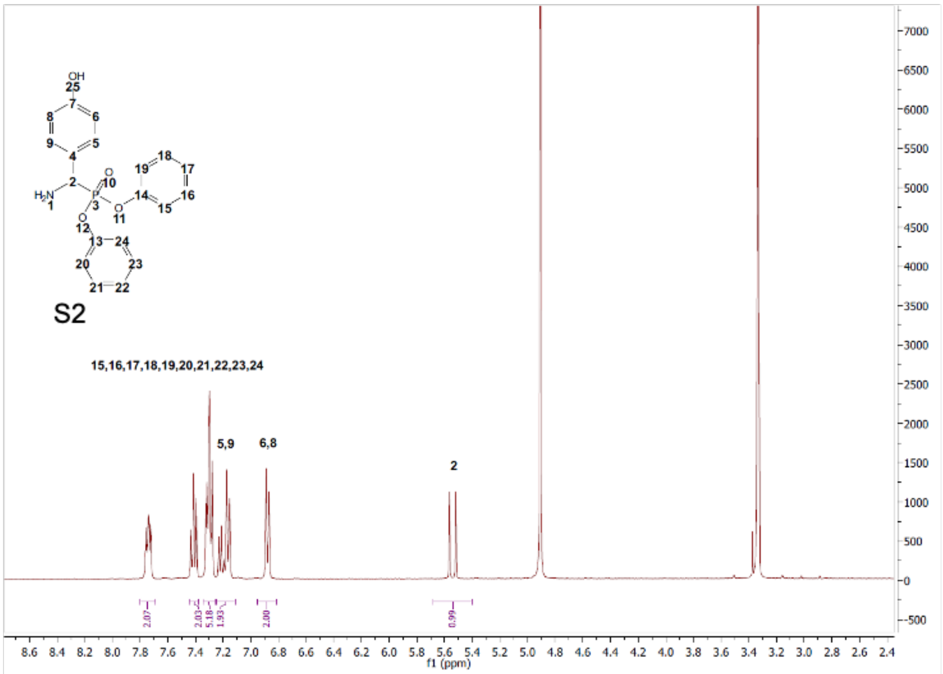

**Figure S39**: $^{1}$H NMR of S1 and S2

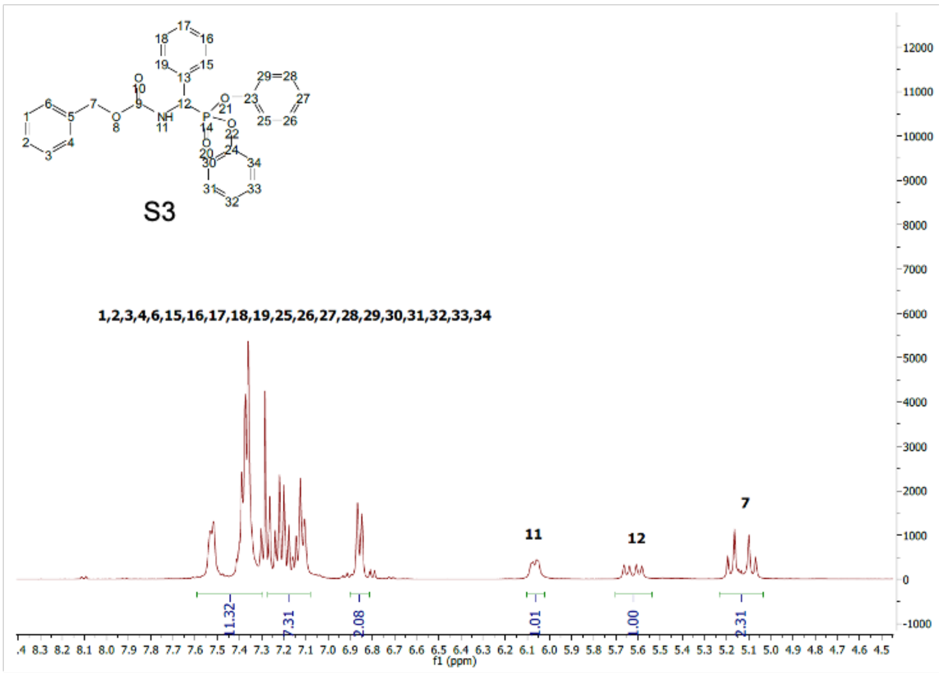

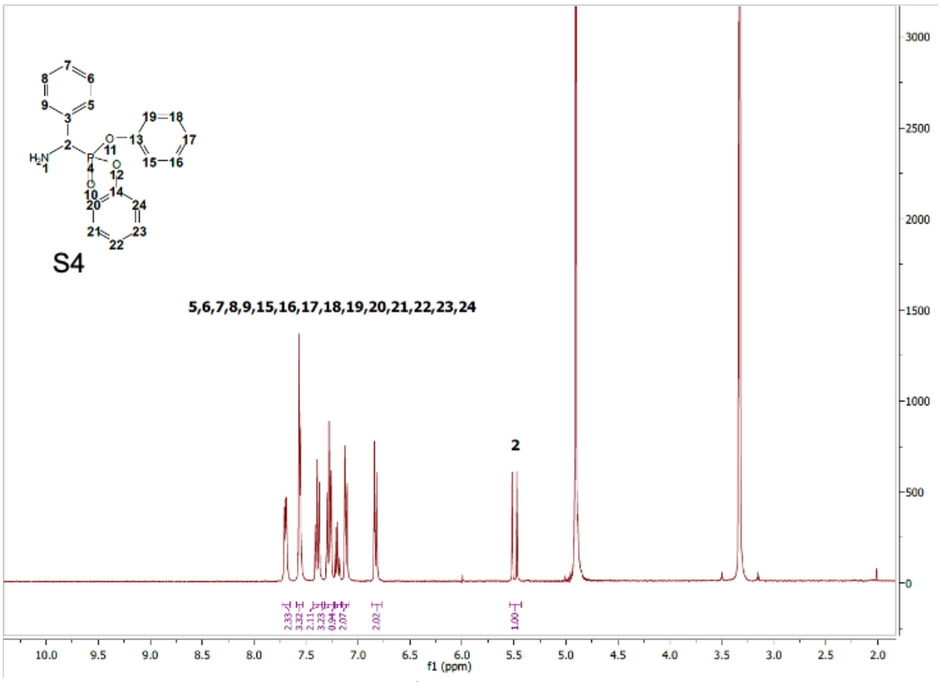

**Figure S40**: $^{1}$H NMR of S3 and S4

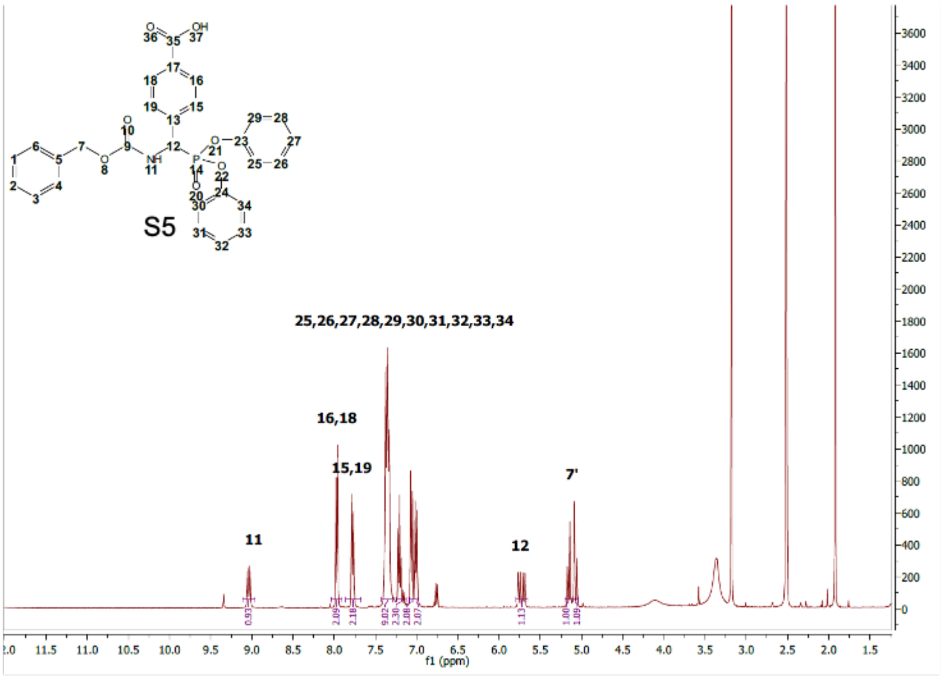

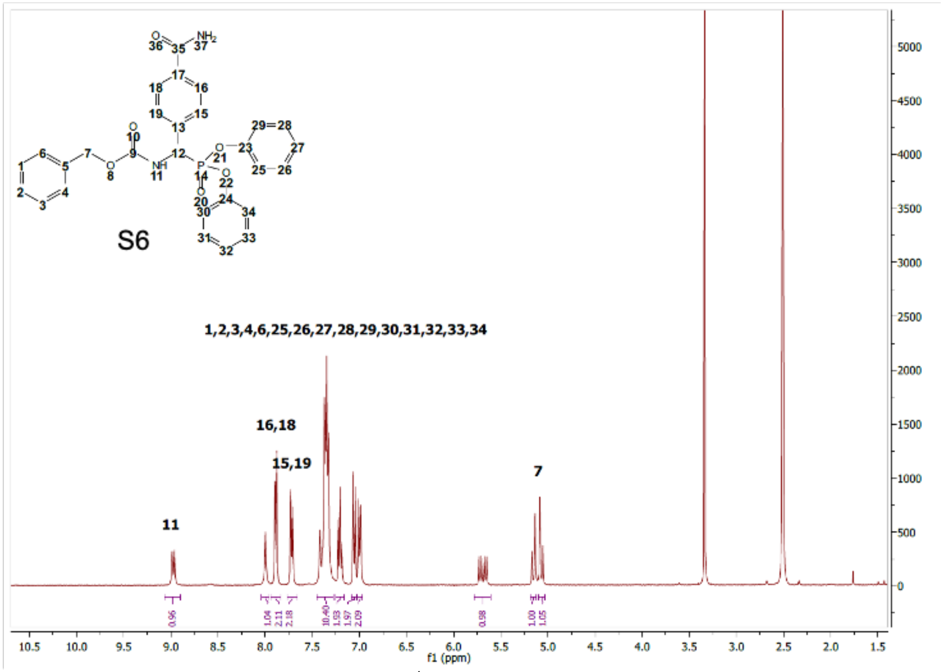

**Figure S41**: $^1$H NMR of S5 and S6

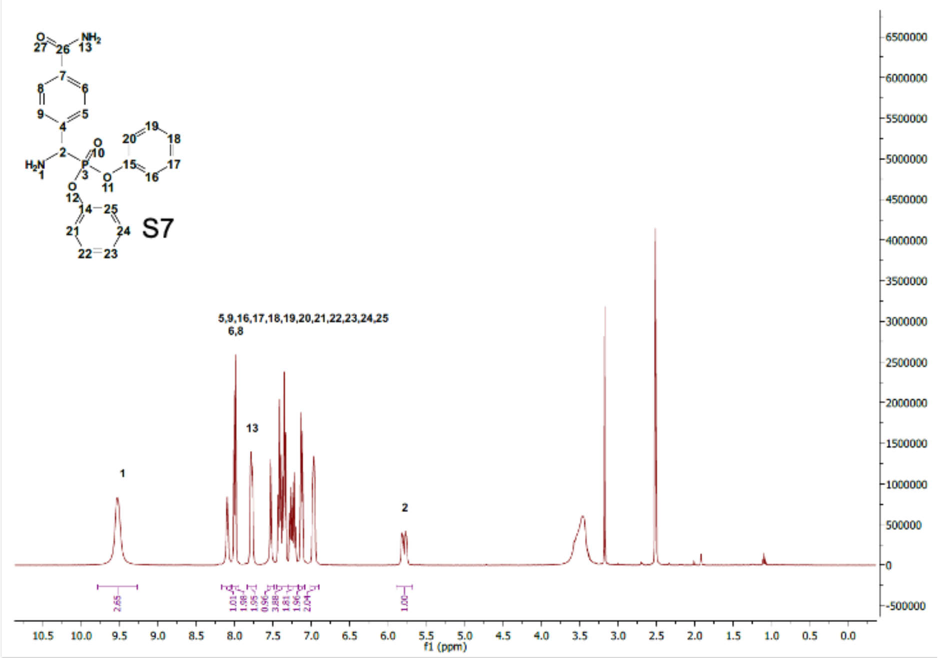

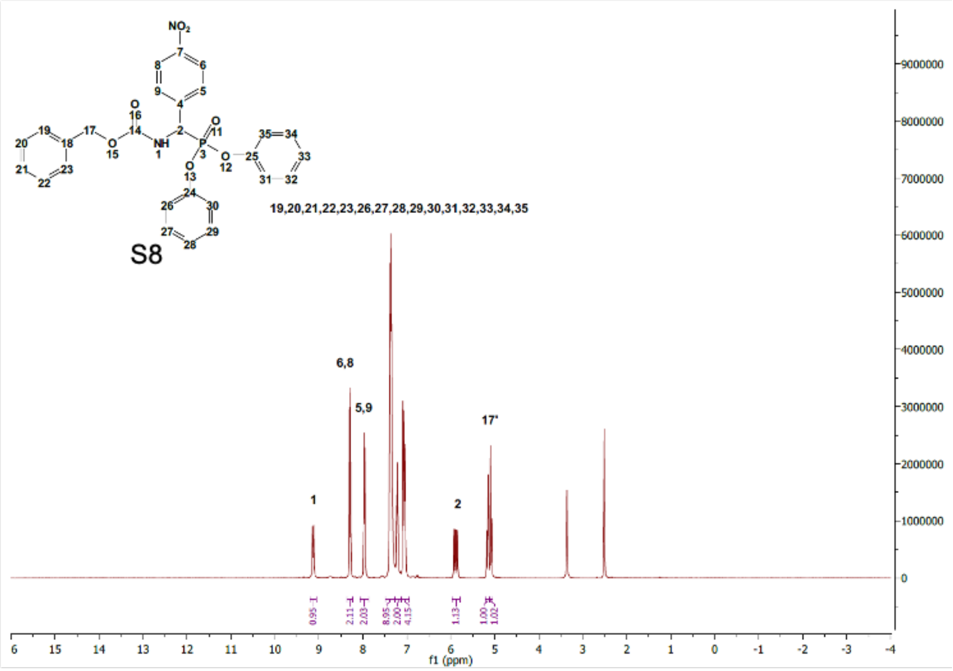

**Figure S42**: $^{1}H$ NMR of S7 and S8

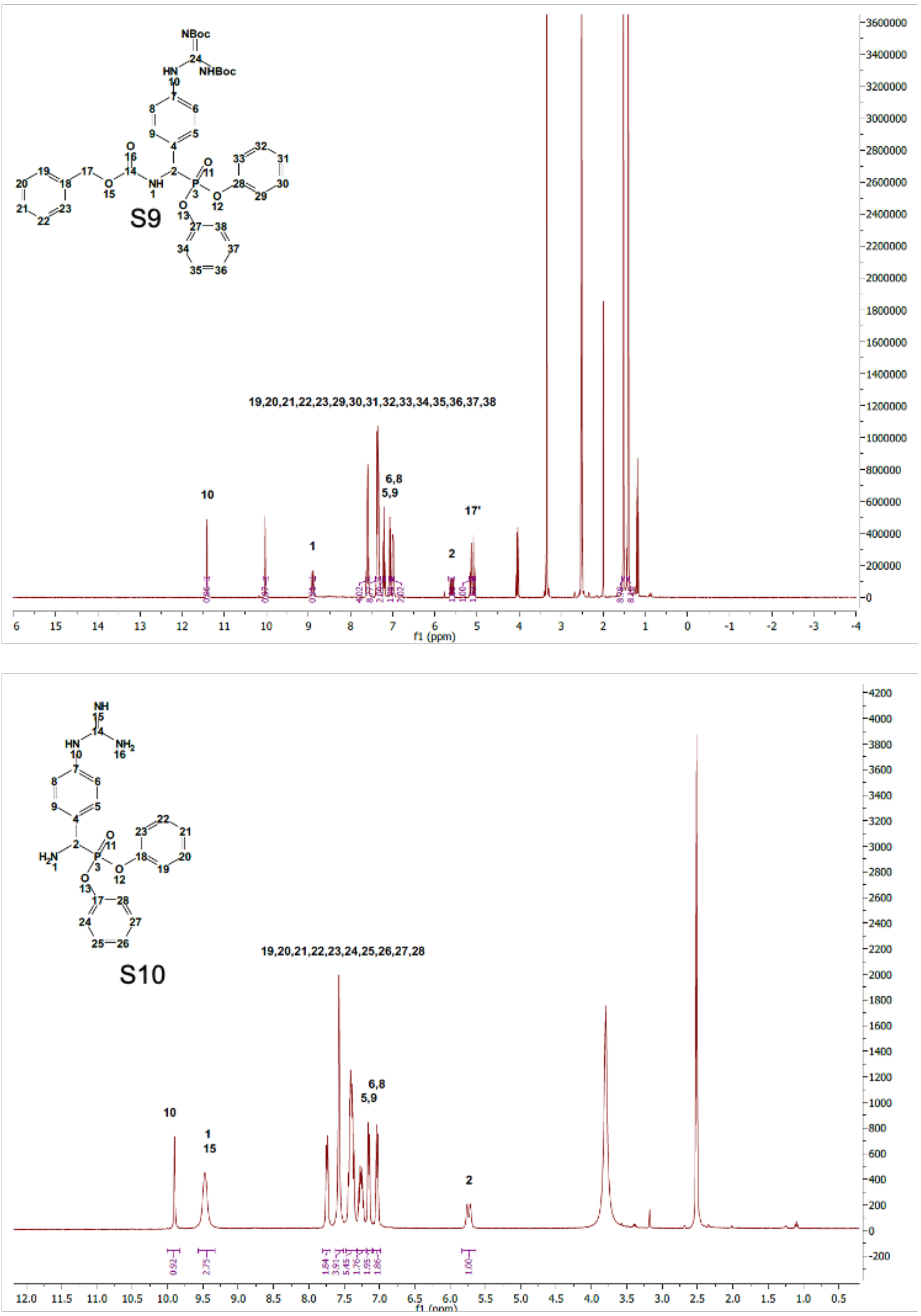

**Figure S43**: $^1$H NMR of S9 and S10

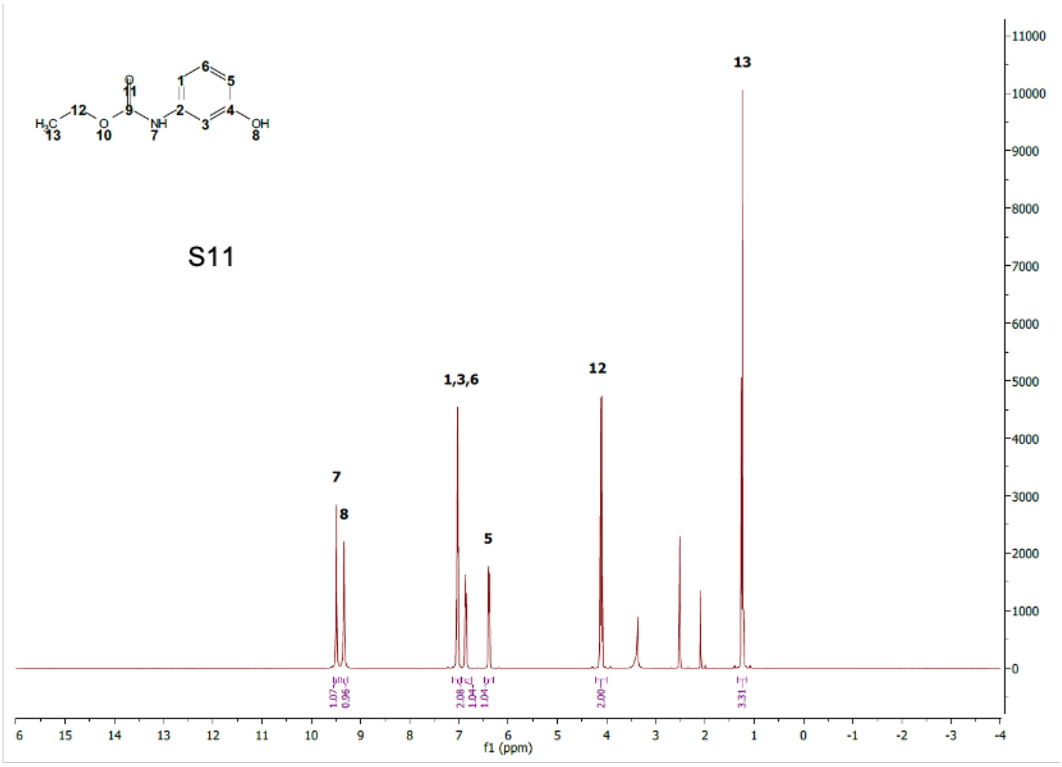

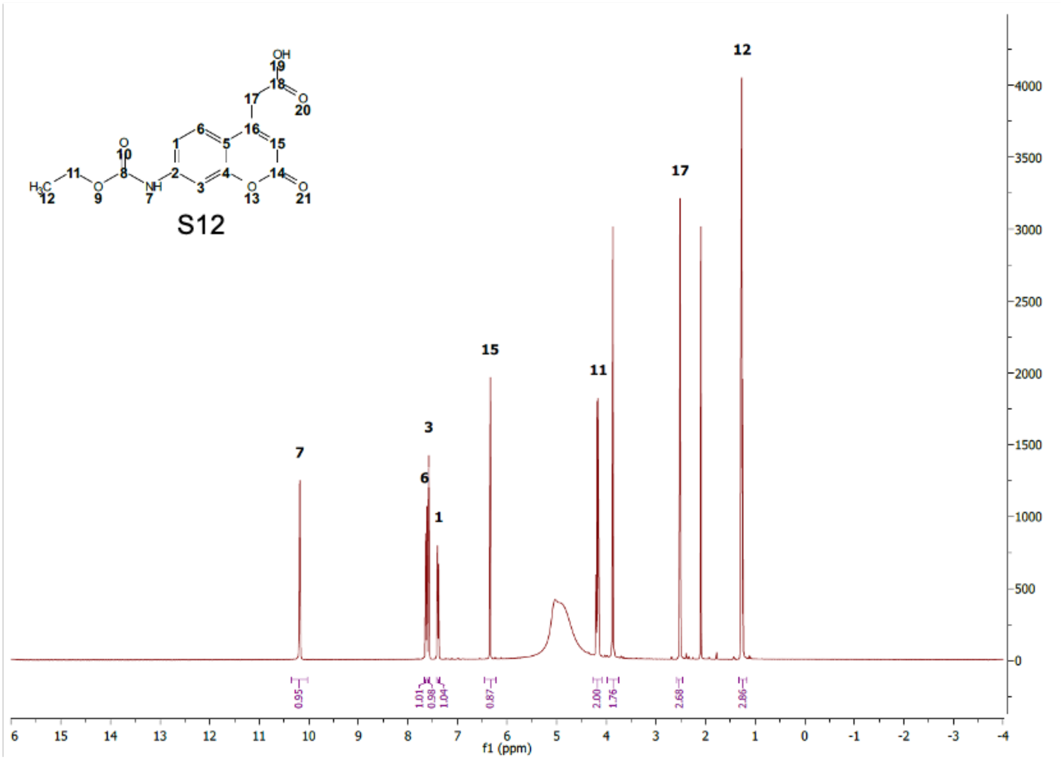

**Figure S44**: $^{1}$H NMR of S11 and S12

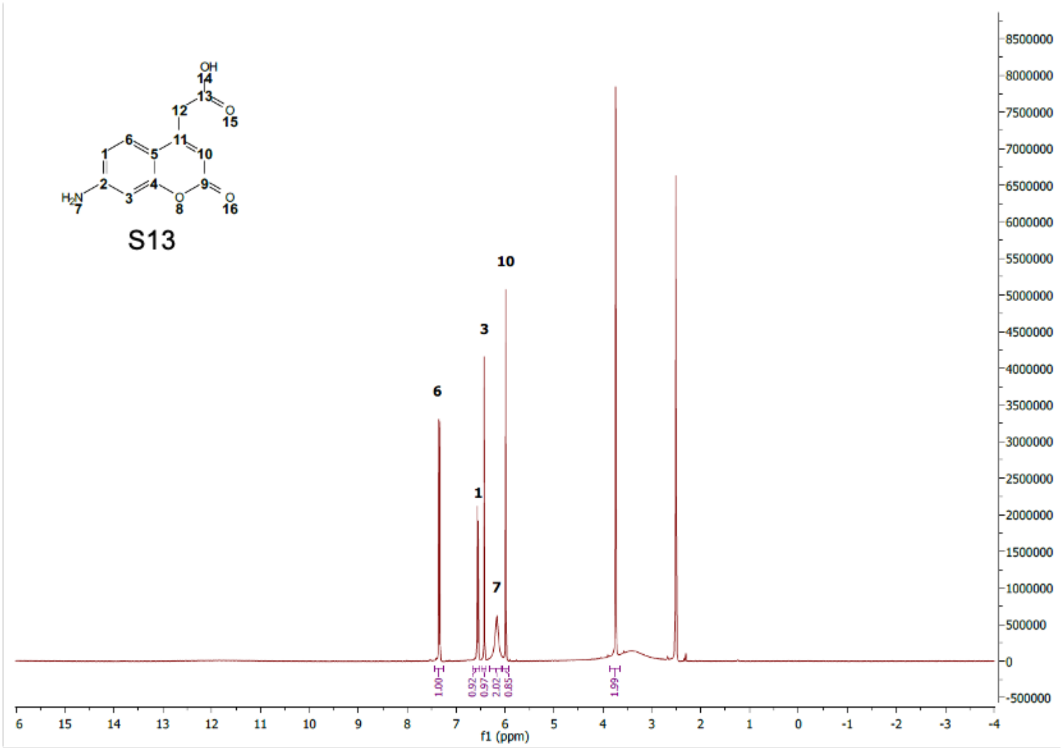

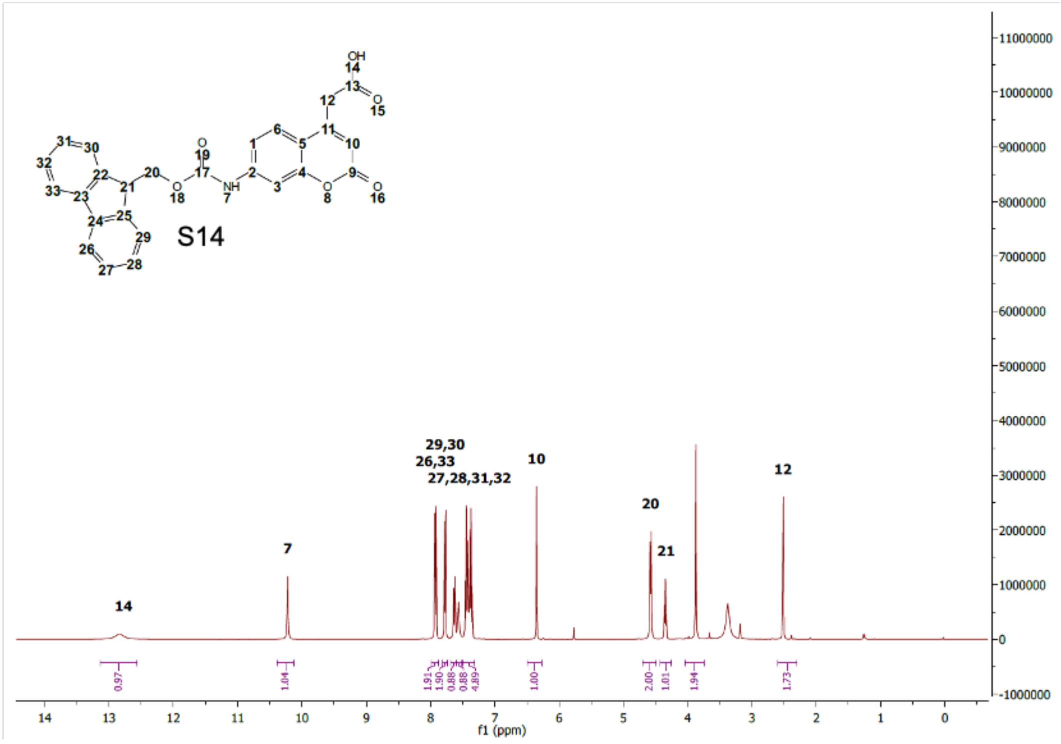

**Figure S45**: $^{1}$H NMR of S13 and S14

HPLC

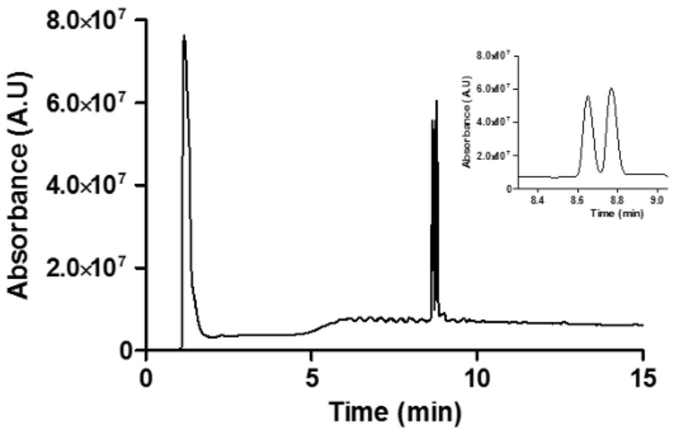

HRMS

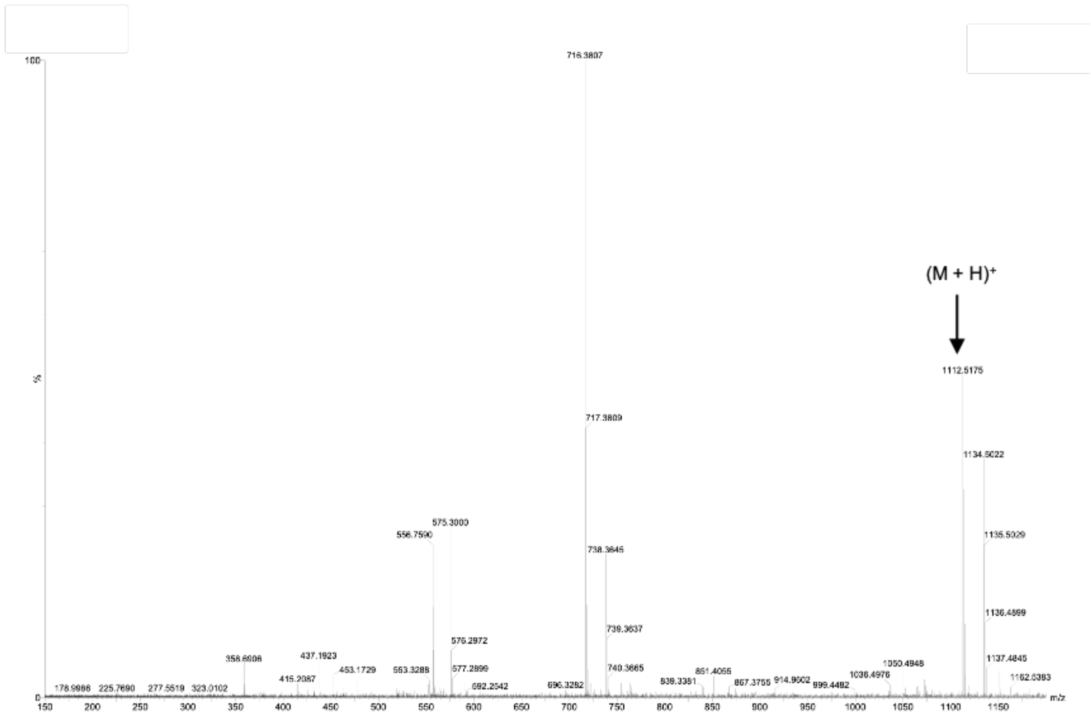

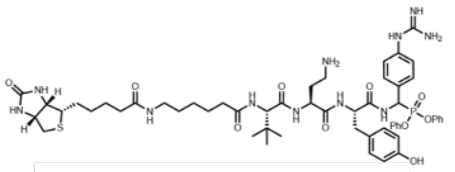

KLK2 Optimized ABP - **31** (KLK2_bABP)

**Figure S46**: HPLC (20-98% acetonitrile gradient) and HRMS data for KLK2_bABP

HPLC

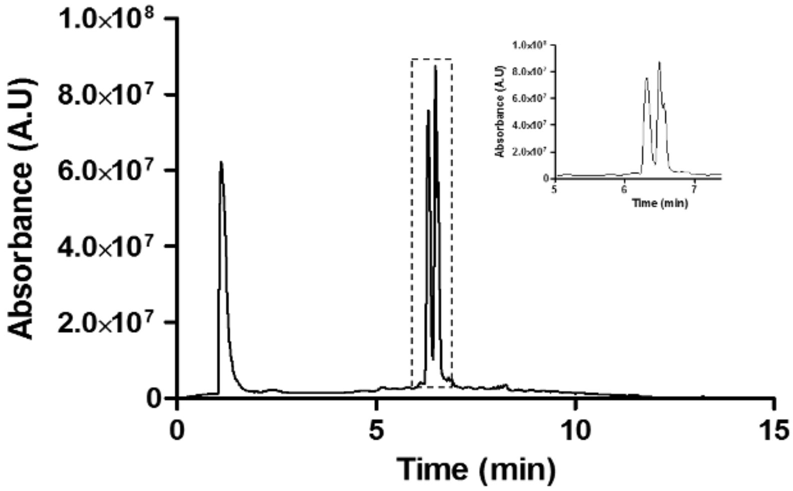

HRMS

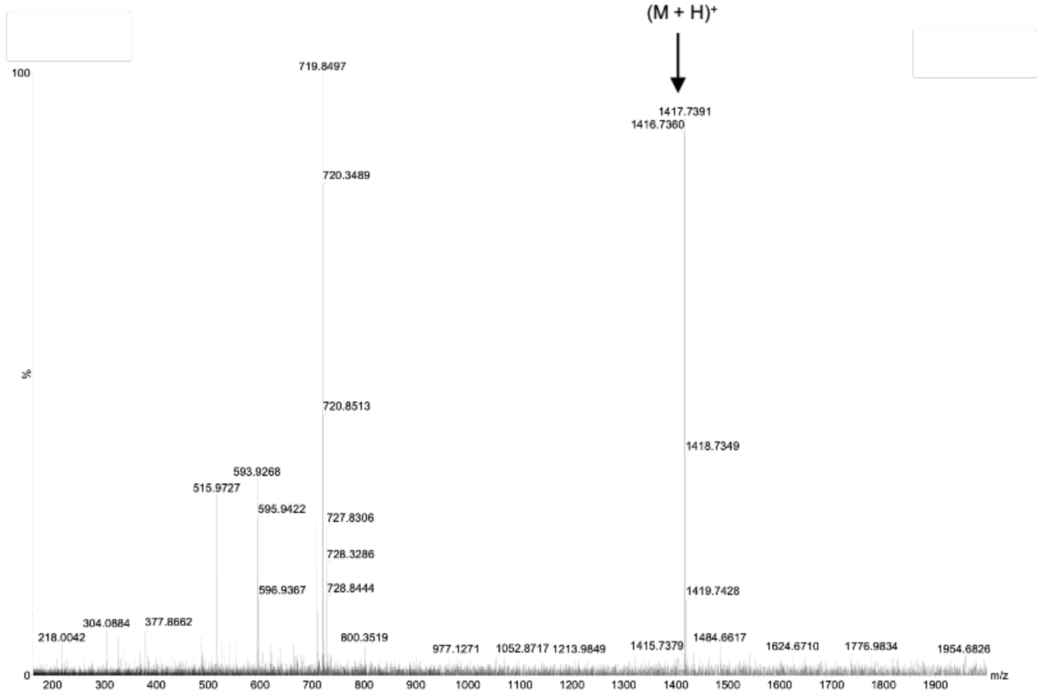

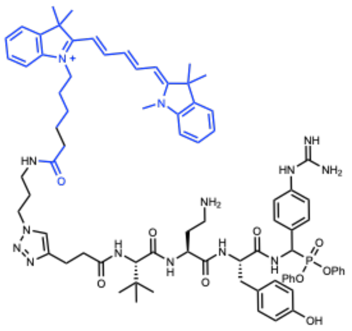

KLK2 Optimized ABP - **42** (KLK2_fABP)

**Figure S47**: HPLC (20-98% acetonitrile gradient) and HRMS data for KLK2_fABP

HPLC

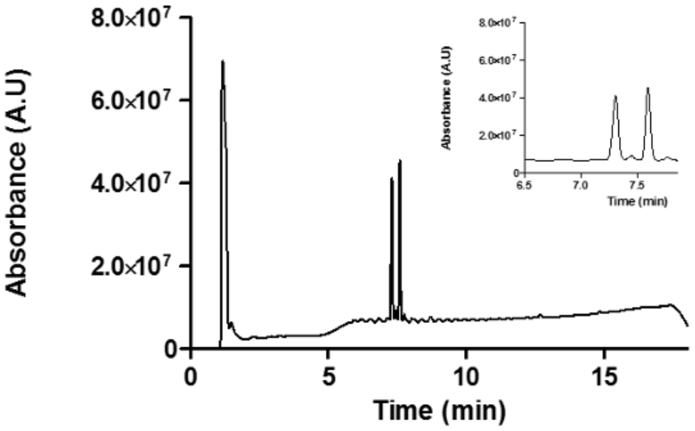

HRMS

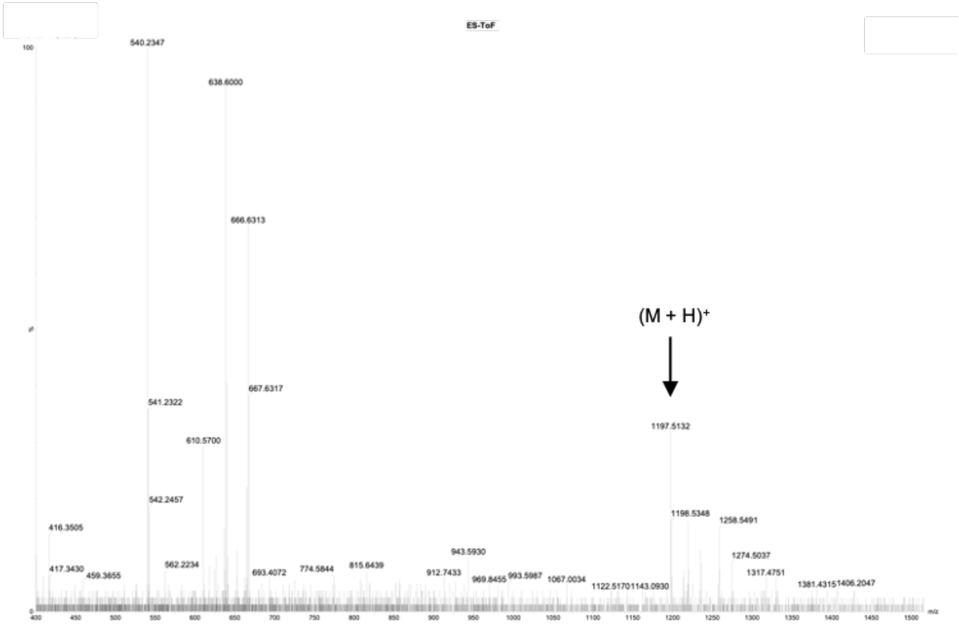

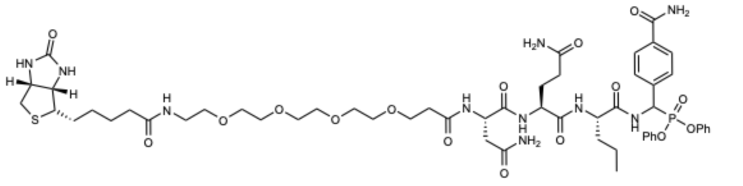

KLK3 Optimized ABP - **20** (KLK3_bABP)

**Figure S48**: HPLC (20-98% acetonitrile gradient) and HRMS data for KLK3_bABP

HPLC

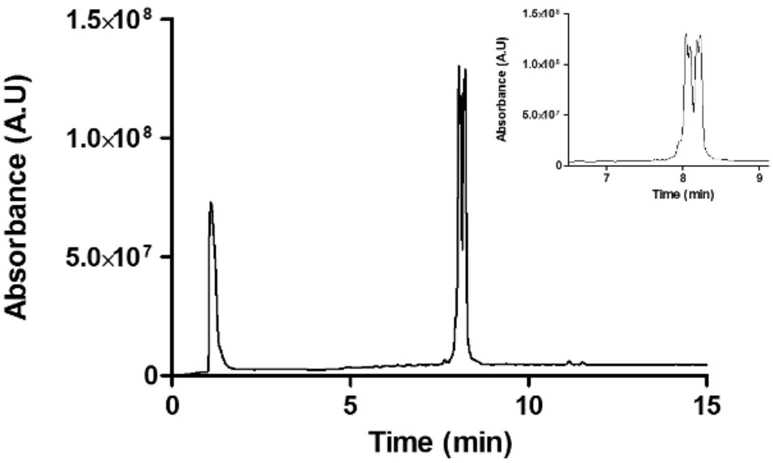

HRMS

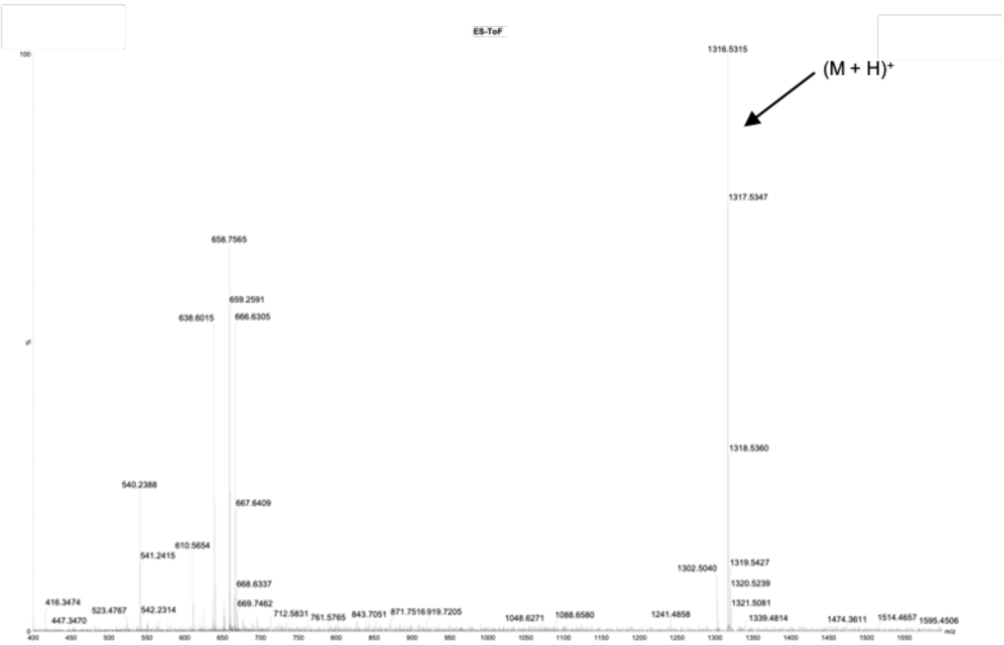

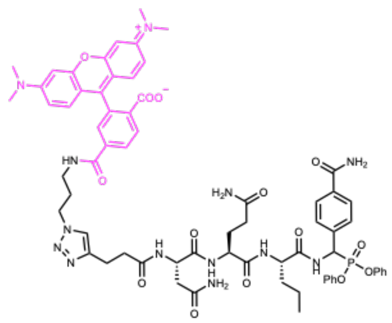

KLK3 Optimized ABP - **20** (KLK3_fABP)

**Figure S49**: HPLC (20-98% acetonitrile gradient) and HRMS data for KLK3_fABP

HPLC

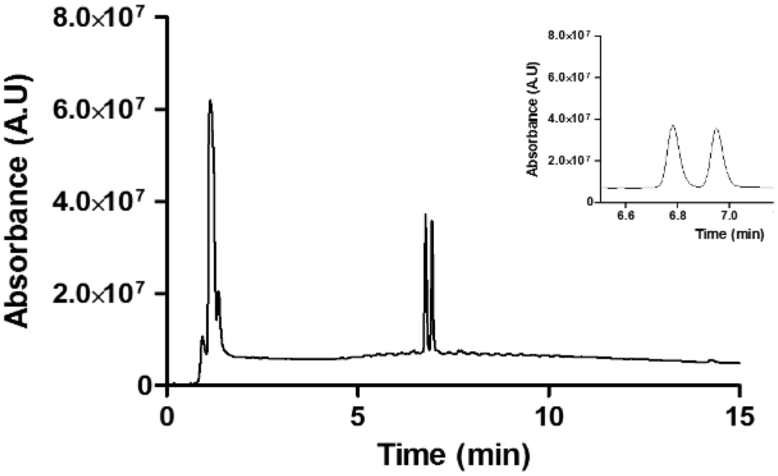

HRMS

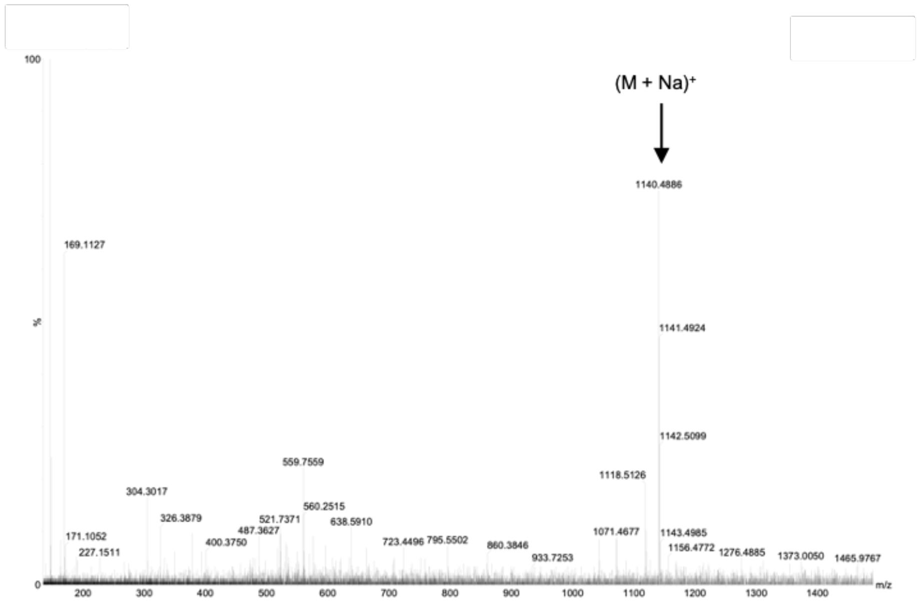

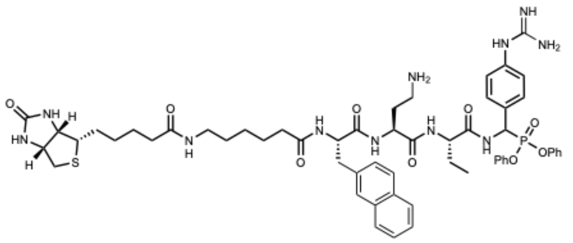

KLK14 Optimized ABP - **39** (KLK14_bABP)

**Figure S50**: HPLC (20-98% acetonitrile gradient) and HRMS data for KLK14_bABP

HPLC

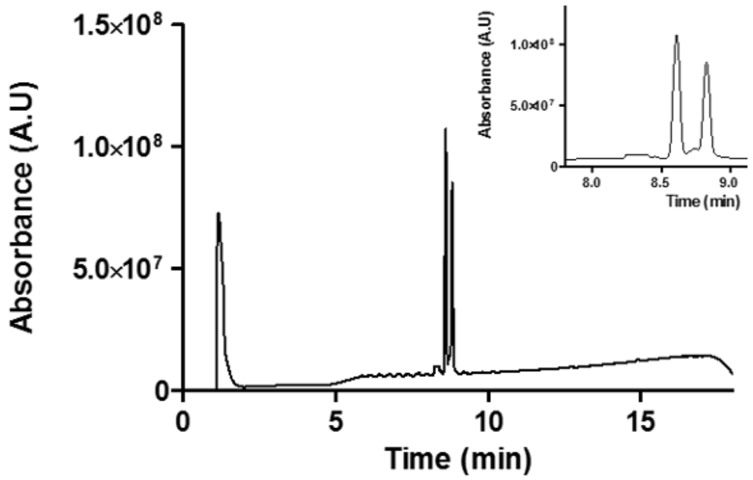

HRMS

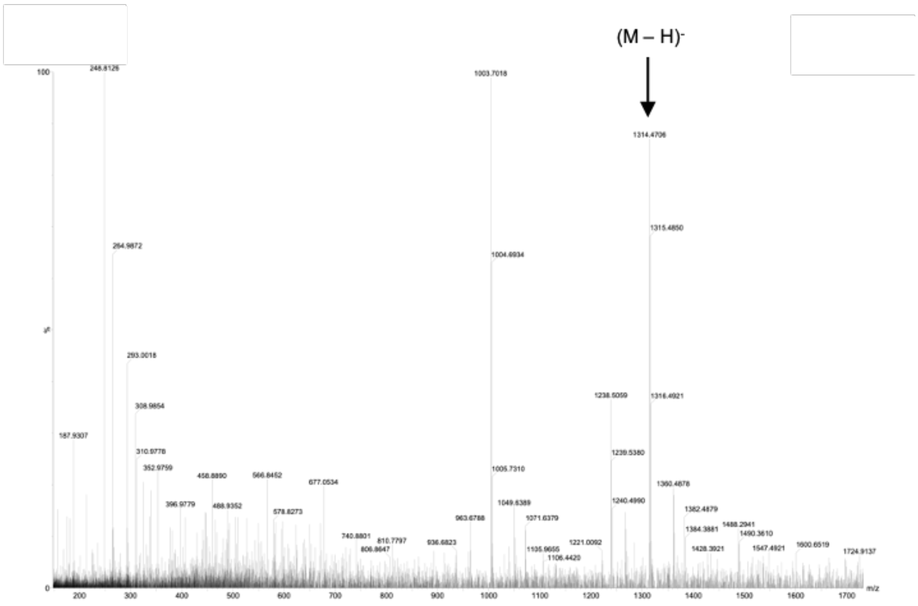

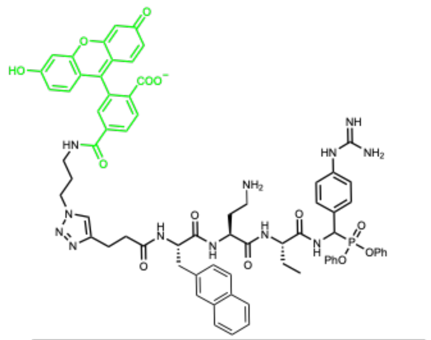

KLK14 Optimized ABP - **43** (KLK14_fABP)

**Figure S51**: HPLC (20-98% acetonitrile gradient) and HRMS data for KLK14_fABP
